# O-GlcNAcylation and an importin-β radial gradient keep the FG barrier liquid in live-cell nuclear pores

**DOI:** 10.64898/2025.12.09.693204

**Authors:** Miao Yu, Maziar Heidari, Karen Palacio-Rodriguez, Hao Ruan, Sara Mingu, Alam Ahmad Hidayat, Mathis Bode, Mateusz Sikora, Gerhard Hummer, Edward A. Lemke

## Abstract

The nuclear pore complex (NPC) regulates the molecular traffic between nucleus and cytoplasm.^1,2^ Its permeability barrier is formed by intrinsically disordered proteins (IDPs) known as FG-nucleoporins (FG-NUPs), whose physical state has long been debated.^3–13^ Deciphering how FG-NUPs behave inside living cells is crucial for understanding how the NPC achieves selective and rapid transport. Here, by combining site-specific labelling with picosecond time-resolved fluorescence anisotropy, we reveal that FG domains exhibit nanosecond-scale, liquid-like mobility in live cells, yet undergo a liquid-to-solid transition in vitro. Experiments and coarse-grained molecular dynamics simulations further show that importin-β, a major nuclear transport receptor, and O-linked β-N-acetylglucosamine (O-GlcNAc), a key post-translational modification of FG-NUPs together stabilise the dynamic FG network. Finely balanced FG–FG interactions modulated by O-GlcNAcylation, along with FG–importin-β interactions, maintain liquidity and enrich importin-β near the NPC periphery, while extended FG domains remain in the central channel to form the transport barrier. These findings reconcile conflicting models of FG-NUP organisation^4,6,8,10,14,15^ and explain the recent observations in high-resolution MINFLUX studies of importin-β depletion from the pore centre in mammalian cells.^16^ Beyond resolving debates over FG-NUP behaviour, our study underscores the importance of studying IDPs in their cellular context, with broader implications for understanding IDP-related diseases, including viral infections, cancer, and neurodegenerative disorders.

## Introduction

Eukaryotic cells rely on the efficient communication between the nucleus and the cytoplasm for the regulated exchange of genetic information, signalling molecules, and metabolic products. This vital process is orchestrated by the nuclear pore complex (NPC), a massive macromolecular assembly embedded in the nuclear envelope. Central to the NPC’s function is a permeability barrier formed by intrinsically disordered proteins (IDPs) known as FG-nucleoporins (FG-NUPs), which regulate selective transport by allowing passive diffusion of small molecules while facilitating the transit of larger cargos in association with nuclear transport receptors (NTRs).^1,2,17–22^ Elucidating how FG-NUPs maintain both high selectivity and efficient transport is key to deciphering NPC function.

Probing the dynamic properties of FG-NUPs in living cells poses a significant challenge. The NPC’s highly crowded and nanoconfined environment limits the effectiveness of many traditional biophysical techniques. Single-molecule tracking (SMT), fluorescence correlation spectroscopy (FCS), and fluorescence recovery after photobleaching (FRAP) can assess the overall diffusion of IDPs in live cells,^23–26^ but do not typically resolve local motion of the disordered protein chain. Such segmental motion is particularly relevant for FG-NUPs, which are tethered to the NPC scaffold. While the mobility is reduced at the anchored terminus, the rest of the tethered peptide chain is free to move. Although atomic force microscopy (AFM) provides insights into mechanical and dynamic properties at millisecond timescales in surface-accessible systems,^3,27–29^ it is not readily applicable to intact NPCs in living cells. Consequently, much of our current understanding relies on in vitro experiments and computer simulations,^3–5,30–33^ which suggest that FG-NUPs can adopt a range of physical states—from flexible polymer chains^34,35^ and polymer brushes^14^ to phase-separated condensates exhibiting liquid-like, gel-like, or even amyloid-like properties.^4,6,12,13^ These disparate observations have generated considerable debate over the nature of the permeability barrier in the NPC, with some models favouring a liquid-like network and others suggesting a more solid-like structure.^3–11^ Our previous application of fluorescence lifetime imaging microscopy with Förster resonance energy transfer (FLIM-FRET) clarified conformational preferences of FG-NUPs in permeabilised cells.^36^ However, directly measuring their dynamics in live cells remained difficult. Addressing this gap is key to determining how FG domains behave under physiological conditions.

Here, we integrate site-specific protein labelling via genetic code expansion (GCE) employing orthogonally translating organelles (OTO) with picosecond time-resolved fluorescence anisotropy (TRFA) to directly measure FG-NUP segmental dynamics in living mammalian cells. For the vital NUP98, we reveal that its FG domains exhibit nanosecond-scale, liquid-like mobility inside the NPC, in stark contrast to their solid-like states in aged FG hydrogel condensates in vitro. We further demonstrate that importin-β, an abundant NTR, and O-linked β-N-acetylglucosamine (O-GlcNAc), a key post-translational modification (PTM) of FG-NUPs in metazoans, cooperatively maintain FG domain liquidity in cells and reconstituted condensates. In vitro, either factor alone can suppress FG-NUP aggregation and promote liquid-like behaviour, but phase separation of O-GlcNAc–modified FG domains with importin-β most closely recapitulates the nanosecond-scale mobility observed in cells, underscoring their synergistic role in preserving the dynamic permeability barrier of the NPC. In line with these results, coarse-grained molecular dynamics (MD) simulations show that importin-β–FG interactions, together with O-GlcNAc–modulated FG–FG interactions, are finely balanced to prevent solidification while maintaining NPC function.

Notably, in the optimal importin-β–FG interaction strength regime, not just liquidity is maintained, but importin-β is enriched at the NPC periphery, leaving a pronounced depletion as an “NTR void” in the central channel. At the same time, FG domains remain extended into the centre, forming an effective transport barrier. Our results thus reconcile puzzling observations from recent MINFLUX tracking studies,^16^ which observed NTR-bound cargos travelling predominantly near the NPC rim rather than through its geometric centre or uniformly through the channel (as would be expected for laminar flow and diffusive transport through a nanochannel, respectively). While the MINFLUX studies have reignited the discussion on the presence of a physical plug,^29,37–39^ our data show that the peripheral enrichment of NTRs follows naturally from their secondary function as chaperone-like solubilisers of the dense, extremely hydrophobic and thus aggregation-prone FG-NUPs. With interaction strengths matched to experiments, the simulations show dynamic FG-NUPs, NTRs concentrated at the NPC periphery, and transport optimally trading off NTR concentration and mobility.

By directly measuring the dynamics of FG-NUPs in live cells and elucidating how O-GlcNAcylation and importin-β keep them fluid inside NPCs for facile yet selective transport, our study establishes a unifying framework for NPC permeability barrier organisation. Beyond deepening the fundamental understanding of nuclear transport, our findings also set the stage to investigate how dysregulated FG-NUP behaviour may contribute to disease.^40^ Moreover, the methods we developed here by combining in-cell fluorescence measurements and computational modelling provide a versatile platform for studying other IDPs within nanocavities or biomolecular condensates in living cells, opening broader opportunities to probe their conformational dynamics in the native environments down to the nanoscale.

## Results

### Site-specific labelling of NUP98 in live cells

High-precision measurements of NUP98 dynamics in live mammalian cells require a labelling strategy that provides single-amino-acid accuracy without substantially altering protein behaviour. Building on our previously established synthetic orthogonally translating organelles-enabled genetic code expansion (GCE-OTO) approach^36,41^, we selectively reassigned the amber stop codon (TAG) to incorporate the noncanonical amino acid (ncAA) bicyclo[6.1.0]nonyne-lysine (BCN) at specific sites in NUP98 (Fig. 1a). The incorporated BCN allowed site-specific attachment of a small organic fluorophore via click chemistry.

**Fig. 1.**
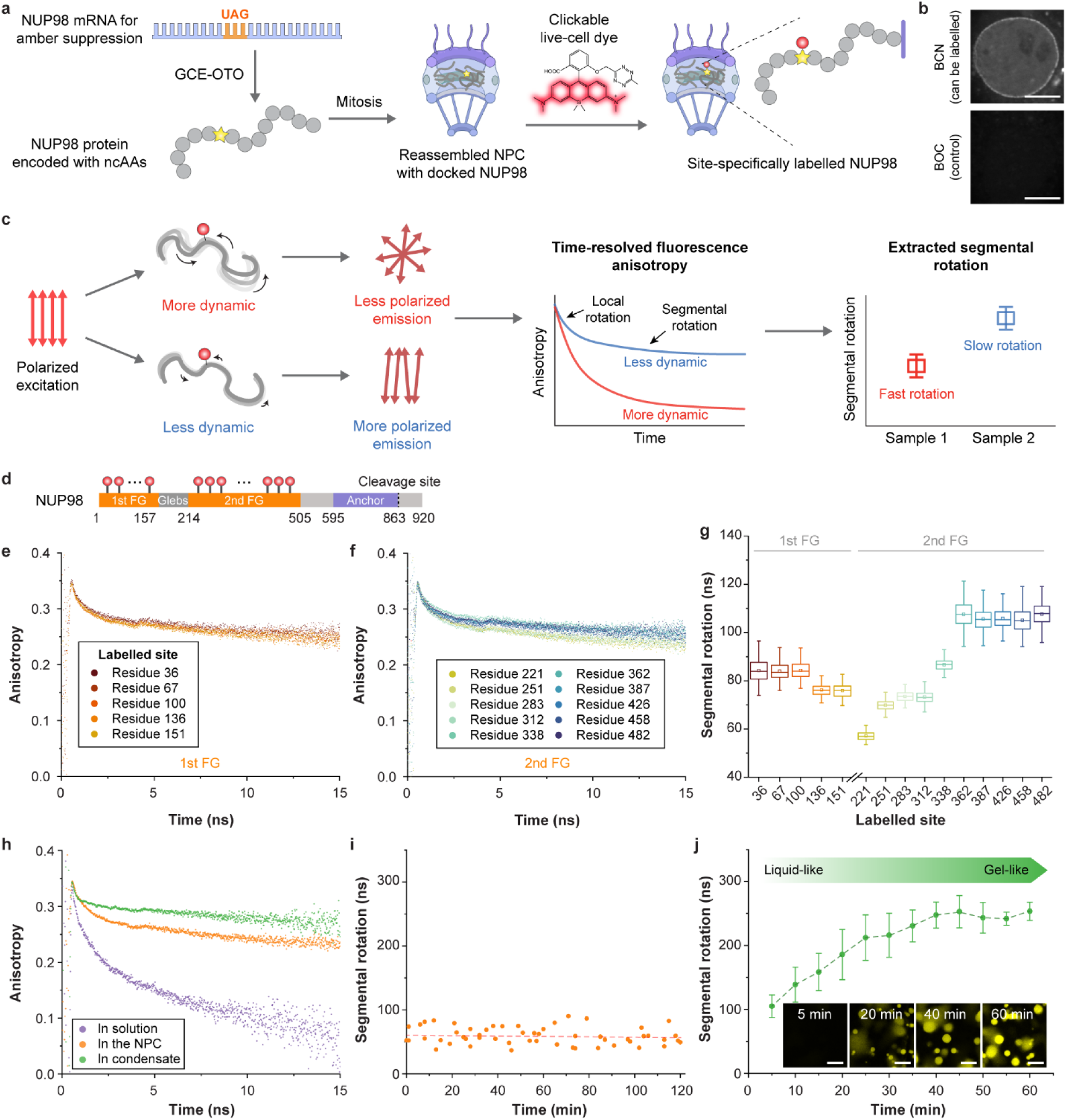
Site-specific labelling and segmental dynamics of NUP98 in live cells compared to in vitro systems. **a**, Schematic of site-specific labelling of NUP98 in the NPC using synthetic orthogonal translating organelle-enabled genetic code expansion (OTO-GCE). A noncanonical amino acids (ncAAs) is incorporated at defined positions, enabling selective labelling with live-cell dye HD653 via click chemistry. **b**, Fluorescence microscopy of COS-7 cells expressing a single-amber mutant (NUP98^A221TAG^) shows successful labelling with HD653 in the presence of bicyclo[6.1.0]nonyne-lysine (BCN, an ncAA that can be labelled), with no detectable signal when using *t*-butyloxycarbonyl-L-lysine (BOC, a control ncAA that cannot be labelled). Scale bars, 10 μm. **c**, Workflow of picosecond time-resolved fluorescence anisotropy (TRFA) measurements, used to extract local and segmental dynamics from anisotropy decay curves. **d**, Schematic of labelled sites spanning the first and second FG domains of NUP98. **e,f,** Representative TRFA decay curves for selected sites in the (**e)** first and (**f)** second FG domains. **g**, Segmental rotational correlation times across labelled sites reveal heterogeneous dynamics along the FG domains. Approximately 50 cells were analysed per mutant; due to the noisy single-cell fits, bootstrap resampling was applied prior to model fitting (see Methods). Box plots display the median (centre line), mean (central square), interquartile range (box), and whiskers extending to data points within 1.5×IQR. **h**, Representative anisotropy decay curves at residue 221 measured in the NPC of live cells, for isolated FG chains in solution, and in freshly formed FG condensates. **i**, Segmental dynamics at residue 221 in live cells remain stable over a two-hour imaging window, indicating no detectable molecular ageing in the native NPC. **j**, Time-dependent increase in segmental rotational correlation times at residue 221 during in vitro condensate maturation, reflecting a liquid-to-gel-like transition over one hour. Data represent mean ± s.d. from *n* = 3 independent replicates. Inset: Thioflavin-T staining shows emergence of amyloid-like structures during condensate ageing. Scale bars, 10 μm.

Our earlier attempts to label NUP98 with various live-cell dyes were hindered by low signal-to-noise ratios, limiting measurement accuracy.^36^ It was recently described that the silicon rhodamine derivative HDye653 (HD653) features a self-quenching tetrazine moiety attached via a short linker.^42^ Because HD653 fluorescence is activated upon reaction with BCN, the background signal is reduced, ensuring higher signal specificity. Furthermore, the short linker makes the HD653 dye a promising candidate for anisotropy studies.

To validate this approach, we expressed NUP98 bearing BCN at residue 221 (NUP98^A221TAG^, see Methods) in COS-7 cells. Upon reaction with HD653, a distinct nuclear rim signal was observed (Fig. 1b, Extended Data Fig. 1a, b), confirming efficient incorporation and labelling. By contrast, substituting BCN with a non-clickable ncAA, *t*-butyloxycarbonyl-L-lysine (BOC), abolished the signal. This comparison underscores both the specificity and the high signal-to-noise ratio achievable with HD653 labelling.

The short linker of HD653 minimises independent rotational motion of the dye (i.e., local motion around the dye-amino acid linker itself),^43^ allowing faithful reporting of peptide backbone rotation. TRFA measurements, a technique that measures the decay of fluorescence anisotropy over time (Fig. 1c), of purified enhanced green fluorescence protein (EGFP) site-specifically labelled with HD653 or the established anisotropy dye Cy3B^44,45^ yielded hydrodynamic radii that agree well with wild-type EGFP (Extended Data Fig. 1c, see Methods), confirming HD653 as a reliable anisotropy probe. Together, the small size, short linker, and fluorogenic properties of HD653 enable high-precision characterisation of FG-NUP dynamics within the nanosized NPC in live cells.

### Segmental mobility of NUP98 in live cells

To investigate the dynamic behaviour of NUP98 FG domains within the NPC, we employed TRFA. This approach provides two key parameters when the fluorophore is attached to an IDP like NUP98: the local motion correlation time of the attached fluorophore and the segmental rotational correlation time of the protein backbone (see Methods).^46,47^ The latter reflects the rotational diffusion of a relatively small region of the disordered protein—in this case, a segment around the labelled site—rather than the entire polypeptide chain. This parameter is particularly relevant for IDPs such as NUP98, where different regions can exhibit varying degrees of flexibility. Unlike nuclear magnetic resonance (NMR) spectroscopy, which has atomic resolution,^9^ TRFA probes motions across a region of the biopolymer. Based on previous studies,^47–49^ it is well established that structural correlations do not extend beyond approximately 50 amino acids, suggesting that our measurements primarily capture segmental motions within this range.

We systematically introduced BCN at 15 sites spanning the first and second FG domains of NUP98 and labelled them in live COS-7 cells with HD653 (Fig. 1d). Time-resolved fluorescence anisotropy decays were collected using a custom-built single-photon counting confocal microscope (see Methods), reporting segmental rotational correlation times ranging from 57 ± 2 ns to 108 ± 5 ns (Fig. 1e–g, Extended Data Fig. 1d, Supplementary Figs. 1-2). Interestingly, a near-sigmoidal trend emerged along the FG domains (Fig. 1g): sites closer to the N-terminus, which extend toward the pore centre, exhibited higher segmental mobility, whereas those near the anchoring domain were more restricted. This spatial gradient is consistent with previous findings that FG regions tethered near the NPC scaffold experience greater confinement.^3,50^

Although the intrinsic fluorescence lifetime of HD653 is only ∼4 ns, we could reliably separate fast local motions from slower segmental rotations occurring on the timescales of tens to hundreds of nanoseconds. This was enabled by accumulating large photon counts and robust multi-exponential fitting of anisotropy decay curves. Repeated measurements yield consistent trends, underscoring the reliability and sensitivity of our approach in capturing the subtle heterogeneity of segmental mobility in live cells. Probing even slower timescales (e.g., microseconds), however, would require complementary techniques like polarised FCS.^51^

### Segmental mobility of FG domains in vitro

To compare the dynamics of NUP98 FG domains in vitro with those observed in live cells, we performed TRFA measurements on two reconstituted systems: single NUP98 FG chains in solution and phase-separated FG condensates (Fig. 1h and 2a-b, Extended Data Fig. 1e, see Methods). For the same labelling site tested in cells (residue 221), isolated FG chains exhibited notably faster anisotropy decay (Fig. 1h), reflecting their higher flexibility. In contrast, freshly formed FG condensates (within the first five minutes) initially exhibited segmental dynamics similar to those observed in live cells, but gradually transitioned to a more solid-like state over an hour (Fig. 1i and j, Supplementary Movie 1). This was accompanied by a progressive slowdown in segmental motion, consistent with previous observations that FG condensates undergo molecular aging, eventually forming a solid-like hydrogel.^4,29^ Supporting this, thioflavin T (ThT) staining of the condensates showed no signal initially but intensified over time (Fig. 1j), indicating the emergence of amyloid-like structures during maturation.

**Fig. 2.**
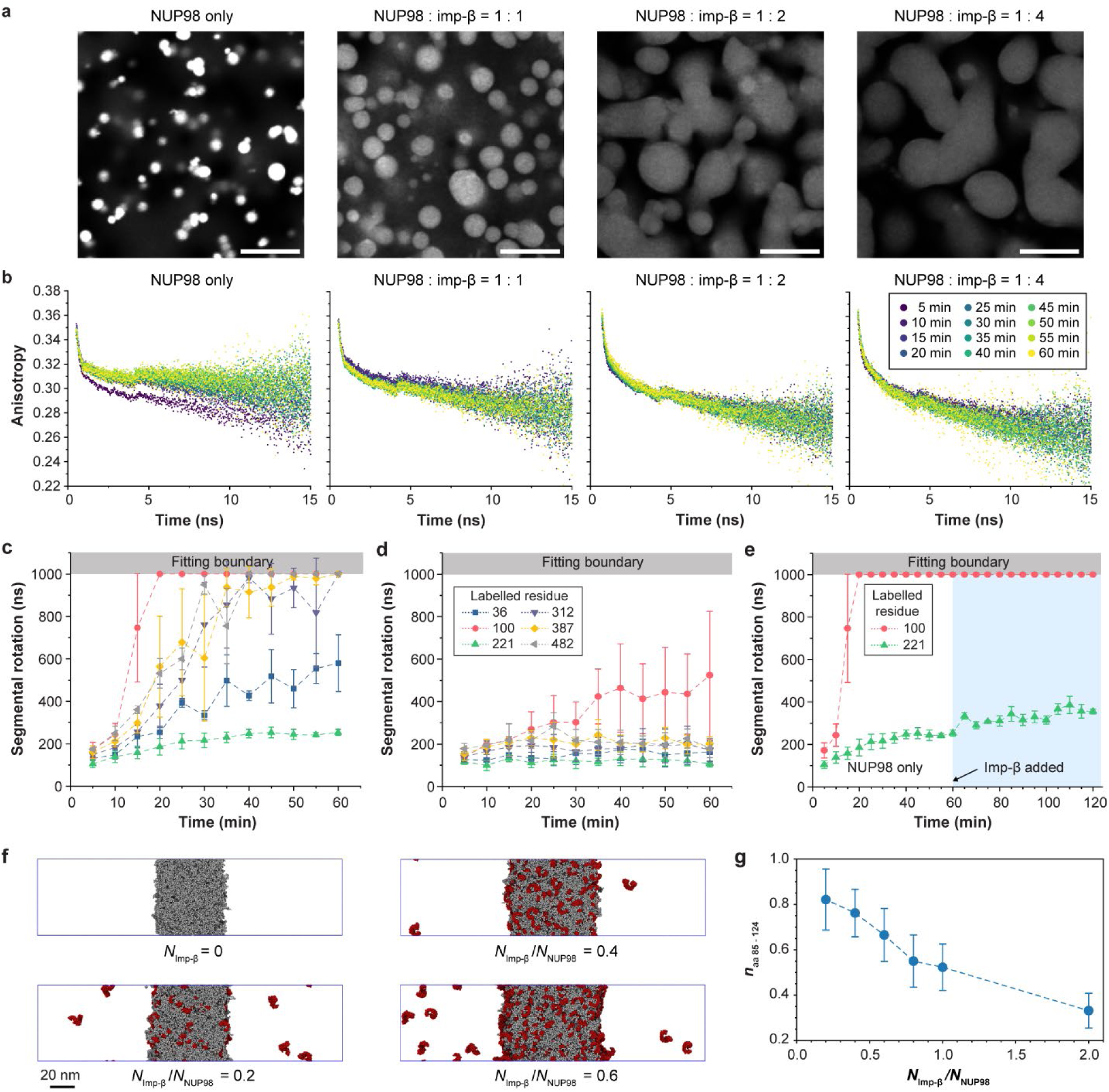
Importin-β enhances FG domain dynamics and stabilises liquid-like states. **a**, Phase separation of purified NUP98 FG domains with increasing concentrations of importin-β. Higher importin-β levels result in larger, more fluid condensates observed one hour after formation. All images use identical intensity scaling for direct comparison. Scale bar, 10 μm. **b**, Representative TRFA anisotropy decay curves recorded over one hour show faster decay with increasing importin-β concentrations, indicating enhanced segmental mobility of FG domains. **c**, Segmental rotational correlation times measured at multiple labelled sites in FG condensates. **d**, The same labelled sites as in (c), measured in FG condensates co-phase separated with importin-β, show accelerated segmental dynamics. **e**, Segmental rotational correlation times for FG condensates to which importin-β was added 1 h after formation, showing that mature condensates cannot be fully re-liquefied. Data in (c-e) represent mean ± s.d. from *n* = 3 independent replicates. **f**, MD simulation showing importin-β (red) partitioning into NUP98 FG condensates (grey) and inducing condensate swelling. Final configurations are shown for different molar ratios, *N*_Imp-β_ / *N*_NUP98_, as indicated. The FG–importin-β interaction strength was tuned to match the experimental binding affinity, ∈^∽^_FG–Imp-β_ = 0.42, and FG-FG interactions were set to reproduce in vitro behaviour^36^, ∈^∽^_FG–FG_ = 0.5. **g,** Importin-β dilutes the aggregation-prone^12^ segment of NUP98 (residues 85-124). Shown is the density of these segments, *n*_aa 85-124_, relative to that in a pure FG condensate within a probe sphere of 15 nm radius placed at the centre of the condensates. Mean and standard deviation were computed from the final 15 × 10^4^ *τ* of each simulation, sampled every 500 *τ*.

Analysis of additional labelling sites in NUP98 confirmed that all tested residues exhibited slower segmental mobility as the condensates matured into a hydrogel state (Fig. 2c). Notably, residue 100 showed the most pronounced reduction in mobility. This residue lies within a region recently identified by NMR as a particularly aggregation-prone FG-rich segment (amino acids 85–124)^12^. Thus, rather than being an outlier, this result aligns with the high propensity of this region to form amyloid-like assemblies. By contrast, no such transition was observed in live cells, even over extended measurement windows (e.g., two hours in Fig. 1i). These findings underscore a critical difference between the stable liquid-like dynamics of NUP98 observed in the cell and its time-dependent aggregation seen in vitro, raising the important question which physiological factors prevent FG-NUPs from undergoing gel-like maturation in their native environment.

### Importin-β tunes the segmental mobility of NUP98

Among the various NTRs, importin-β is particularly abundant in the NPC (Extended Data Fig. 2a), known to interact with FG motifs and proposed to exhibit chaperone-like properties.^52–55^ In vitro, phase separation of purified NUP98 FG domains with importin-β yielded larger, more liquid-like condensates with lower local FG concentration (Fig. 2a). TRFA measurements revealed a clear dose-response increase in segmental mobility; the importin-β:FG domain molar ratio increased from 1:1 to 4:1, the segmental rotational correlation time at residue 221 decreased from 120 ± 24 ns to 80 ± 7 ns, approaching the value measured in live cells (59 ± 14 ns) (Fig. 2d; Extended Data Fig. 2b–e). This mobility-enhancing effect was consistent across all tested sites, including residue 100, which lies within the particularly aggregation-prone region (85–124)^12^ (Fig. 2d and Extended Data Fig. 2f–j), suggesting a generalised role for importin-β in mitigating intermolecular FG–FG interactions. However, once condensates had matured for one hour, subsequent addition of importin-β failed to restore dynamics (Fig. 2e).

To gain mechanistic insights into how importin-β regulates FG-NUP dynamics, we performed coarse-grained MD simulations. We adapted a previously established model^36^ in which residues of the NUP98 FG domain and importin-β proteins are represented as single beads (see Supplementary Text for details). By matching the experimentally measured partitioning of importin-β into NUP98 FG condensates in vitro at varying molar ratios (Extended Data Fig. 3), we determined the interaction strength between FG-NUP and importin-β beads as ∈^∽^_FG–Imp-β_ = 0.42. Remarkably, this interaction perfectly balances that between the FG-NUP beads with ∈^∽^_FG–FG_ = 0.42 inside the NPC, yet it is weaker than the equivalent interaction of isolated chains and in condensates, ∈^∽^_FG–FG_ = 0.5, as obtained previously by matching root-mean-square inter-residue distances (*R*_E_) along NUP98 FG chains.^36^

With interaction strengths matched to the experiments (Extended Data Fig. 3c-e), we simulated the bulk FG-importin-β condensates. We observed that increasing the concentration of importin-β caused the condensates to swell (Fig. 2f, Supplementary Movie 2). Importin-β preferentially interacts with FG domains, diluting and statistically separating aggregation-prone segments (Fig. 2g). The observed dilution of the amyloid-forming region of residues 85-124^12^ mirrors the delay in the liquid-to-solid transition seen in our in vitro experiments (Fig. 2c,d).

### Importin-β shapes the NPC permeability barrier

To further examine the role of importin-β in the NPC function, we analysed its transport kinetics across the pore. We calculated the diffusion coefficient of importin-β within FG condensates (Extended Data Figs. 4-5, Supplementary Fig. 3) and estimated mean first passage times (MFPTs) for importin-β to traverse the pore. The resulting time scales, on the order of 1-10 ms, align well with previously reported experimental nuclear transport rates.^34,56–60^ Interestingly, we observed that the interactions between FG-NUPs and importin-β appear to be optimised to enhance the net rate of cargo transport (Extended Data Fig. 6), maintaining a balance between sufficient binding (to maintain selectivity) and rapid mobility (to allow efficient transport).

Simulations of importin-β-loaded NPCs under in-cell conditions (∈^∽^_FG–FG_ = ∈^∽^_FG–Imp-β_ = 0.42) further revealed a striking spatial organisation, where importin-β preferentially accumulated near the pore periphery (Fig. 3a). The radial gradient of the importin-β concentration leaves a conspicuous NTR void in the central channel (Fig. 3b left, Supplementary Fig. 4, Supplementary Movie 3), echoing the “empty centre” observed recently using two-colour MINFLUX microscopy.^16^ However, the interactions between importin-β and FG-NUP appear to be finely balanced, as the unique organization of importin-β molecules in the NPC is sensitive to their interaction strength with FG-NUPs: made 20% stronger, the importin-β molecules condense into the pore centre (Fig. 3c left; ∈^∽^_FG–Imp-β_ = 0.5, Supplementary Movie 4); made 20% weaker, the importin-β molecules delocalize across the entire pore (Supplementary Fig. 5; ∈^∽^_FG–Imp-β_ = 0.35, Supplementary Movie 5). These findings indicate that small changes in FG–importin-β affinity can shift importin-β localisation and affect transport efficiency (Extended Data Fig. 6c). Future high-resolution experiments, including MINFLUX,^16^ could test these predictions. We noted that small regions of high importin-β density in the MINFLUX experiments^16^ far on the cytoplasmic side are not captured, likely requiring a more refined model of NUP358.

**Fig. 3.**
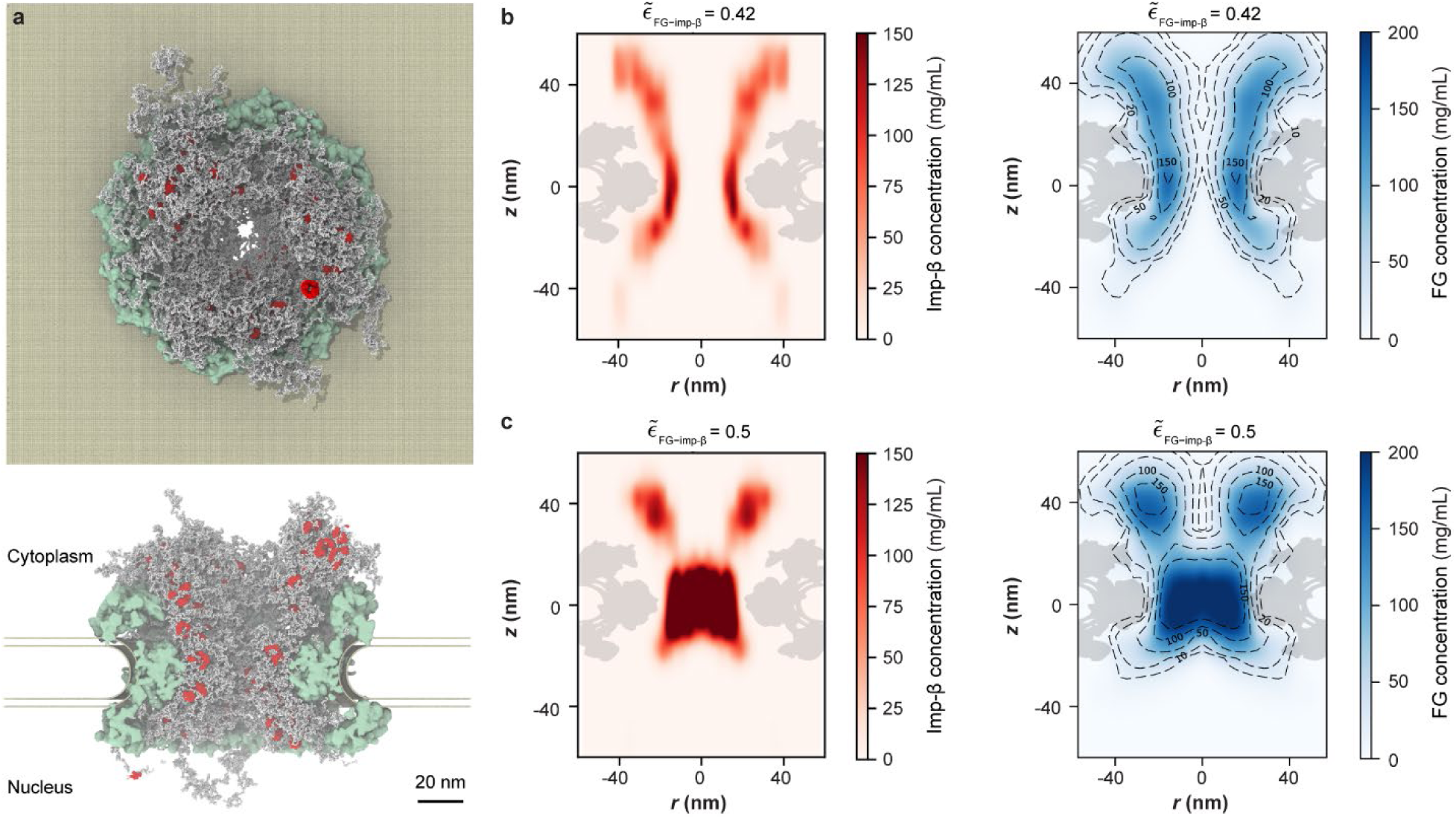
Importin-β shapes the FG-NUP permeability barrier of the NPC. **a**, Final configuration of MD simulation of an NPC with 100 importin-β molecules^77^ (∈^∽^_FG–FG_ = ∈^∽^_FG–Imp-β_ = 0.42). (Top) cytoplasmic side-view and (bottom) cross-sectional side view with all importin-β molecules (red) close (|*Z*| < 66 nm) to NPC (scaffold: green; FG-NUPs: grey; nuclear envelope: gold). **b,c,** Concentration map of (left) importin-β and (right) FG-NUPs in cylindrical coordinates (*r*: radial distance from the symmetry axis; *Z*: axial height from nuclear envelope centre with cytosol at *z* > 0) for interaction strength ∈^∽^_FG–Imp-β_ = 0.42 between FG-NUPs and importin-β proteins, as matched to the experimental affinity (**b**) and for artificially raised attractions, ∈^∽^_FG–Imp-β_ = 0.5 (**c**), with ∈^∽^_FG–FG_ = 0.42 fixed. The map was averaged over the last 5 × 10^4^*τ* sampled at intervals of 10^2^*τ*. Importin-β density maps from two additional simulations are shown in Supplementary Fig. 4.

Importantly, in importin-β–loaded pores the NUP98 FG chains remain highly dynamic and expanded, with their root-mean-square inter-residue distance, *R*_E_, closely matching previous in-cell FLIM-FRET measurements^36^ (Supplementary Fig. 6). Moreover, all major classes of FG-NUPs, depending on their grafting position, interpenetrate the FG–importin-β meshwork (Supplementary Fig. 7) and cooperate to shape the overall importin-β distribution. Collectively, their extended FG-rich tails fill the pore centre to form the permeability barrier (Fig. 3a, b right), while importin-β preferentially populates the periphery.

Tracking the mobility of individual importin-β molecules along simulated transport trajectories revealed pronounced dynamic heterogeneity within the pore (Supplementary Fig. 8). Importin motion is fastest towards the centre, consistent with the MINFLUX measurements^16^ and facilitated by the more rapid importin exchange between FG-NUPs in their more dilute tail region.

To connect the simulations with our TRFA findings, we analysed the bond auto-correlation at multiple NUP98 residues (Extended Data Fig. 7 and Supplementary Fig. 9, 10). In single chains, the segmental motion decayed fastest; in bulk condensates and in the NPC model, motion slowed, reflecting increased confinement imposed by neighbouring chains (Extended Data Fig. 7a). These trends closely parallel our experimental observations (Fig. 1h), further validating the simulation framework. We note that the aging effect was not considered in the simulations of FG condensates, where in the experiments the condensates exhibited slower dynamics over much longer timescales.

### O-GlcNAcylation tunes NUP98 dynamics

The partial ability of importin-β to delay, but not prevent, FG condensates solidification in vitro (Fig. 2d) suggested that additional cellular mechanisms must act to stabilise NUP98 dynamics in the NPC. The metazoan NPC is among the most heavily O-GlcNAc–modified subcellular structures, with FG-NUPs, particularly NUP98, being prominent glycosylation targets.^61^ O-GlcNAcylation has been implicated in modulating the material properties of FG-NUP hydrogels^62^ and has also been linked to changes in nuclear transport rates^63^. O-GlcNAcylation is a reversible post-translational modification in which a single GlcNAc moiety is attached to serine or threonine residues by O-GlcNAc transferase (OGT) and removed by O-GlcNAcase (OGA). However, its impact on nanosecond-scale segmental dynamics of FG-NUPs in living cells has remained unknown. To address this gap, we perturbed O-GlcNAc levels in live cells and in vitro.

Pharmacological inhibition of OGT with OSMI-4 in live COS-7 cells reduced global O-GlcNAc levels (confirmed by RL2 Western blotting) and led to significantly slower segmental motion near amino acid sites 221, 251, 283, 312, 338, 362, and 387 within the second FG domain, as measured by TRFA (Fig. 4a,b). In contrast, motion near sites 426, 458, and 482 was not significantly affected. This spatial pattern parallels predicted glycosylation propensities, with the more strongly affected regions corresponding to higher likelihoods of O-GlcNAc modification (Extended Data Fig. 8a). Conversely, inhibiting OGA with Thiamet-G maintained or modestly enhanced mobility (Fig. 4a), indicating that endogenous O-GlcNAcylation is already sufficient to support nanosecond-scale flexibility of FG domains under physiological conditions.

**Fig. 4.**
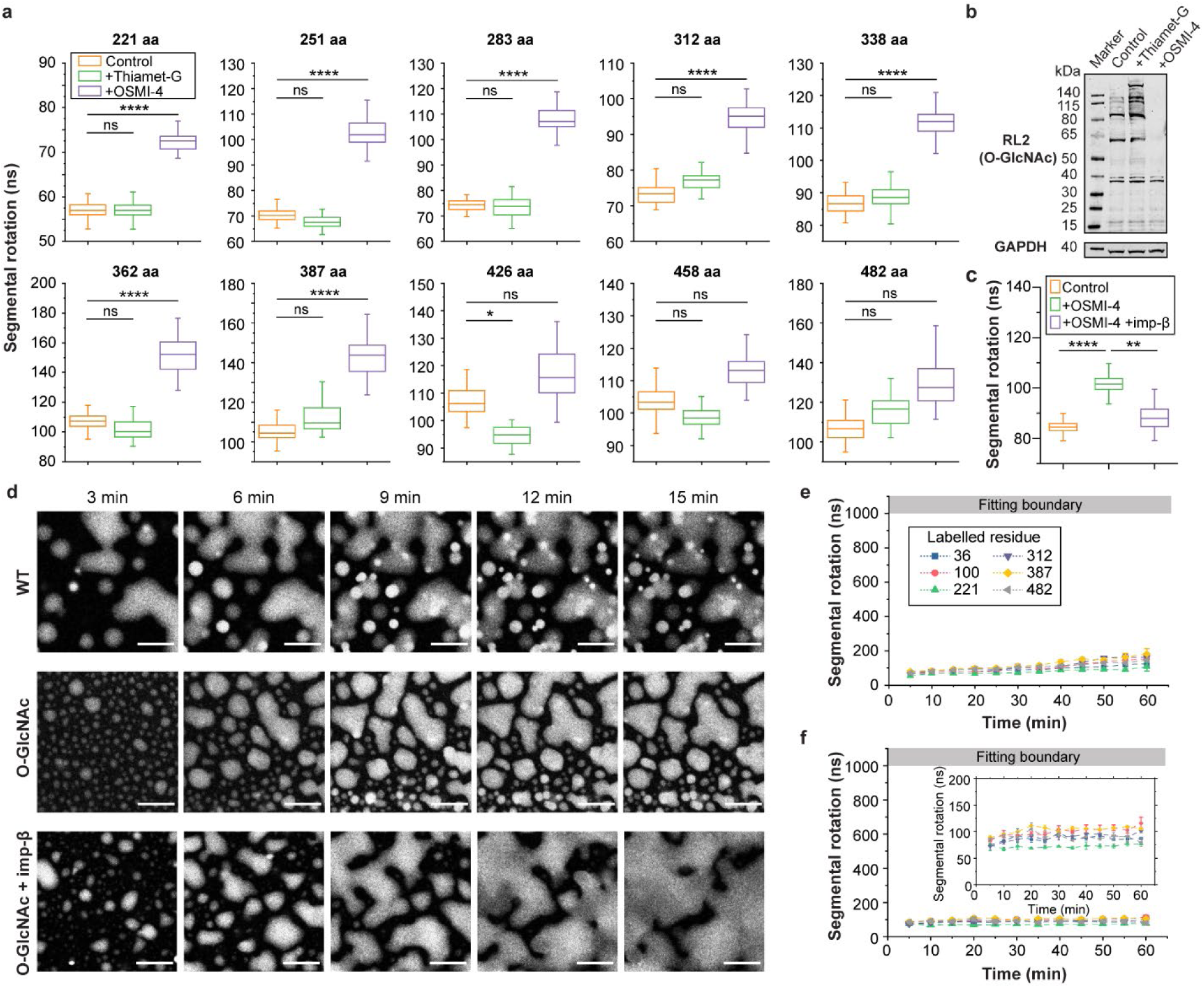
O-GlcNAcylation modulates FG domain dynamics and stabilises liquid-like states. **a**, Segmental rotational correlation times at selected FG domain sites in live COS-7 cells treated with 10 μM Thiamet-G (O-GlcNAcase inhibitor) or OSMI-4 (O-GlcNAc transferase inhibitor). Data are based on approximately 50 cells per condition. **b**, Western blot analysis of global O-GlcNAc levels (RL2) following 20 h treatment with Thiamet-G or OSMI-4. **c**, Overexpression of importin-β partially restored segmental rotational dynamics at residue 100 in OSMI-4–treated live cells. Data are based on approximately 80 cells per condition. For (a) and (c), 200 bootstrap replicates were generated by summing randomly resampled anisotropy decay curves before curve fitting. Box plots display the median (centre line), interquartile range (box), and whiskers extending to data points within 1.5×IQR. Statistical significance was assessed using two-sided bootstrap hypothesis testing, with thresholds: ns (not significant, p > 0.05); *p ≤ 0.05; **p ≤ 0.01; ***p ≤ 0.001; ****p ≤ 0.0001. **d**, Time-lapse fluorescence images showing phase separation of purified wild-type (WT) NUP98 FG domains, enzymatically O-GlcNAcylated FG domains, and phase separation of O-GlcNAcylated FG domains in the presence of importin-β in vitro. Scale bar, 10 μm. **e,f**, TRFA measurements of segmental rotational correlation times at multiple labelled sites in (**e**) O-GlcNAcylated NUP98 FG condensates and (**f**) O-GlcNAcylated NUP98 FG condensates phase separated with importin-β over a one-hour time course in vitro. Data represent mean ± s.d. from *n* = 3 independent replicates.

To determine whether the modulation effect of O-GlcNAcylation is recapitulated in vitro, we enzymatically glycosylated purified NUP98 FG domains and examined their behaviour in phase-separated condensates (Fig. 4d, Extended Data Fig. 8b–e). Glycosylated FG domains exhibited markedly delayed maturation, retaining liquid-like segmental dynamics for extended periods, in contrast to unglycosylated counterparts that progressively transitioned to solid-like states (Fig. 4d,e, Fig. 2c, Supplementary Movie 1). This effect was especially pronounced at residue 100, located within a region known for amyloidogenic propensity.^12^ Although mass spectrometry confirmed only partial glycosylation (approximately 17 GlcNAc moieties per chain; Extended Data Fig. 8c), this modification effectively suppressed condensation and delayed hydrogel formation, suggesting that local glycosylation enhances chain flexibility and disrupts pathological intermolecular FG–FG interactions.

We next asked whether importin-β can cooperate with O-GlcNAcylation to stabilise FG domain dynamics. Indeed, in vitro condensates formed from glycosylated NUP98 FG domains retained nanosecond-scale segmental mobility when phase separated in the presence of importin-β, closely reproducing the dynamics observed in live cells (e.g., at residue 221: 59 ± 14 ns in cells versus 72 ± 3 ns in vitro; Fig. 1i and 4d,f, Extended Data Fig. 3f, Supplementary Movie 1). This establishes a biophysically accurate in vitro model of the NPC permeability barrier down to the level of protein conformational dynamics.

Finally, to test this cooperativity in a cellular context, we overexpressed importin-β in OSMI-4–treated COS-7 cells. Remarkably, importin-β overexpression partially restored segmental mobility, rescuing faster rotational dynamics at residue 100, one of the most aggregation-prone regions identified in vitro (Fig. 4c). This rescue demonstrates that importin-β not only suppresses FG aggregation in vitro but can also compensate for impaired O-GlcNAcylation in cells, reinforcing its role as a physiological stabiliser of FG-NUP liquidity within the NPC.

### O-GlcNAcylation places FG-NUPs at a critical point for condensate formation

To mechanistically interpret our experimental findings, we built a near-atomistic Martini 2.2 model^64,65^ of a constricted human NPC, derived from the dilated-state structure (Protein Data Bank id, PDBid: 7R5J^66^), and simulated FG-NUP behaviour under varying levels of O-GlcNAcylation. We examined four glycosylation states, 0%, 18%, 30%, and 60% of candidate serine and threonine residues, representing progressively increased modification of FG domains (Supplementary Fig. 11). Protein–protein, protein–sugar, and sugar–sugar interactions were scaled to reproduce experimentally measured binding affinities^67,68^ (Supplementary Figs. 12–14).

In the absence of glycosylation (*f*_gly_ = 0), FG-NUPs collapsed into a surface condensate on the NPC scaffold (Fig. 5a; Supplementary Movie 6), leaving a large central void and producing NUP98 chain extensions inconsistent with in situ FLIM-FRET measurements (Fig. 5e). Increasing levels of O-GlcNAcylation progressively expanded the FG domains into the central channel (Fig. 5b–d; Supplementary Movies 7–9). At *f*_gly_ = 0.3, the extension *R*_E_ of NUP98 chains closely matched FLIM-FRET data (Fig. 5e). The 48 glycosites at *f*_gly_ = 0.3 closely match the 46 sites identified experimentally in *Xenopus* NUP98.^62^ Higher glycosylation levels yielded even more extended chains.

**Fig. 5.**
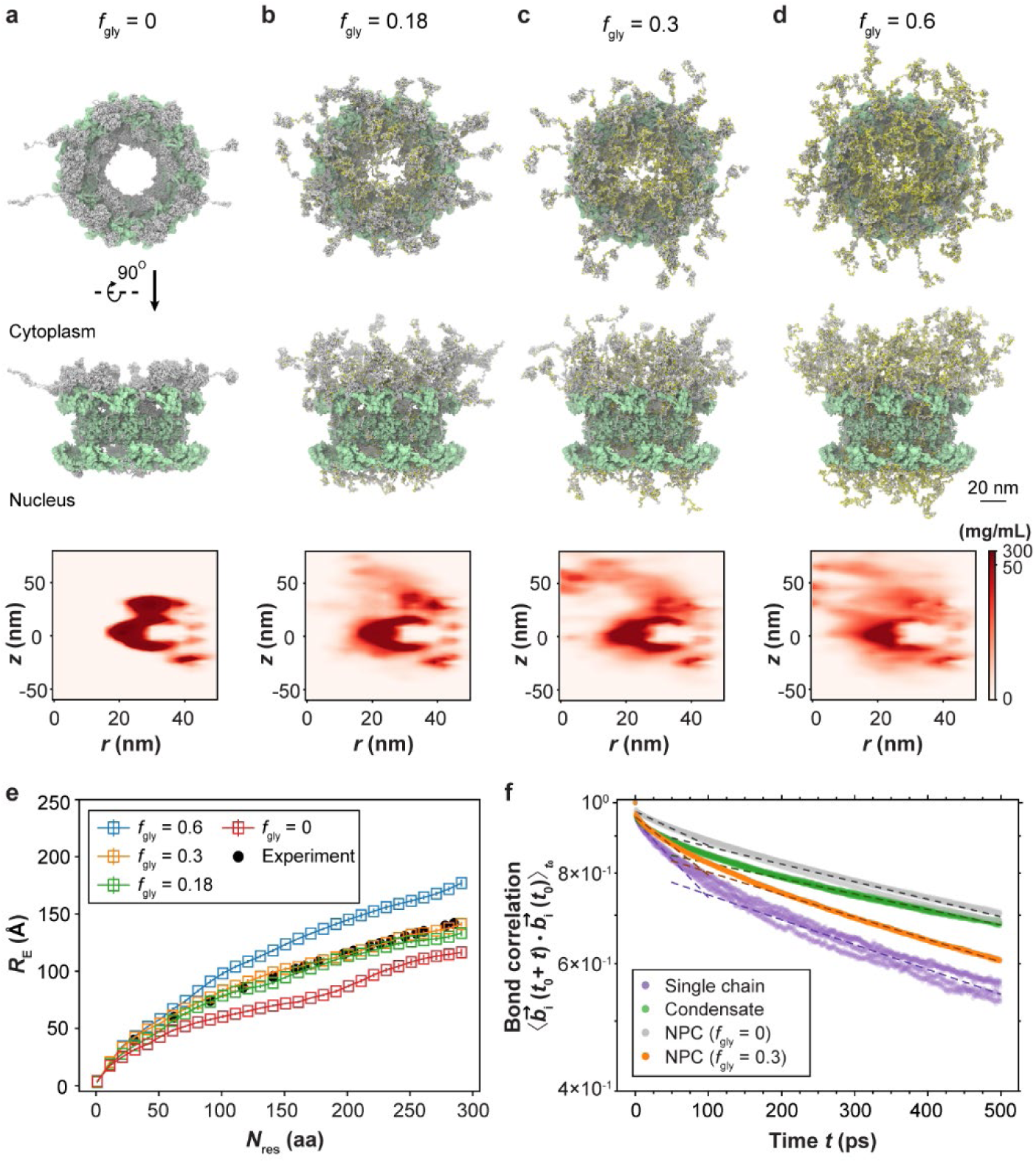
O-glycosylation controls FG-NUP fluidity in the NPC. **a-d**, Snapshots from Martini MD simulations of the NPC at increasing levels of O-glycosylation of FG-NUPs (NUP54, NUP58, NUP62, NUP98, NUP214, NUP358, and POM121). The fraction of glycosylated serine/threonine residues was (**a**) 0% to (**b**) 18%, (**c**) 30%, and (**d**) 60%. The top and middle panels show cytoplasmic-side view and cross-sectional side view of the final configuration (FG-NUPs, grey; glycosylated Ser/Thr residues, yellow; NPC scaffold, green). Bottom panels show axially averaged FG-NUP protein concentration (excluding glycan beads and NUP358) averaged over the final 600 ns. Colour scale: linear from 0 to 50 mg/mL, with concentrations >50 mg/mL shown in uniform dark red; bin size Δ*r* = Δ*z* = 3 nm. Total simulation times are (a) 3.5 μs and (b-d) 2 μs. **e**, Root-mean-square inter-residue distance (*R*_E_) of NUP98 FG domains within the NPC for the different glycosylation levels. Open squares and error bars denote the mean and standard error obtained from four non-overlapping 150-ns block averages. Experimental FLIM-FRET results are shown as black circles for comparison^36^. **f**, Auto-correlation function of bond vectors for the NUP98 bond between residues 221 and 222, *b*^⃗^_i_, computed from Martini simulations of NPCs without glycosylation (silver), with 30% glycosylation (orange), isolated FG chains (purple), and FG condensates (green). Curves were obtained from three independent 100-ns simulations sampled at 1-ps intervals. Dashed lines show exponential fits of the form *a* exp(−*t*/τ_fast_) and *a*^′^ exp(−*t*/τ_slow_) over time windows [0, 50 ps] and [50 ps, 500 ps], respectively (fit parameters in Supplementary Table 6). All snapshots were rendered with VMD.^78^

O-GlcNAcylation also accelerated backbone relaxation, mirroring the experimental trend in which NUP98 FG domains exhibit faster segmental dynamics inside the NPC than in unglycosylated in vitro condensates, yet slower dynamics than isolated chains (Figs. 1h and 5f, Supplementary Fig. 15). The effect of O-GlcNAcylation on FG-NUPs can be reproduced by weakening FG-NUP attractive interactions through *α*-scaling of protein-protein interactions (Extended Data Fig. 9, Supplementary Fig. 15). Remarkably, by matching the NUP98 extension to the FLIM-FRET measurements, we find the FG-NUPs to be near the critical point to phase separation also for the Martini model, confirming our earlier observations from the residue-level coarse-grained FG-NUP models^36^.

Together, the combination of Martini and residue-level coarse-grained simulations recapitulates key mesoscopic and microscopic features—from chain extension and segmental mobility to importin-β partitioning and transport behaviour—and bridges simplified in vitro studies with the crowded environment of the intact NPC. These results highlight how O-GlcNAcylation and importin-β jointly prevent solid-like aggregation of FG domains and generate the spatial and dynamic heterogeneity characteristic of the permeability barrier.

## Discussion

We combined site-specific labelling via OTO-GCE with picosecond TRFA to directly measure the segmental dynamics of FG-NUPs in living cells. Our experiments, together with MD simulations, provide a unified view of how the NPC maintains a selective yet dynamic permeability barrier. A key finding is that, under physiological conditions, NUP98—used here as a model FG-NUP—exhibits nanosecond-scale segmental mobility in live cells, including within regions that aggregate readily in vitro. This sharply contrasts with aged FG condensates, which rapidly transition into solid-like hydrogels.^6,12,13,62^ In these condensates, FG chains progressively lose mobility over minutes (Fig. 1j), in agreement with solid-state NMR data showing the development of amyloid-like interactions (Fig. 1j), as well as electron microscopy observations of fibre-like structures that form in certain FG-NUP hydrogels.^13,52^ Although previous in vitro and simulation studies proposed a spectrum of possible conformational states for FG domains,^3–11^ our direct in-cell measurements provide clear evidence that the NPC permeability barrier is liquid-like under physiological conditions, rather than forming the gel- or solid-like states often seen in vitro.^4–6,12^ This resolves a longstanding debate over the physical nature of FG-NUP assemblies in their native environment.

This discrepancy raises a key question: why do the same FG domains rapidly age into hydrogels in vitro, yet remain dynamic for days within one of the longest-lived protein assemblies in the cell?^23,69^ Classically, long-lived protein assemblies tend to adopt static or gel-like structures—amyloids being prime examples—whereas nucleocytoplasmic transport relies on ultrafast, reversible FG–FG and FG–NTR interactions. During transit, NTRs engage in thousands of sequential contacts, facilitated by nanosecond-scale FG segmental mobility, allowing cargo passage through the NPC on the millisecond timescale.^34,35^ Our findings reveal two regulatory mechanisms that cooperatively preserve FG-NUP liquidity: O-GlcNAcylation and importin-β. In in vitro condensates, each factor alone partially suppresses molecular aging, but together they recapitulate the nanosecond-scale segmental mobility observed in living cells (Extended Data Fig. 8f), thereby establishing an in vitro model that faithfully reproduces the liquid-like permeability barrier.

MD simulations further revealed how importin-β contributes to this stabilisation. Importin-β dilutes aggregation-prone FG segments, produces more swollen condensates in vitro, and enhances segmental mobility in situ. Although chaperone-like properties of importin-β had been proposed,^52,53,55^ our work explicitly demonstrates how this transport protein shapes segment-level dynamics of FG domains. Thus, importin-β performs a second essential function beyond cargo transport: it actively helps maintain the fluidity of the FG-based barrier, in line with the Karyopherin-centric model of the NPC barrier.^11^

Serendipitously, our simulations answered another major unresolved mystery in nuclear transport.^16,37–39,56^ Slightly overestimating the FG–importin-β interaction strength causes NTRs to cluster in the pore centre while reducing FG-NUP mobility (∈^∽^_FG–Imp-β_ = 0.5; Fig. 3b,c bottom, Supplementary Movie 4). However, under physiological conditions with experimentally matched interaction strengths, importin-β preferentially accumulates near the pore periphery, creating an apparent importin-β-void that is predominantly occupied by extended FG domains (∈^∽^_FG–Imp-β_ = 0.42; Fig. 3b,c top, Supplementary Movie 3). This spatial organisation explains recent MINFLUX observations showing NTR-bound cargos traveling primarily along the pore rim^16^ rather than through its geometric centre, as would be expected from laminar flow in a nanochannel. While those recent studies have reignited the discussion about the physical presence of a central plug,^37–39^ our data show that peripheral enrichment of NTRs is sufficiently explained by a pure physics-based effect, i.e., in a nanochannel densely packed with extremely high amounts of hydrophobic residues, NTRs with the right interaction strength are needed to prevent aggregation. Grafting on the NPC scaffold^36^ leads to a locally enhanced FG-NUP concentration at the pore periphery, which in turn leads to importins depletion at the centre. Together, these principles elucidate both the dynamic architecture of the NPC and the mechanisms that prevent molecular aging.

These findings prompt us to propose a dual-route model for nucleocytoplasmic transport. Small and medium-sized cargos that bind efficiently to importin-β travel rapidly along the periphery, where NTRs are enriched. By contrast, large cargos such as mRNPs or HIV capsids,^70^ which interact less efficiently with importin-β (and other NTRs) or have reduced mobility, may instead traverse the central channel. This model aligns with electron microscopy observations of a “central plug” which likely represents slow-moving or stalled large cargos.^37–39^ This framework reconciles multiple lines of evidence regarding FG cohesiveness, cargo trajectories, and transport efficiency, offering a unifying view of how the NPC adapts to diverse cargo requirements while maintaining strict selectivity.

Given the essential role of the NPC and its extremely slow turnover in postmitotic cells such as neurons, it is likely that further factors contribute to maintaining the liquid-like state. One intriguing possibility is that the diverse mixture of FG-NUPs, confined together within the nanocavity, stabilises the barrier via self-solubilising interactions, analogous to how β-synuclein suppresses α-synuclein aggregation.^71–73^ Although laborious to prove experimentally, this mechanism may explain why NPCs across species contain multiple distinct FG-NUPs (typically around ten types, with multiple copies) whose sequences are poorly conserved, even though in vitro a single FG-NUP type, such as NUP98, can form a permeability barrier at high concentration. Beyond FG-NUP diversity, other PTMs such as phosphorylation may further modulate FG dynamics.^74^ However, selectively altering NUP98 phosphorylation in live cells remains challenging due to the broad action of kinases and phosphatases. Furthermore, hyperphosphorylation of NUP98 is a crucial step in facilitating NPC disassembly.^75^ In contrast, NUP98 is among the most abundantly O-GlcNAcylated NUPs, and global perturbation of O-GlcNAc levels resulted in measurable and physiologically relevant changes in segmental mobility. Further studies using site-specific mutagenesis or phospho-mimetic variants could clarify how these modifications fine-tune NPC fluidity.

Alongside offering a mechanistic framework for NPC function, our study sheds light on how IDPs maintain functional dynamics in crowded intracellular environments. Given that FG-NUP dysfunction has been implicated in cancer, viral infection, and neurodegenerative disease,^40,76^ it is tempting to speculate that pathological states may arise when regulatory factors, such as PTMs or the importin-β gradient, become perturbed. More broadly, our approach—integrating in-cell fluorescence measurements with multiscale computational modelling—provides a powerful strategy for studying the conformational dynamics of IDPs in their native environment, with potential applications across diverse biological systems where dynamic heterogeneity underlies function.

## Acknowledgement

We thank the Wombacher laboratory for generously providing HDye653. We thank Dr. Saskia Hutten for providing purified O-GlcNAc transferase. We thank all of the members of the Lemke laboratory for helpful discussions, the core facilities of the Faculty of Biology at Johannes Gutenberg University Mainz, and the Protein Production Core Facility at the Institute of Molecular Biology Mainz for expert assistance. We thank Martin Beck for many insightful discussions. M.Y. acknowledges funding from the National Natural Science Foundation of China (Project No. 32471521), the Fundamental Research Funds for the Central Universities (2042025kf0029), Hubei Provincial Natural Science Foundation of China (2025AFA021) and MSCA Individual Fellowship (TFNUP 89410). E.A.L. acknowledges funding from the ERC-ADG grant ‘MultiOrganelleDesign’, as well as CRC1551 ‘Polymer concepts in cellular function’ of the Deutsche Forschungsgemeinschaft (DFG project number 464588647). E.A.L. and G.H. acknowledge funding from CRC 1507 ‘Membrane-associated protein assemblies, machineries, and supercomplexes’ of the Deutsche Forschungsgemeinschaft (DFG project number 450648163). M.H., K. P-. R. and G.H. thank the Max Planck Society for support. M.S. and A.A.H: project financed under Dioscuri, a programme initiated by the Max Planck Society, jointly managed with the National Science Centre in Poland, and mutually funded by Polish Ministry of Education and Science and German Federal Ministry of Education and Research. K.P.R. acknowledges support from the “Hessen Horizon Marie Skłodowska-Curie-Stipendium” program. M.H. thanks Dr. Marc Siggel for insightful discussions and assistance in simulating large-scale nuclear pore complex models. This project received access to the JUPITER supercomputer, which is funded by the EuroHPC Joint Undertaking, the German Federal Ministry of Research, Technology and Space, and the Ministry of Culture and Science of the German state of North Rhine-Westphalia, through the JUPITER Research and Early Access Program (JUREAP).

## Author contributions

E.A.L., G.H. conceived the project. M.Y. designed the experimental study. M.Y. performed experiments together with H. R. and S. M.. M.H. and K.P.-R. performed the MD simulations with help from A.A.H. and M.S. for glycan modelling, and M.B. for JUPITER use. M.Y. and M.H. analysed the experimental and simulation data, and co-wrote the original draft together with E.A.L. and G.H. All authors contributed to manuscript editing and approved the final manuscript.

## Data availability

All data are available in the main text or the supplementary materials.

## Materials & Correspondence

All plasmids are available via cost-free academic material transfer agreement.

## Methods

### Cell culture, transfection, and labelling

#### Cell culture

COS-7 cells (Sigma 87021302) were cultured in Dulbecco’s Modified Eagle Medium (DMEM, Thermo Fisher 41965-039) supplemented with 10% (v/v) fetal bovine serum (Sigma F7524), 1% Penicillin-Streptomycin (Thermo Fisher 15140-122), 1% L-glutamine (Thermo Fisher 25030-081), and 1% sodium pyruvate (Thermo Fisher 11360-070). Cells were maintained at 37 °C in a humidified 5% CO₂ atmosphere, passaged every 2-3 days, and used within 15-20 passages.

#### Transfection

COS-7 cells were seeded on 35-mm imaging dishes (ibidi 81158) 24 hours before transfection to reach 60-70% confluency. Cells were transfected with plasmids listed in Supplementary Table 1 using jetPRIME® reagent (Polyplus) according to the manufacturer’s instructions. Four to five hours post-transfection, the medium was replaced with fresh medium containing 10 mM HEPES and 50 μM bicyclo[6.1.0]nonyne-lysine (BCN) for specific labelling or 50 μM *t*-butyloxycarbonyl-L-lysine (BOC) for control experiments. Where indicated, 10 μM Thiamet-G (MCE HY-12588) or 10 μM OSMI-4 (MCE HY-114361) was added at this stage to modulate cellular O-GlcNAcylation levels and maintained throughout all subsequent steps.

#### Live-cell labelling

At 20-24 h post-transfection, cells were washed with medium containing 50 μM BOC and 10 mM HEPES and incubated at 37 °C for 2 hours to remove excessive BCN. Subsequently, cells were labelled with 2 μM HDye653-tetrazine (HD653) in the serum-free culture medium at 37 °C for 30 minutes. For O-GlcNAc perturbation experiments, the labelling buffer also contained 10 μM Thiamet-G or 10 μM OSMI-4, as appropriate, to maintain treatment consistency. For comparison groups, other tetrazine-functionalized dyes, including SiR-tetrazine (SiR-tz) and Janelia Fluor 646-tetrazine (JF646-tz), were used following the same labelling procedure. After labelling, cells were washed with 1× PBS and imaged immediately in FluoroBrite™ DMEM (Thermo Fisher, A1896701) supplemented with 10% (v/v) fetal bovine serum, 1% L-glutamine, 1% sodium pyruvate and 10 mM HEPES. Imaging was performed at room temperature with the dish lid closed to maintain environmental stability for the ∼2-hour imaging session.

### Protein expression, purification, and labelling

#### Expression and purification of NUP98 FG domain

Homo sapiens NUP98 FG domain (residues 1-505) was cloned into a pQE-14his-TEV vector. As in previous work,^36,79^ the GLEBS domain (residues 157–213) was excluded to avoid potential misfolding of this structured domain upon rapid dilution from denaturant during phase separation (see *in vitro phase separation assay* below); intrinsically disordered FG regions, by contrast, readily regain their dynamic ensemble under native buffer conditions. The plasmid was transformed into *E. coli* BL21 AI cells and grown in terrific broth supplemented with 50 µg/mL of kanamycin at 37 °C with shaking at 200 rpm. Protein expression was induced with 0.02% (w/v) arabinose and 1 mM IPTG when the culture reached OD_600_ ∼0.6, followed by 16 h incubation at 18 °C.

Cells were harvested and lysed using a high-pressure cell disruptor (CF1, Constant Systems) in lysis buffer containing 6 M guanidine hydrochloride (GdmCl), 0.2 mM TCEP, 20 mM imidazole, and 50 mM Tris-HCl (pH 8.0). Lysates were clarified by centrifugation (12,000 × g, 1 h, 4 °C) and incubated with Ni-NTA resin for 2 hours at 4 °C. The resin-bound protein was washed with 2 M GdmCl buffer and eluted with 500 mM imidazole in 2 M GdmCl and 50 mM Tris-HCl (pH 8.0). The His-tag was cleaved overnight with TEV protease after dialysing into a buffer containing 0.5 M GdmCl and 50 mM Tris-HCl (pH 8.0). The cleaved protein was further purified via Ni-NTA resin to remove uncleaved protein and TEV, then subjected to size-exclusion chromatography (Superdex 200, Cytiva) in 2 M GdmCl, 0.2 mM TCEP, and 50 mM Tris-HCl (pH 8.0). Protein purity was confirmed by SDS-PAGE and Coomassie staining. Concentrated fractions (∼15 mg/mL) were measured using a BCA protein assay kit (Thermo Fisher) and stored at −80 °C.

#### In vitro O-GlcNAcylation of purified NUP98 FG domain

Purified NUP98 FG domain was enzymatically glycosylated using recombinant O-GlcNAc transferase (OGT1) and UDP-GlcNAc (Sigma U4375). The reaction mixture contained 10 µM NUP98 FG domain, 1 µM OGT1, and 1 mM UDP-GlcNAc in glycosylation buffer (50 mM Tris-HCl, pH 7.5, 200 mM NaCl, 20 mM MgCl₂, 0.2 mM TCEP, and 1% Tween-20). The reaction was incubated at room temperature for 16 h with gentle rotation. After glycosylation, Ni-NTA resin was added to remove His-tagged OGT1. The glycosylated NUP98 was further purified via size-exclusion chromatography (Superdex 200, Cytiva) in 2 M GdmCl, 0.2 mM TCEP, and 50 mM Tris-HCl (pH 8.0), and concentrated to ∼40 mg/mL.

#### Labelling of purified FG Domain

Purified NUP98 FG domains with single cysteine mutations (at residues 36, 100, 221, 312, 387, or 482) were buffer-exchanged to 4 M GdmCl, 1× PBS, 0.1 mM EDTA, and 0.2 mM TCEP (pH 7.0). Proteins were labelled with Cy3B (GE Healthcare, PA63131, for TRFA measurements) or Alexa Fluor 594 maleimide dye (for quantification of concentration of FG condensates) at a 1:2 molar ratio overnight at 4 °C. Excess dye was quenched with 10 mM DTT and removed by ultrafiltration using 3 kDa MWCO centrifugal filter. Labelled proteins were further purified using size-exclusion chromatography (Superdex 200) to remove free dyes and aggregates. Final concentrations were determined by an absorbance spectrometer Duetta (Horiba), and labelled proteins were flash-frozen and stored at −80 °C.

#### Expression, purification and labelling of importin-β

The *Homo sapiens* importin-β coding sequence was cloned into a pTXB3-intein-CBD-12His vector and transformed into *E. coli* BL21-AI cells. Cultures were grown in terrific broth with 50 µg/mL kanamycin at 37 °C, 200 rpm. At OD₆₀₀ = 0.4–0.6, protein expression was induced with 0.02% (w/v) arabinose and 1 mM IPTG, followed by incubation for 5–6 h at 37 °C. Cells were lysed by high-pressure homogenization in lysis buffer (50 mM Tris, 650 mM NaCl, 5 mM MgCl₂, 5 mM imidazole, 1 mM PMSF, 0.2 mM TCEP, pH 7.0). Lysates were clarified by centrifugation (12,000 × g, 1 h, 4 °C), and the supernatant was incubated with Ni-NTA resin pre-equilibrated in lysis buffer for 2 h at 4 °C. The resin was washed with lysis buffer containing 10 mM imidazole and 1.5 M NaCl, and importin-β was eluted with 400 mM imidazole. Eluate was dialysed into imidazole-free buffer and subjected to a second Ni-NTA step to remove cleaved tags and contaminants. The flow-through was further purified by size-exclusion chromatography (Superdex 200, Cytiva) in 50 mM Tris, 650 mM NaCl, 5 mM MgCl₂, and 0.2 mM TCEP (pH 7.0). Purity was assessed by SDS-PAGE. The final protein was concentrated to ∼500 μM, flash-frozen in liquid nitrogen, and stored at −80 °C.

#### Immunostaining of endogenous importin-β

COS-7 cells labelled with HD653 were fixed with 2% paraformaldehyde (PFA) in PBS for 10 min, washed twice with PBS, and permeabilized with 0.5% Triton X-100 in PBS for 15 min. After two additional PBS washes, cells were blocked with 3% BSA in PBS for 90 min and incubated with anti-KPNB1 antibody (Abcam, ab2811) followed by Alexa Fluor 488–conjugated anti-mouse secondary antibody (Thermo Fisher, A-11001). Immunofluorescence confirmed that endogenous importin-β colocalized at the nuclear envelope after live-cell labelling (Extended Data Fig. 1b).

#### Estimation of endogenous importin-β concentration at the nuclear pore

To estimate the local concentration of importin-β at the nuclear pore, free Alexa Fluor 488 dye solutions of known concentrations (1 µM, 10 µM, 100 µM, and 1 mM) were added to the immunostained cells as described above. Nuclear pore fluorescence remained clearly visible above the background up to 100 µM free dye but became indistinct at 1 mM (Extended Data Fig. 2a), indicating that the effective local concentration of importin-β at the nuclear pore is on the order of millimolar.

#### Western blotting of O-GlcNAc levels

To verify that pharmacological perturbations altered global O-GlcNAcylation, COS-7 cells were cultured in medium containing 10 μM Thiamet-G or 10 μM OSMI-4 (as described above) and lysed in Laemmli buffer. Proteins were resolved by SDS–PAGE, transferred to nitrocellulose membranes, and probed with anti-O-linked N-acetylglucosamine antibody RL2 (Invitrogen, MA1-072), followed by IRDye 800CW goat anti-mouse IgG secondary antibody (LI-COR Biosciences; Fig. 4b).

### Time-resolved fluorescence anisotropy (TRFA) measurements

#### Instrument configuration

TRFA measurements were conducted on a custom-built confocal microscope equipped with picosecond pulsed laser diode (560 nm: LDH-D-TA-560, PicoQuant; 660 nm: LDH-D-C-660, PicoQuant) controlled by a Sepia II PDL 828 multichannel laser driver.^36^ The excitation beam was coupled into a single-mode polarisation-maintaining optical fibre (KineFLEX-P-2-S-405/640-2.5-2.5-p2) and passed through a Glan-laser polariser (Thorlabs) to ensure linear polarisation. The beam was directed through a FLIMbee laser scanning system (PicoQuant) and focused onto the sample via a 60× water immersion objective lens (SR Plan Apo IR, 1.27 NA, Nikon). Emission light was collected and spatially filtered through a 100 μm pinhole for live-cell measurements and a 50 μm pinhole for in vitro samples. The fluorescence was split into parallel and perpendicular polarisation components using a polarising beam splitter. Each component was further separated spectrally by dichroic filters (ZT561 RDC and T647 LPXR, Chroma), passed through bandpass filters (Cy3B: 612/69 BrightLine HC, Semrock; HD653: ET700/75m, Chroma), and detected with hybrid photon-counting detectors (PMA Hybrid 40, PicoQuant). Signals were recorded with a HydraHarp 400 time-correlated single photon counting (TCSPC) module, providing a time resolution of 16 ps. Data were acquired using SymPhoTime 64 software (v2.6, PicoQuant). An instrument response function (IRF) was measured daily using a freshly prepared saturated solution of potassium iodide and Erythrosine B to account for instrumental timing variations. Temporal offsets between parallel and perpendicular polarisation detection channels were pre-aligned using the measured IRFs. We note that a small discontinuity around ∼4 ns is observed in the raw anisotropy decay curves; this arises from weak optical reflections in the TCSPC setup and is accounted for in the analysis by reconvolution with the measured IRF.

#### Live-cell measurements

The acceptor intensity at the nuclear rim was first verified under 660 nm laser excitation (optimised to 50 μW average power at 40 MHz) to assess the expression level, ensuring comparable expression levels and avoiding overexpression artefacts. Data acquisition was performed using T3 mode, with fluorescence photons collected separately by parallel and perpendicular detectors. Imaging parameters included a pixel size of 100 nm, an image size of 256 × 256 pixels, a pixel dwell time of 150 µs, imaging time of 5 min. Regions of interests (ROIs) corresponding to the nuclear rim were selected, where thresholding algorithms were applied to identify ROIs based on pixel intensity and average fluorescence lifetime. The time-resolved intensity curves for parallel and perpendicular channels were extracted respectively, for further analysis.

#### In vitro phase separation assay

The purified and labelled NUP98 FG domain was mixed with unlabelled protein at a molar ratio of 1:5000 in 2 M GdmCl, 1× PBS, pH 7.0. To induce phase separation, 1 µL of this mixture was rapidly diluted into 24 µL transport buffer (20 mM HEPES, 110 mM KOAc, 5 mM NaOAc, 2 mM Mg(OAc)_2_, 1 mM EGTA, 2 mM DTT, pH 7.3) on a chambered coverslip (ibidi 81507) and immediately imaged using a custom-built confocal microscope.^36^ The final total concentration of NUP98 in the system was 10 µM for unlabelled and 2 nM for labelled FG domain. The residual GdmCl concentration after dilution was ∼80 mM, which did not interfere with droplet formation.

At 10 μM, wild-type FG domains consistently formed micrometer-scale condensates suitable for TRFA measurements. In contrast, enzymatically O-GlcNAcylated FG domains exhibited reduced phase separation efficiency at this concentration and failed to form droplets large enough for reliable imaging (Extended Data Fig. 8d). To ensure comparable condensate size and morphology across experimental conditions, glycosylated FG domains were used at a higher concentration (25 μM; Fig. 4d).

For co-phase separation assays, importin-β was pre-diluted in transport buffer to the desired concentration before mixing with the FG domain. Wild-type FG domains were used at 10 μM, with importin-β added as indicated. For glycosylated FG–importin-β mixtures, 25 μM glycosylated FG domain and 10 μM importin-β were used to ensure consistent droplet formation. The residual GdmCl concentration (∼80 mM) was kept the same for all measured conditions.

The formed condensates were imaged over a 1-hour period using 560 nm laser excitation with an average power of 35 μW at 40 MHz. The imaging field was changed every 5 minutes during the imaging session to measure fresh condensates in different areas, avoid photobleaching and ensure representative sampling. Imaging parameters included a pixel size of 150 nm, an image size of 256 × 256 pixels, a pixel dwell time of 203 μs, and a time resolution of 16 ps. Fluorescence photons were detected in T3 mode and collected separately from perpendicular and parallel detectors. FG-condensates were identified as ROI based on the intensity, and the time-resolved intensity curves were extracted using SymPhoTime 64 software.

#### Single FG Chain in solution

For measurements on single FG chains, purified and labelled FG-NUPs were diluted in transport buffer to a final concentration of 10 nM. Similar imaging and analysis procedures were performed for the FG condensates as described.

### Quantification of dense- and dilute-phase concentrations of FG condensates in vitro

To determine the concentration of FG-NUPs in the dense phase of NUP98 FG condensates, phase separation assays were performed using mixtures of unlabelled wild-type NUP98 FG domain and trace amounts of Alexa Fluor 594–maleimide–labelled FG domain at molar ratios of 2000:1, 5000:1, or 10000:1. Condensates were formed as described in the in vitro phase separation assay section and immediately centrifuged at 4300 × g for 5 min to sediment the dense phase. Fluorescence intensity from the dense phase was measured using a custom-built confocal microscope equipped with a 560 nm pulsed laser (average power ∼70 μW, 25 μm pinhole). Dense-phase regions were identified based on their high fluorescence intensity and characteristic morphology. A calibration curve was generated using known concentrations of free Alexa Fluor 594 in matched buffer conditions. To correct for differences in fluorescence efficiency between free dye and dye-conjugated protein, quantum yields of both species were independently measured using a Duetta fluorometer, referencing standard dyes with known quantum yields. The relative quantum yield of the labelled protein was used to adjust the calculated protein concentration.

Dilute-phase concentrations were measured by adjusting the focusing layer much higher than the condensate layer, and using identical calibration and correction procedures. For NUP98 FG, a 20:1 unlabelled:labelled ratio was used to ensure sufficient signal in the dilute phase while avoiding perturbation of phase behaviour. Small dynamic nanoclusters were occasionally detected in the dilute phase for wild-type NUP98 FG domain when no importin-β was added (Extended Data Fig. 3d).

These experimentally determined concentrations served as constraints for selecting the FG–importin-β interaction strengths in our coarse-grained MD simulations (Extended Data Fig. 3e).

### Anisotropy decay analysis

TRFA decay, *r*(*t*), was determined by collecting fluorescence emissions parallel (*I*_∥_(*t*)) and perpendicular (*I*_⊥_(*t*)) to the excitation polarisation, and calculated as^80^

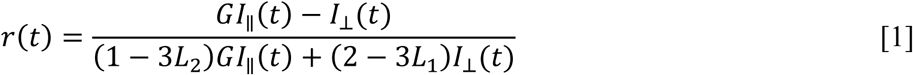

where *L*_1_ and *L*_2_ are polarisation mixing factors caused by the high numerical aperture of the objective lens and *G* accounts for the differential detection efficiencies between the parallel and perpendicular polarisation components:

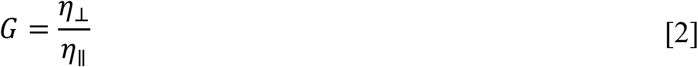

To determine *L*_1_ and *L*_2_, the fluorescence decays of a reference sample, yellow fluorescent protein (YFP), were measured. YFP has a known fluorescence rotational relaxation time (*ρ* = 16 ns^81^). The fluorescence decays for the parallel (*I*_par_) and perpendicular components (*I*_per_) of YFP were globally fitted using the following equations,^81^

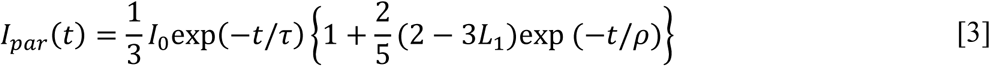

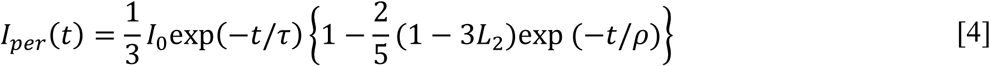

where *D*_0_ is the initial fluorescence intensity, *τ* is the fluorescence lifetime, and *ρ* is the rotational correlation time of YFP. *G* factor was determined by measuring the ratio of the average intensities in the perpendicular and parallel channels using a solution of tris(2,2’-bipyridyl)dichlororuthenium(II) chloride ([Ru(bpy)_3_]Cl_2_), which exhibits isotropic rotational motion.

### Fitting procedures and statistical analysis

For an IDP, *r*(*t*) can be modelled using a double-exponential decay,^46^

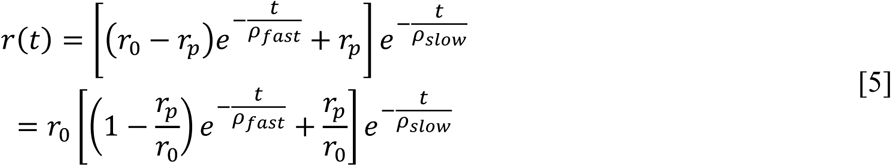

where *r*_0_ is the fundamental anisotropy of the fluorophore, *ρ*_fast_ is the rotational correlation time for local (fast) motions, *r*_p_ is the residual anisotropy from slower, larger-scale chain motions, and *ρ*_slow_ is the segmental rotational correlation time. When *ρ*_fast_ is much smaller than *ρ*_slow_, Eq. 5 can be approximated and simplified to,

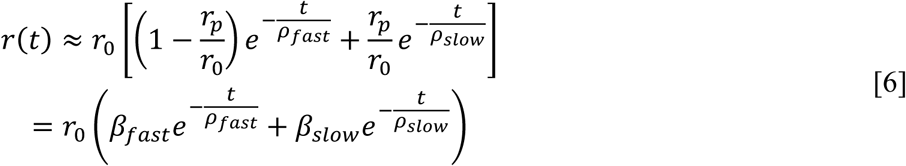

Where *β*_fast_ =1−*r*_p_/*r*_0_ and *β*_slow_= *r*_p_/*r*_0_ represent relative amplitudes of the fast (local) and slow (segmental) components, respectively.

By fitting the obtained anisotropy decay *r*(*t*) to Eq. 1 and 2 using PAM,^82^ detailed insights into the dynamics of distinct protein regions were extracted, revealing both local and segmental motion contributions. We have also tried fitting with three-exponential decay model. However, in all cases, the third component was either negligible or degenerate with one of the existing components.

For live-cell measurements, fluorescence anisotropy decay curves were acquired from ∼50 single cells per condition. In Fig. 1i, segmental rotational correlation times were calculated on a per-cell basis to capture temporal trends. For all other experiments, we applied a non-parametric bootstrap strategy to improve signal-to-noise and ensure robust curve fitting. Specifically, for each of 200 bootstrap replicates per condition, single-cell decay traces from the nuclear rim were resampled with replacement, summed, and fit to Eq. 5 to estimate the segmental rotational correlation time. We selected a bootstrap depth of 200 because increasing the number of replicates beyond this point yielded highly similar mean estimates and variance, indicating convergence (Supplementary Fig. 2). To assess perturbation effects, we compared bootstrap distributions from drug-treated and matched control conditions. Two-sided bootstrap p-values were calculated as the proportion of null resamples with absolute differences greater than or equal to the observed effect. Percentile confidence intervals were derived from the empirical distribution of independent bootstrap differences. Statistical significance was reported as ns (p > 0.05), *p ≤ 0.05, **p ≤ 0.01, ***p ≤ 0.001, and ****p ≤ 0.0001.

For in vitro measurements, photons were collected every 5 minutes, followed by fitting to Eq. 5. The mean and standard deviation were calculated from *n* = 3 independent experimental replicates.

To calculate steady-state anisotropy (*r*_ss_), the integrated intensities of the fluorescence emissions collected in the parallel (*I*_∥_) and perpendicular (*I*_⊥_) channels were used. The steady-state anisotropy was computed using the equation:

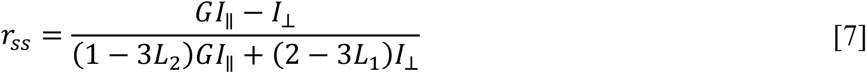

The steady-state anisotropy value provides a single, time-averaged measurement of the overall fluorescence anisotropy, which complements the dynamic information obtained from TRFA decay analysis.

### Validation of HD653 for anisotropy measurements

To assess the suitability of HD653 as a probe for TRFA, control experiments were performed using a globular protein with known rotational properties. Purified EGFP containing a single BCN at residue 39 was site-specifically labelled with either HD653 or the rigid, well-characterized anisotropy dye Cy3B. TRFA measurements were conducted on both unlabelled and labelled EGFP at 10 nM in 1× PBS. The rotational correlation times was obtained by fitting TRFA decay curves using Eq. 5, where *ρ*_slow_ is the overall correlation time of EGFP. The fitted *ρ*_slow_ was used to determine the hydrodynamic radius:

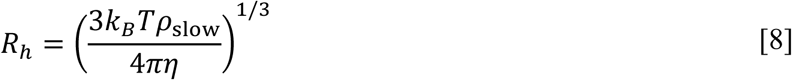

where *k*_B_ is the Boltzmann constant, *T* is the absolute temperature, and η is the solvent viscosity. Hydrodynamic radii determined for labelled EGFP were consistent with wild-type EGFP and previously reported values,^83,84^ confirming HD653 as a reliable anisotropy probe and supporting its use for segmental dynamics measurements in live-cell and in vitro assays.

### MD simulations

#### Residue-based coarse-grained model of NPC and importin-β

We adapted a previously established coarse-grained model of the NPC^36^ to incorporate importin-β proteins. In this model, each amino acid (*p*) of NPC and importin-β proteins is represented by a single bead, the nuclear envelope is modelled using fixed membrane particles (*m*), and the solvent is treated implicitly.

Protein residues are divided into three sub-categories: scaffold residues (*sc*) and FG residues (*FG*) of NPC proteins, and importin-β residues (*Imp-β*). The potential energy of the system is given by

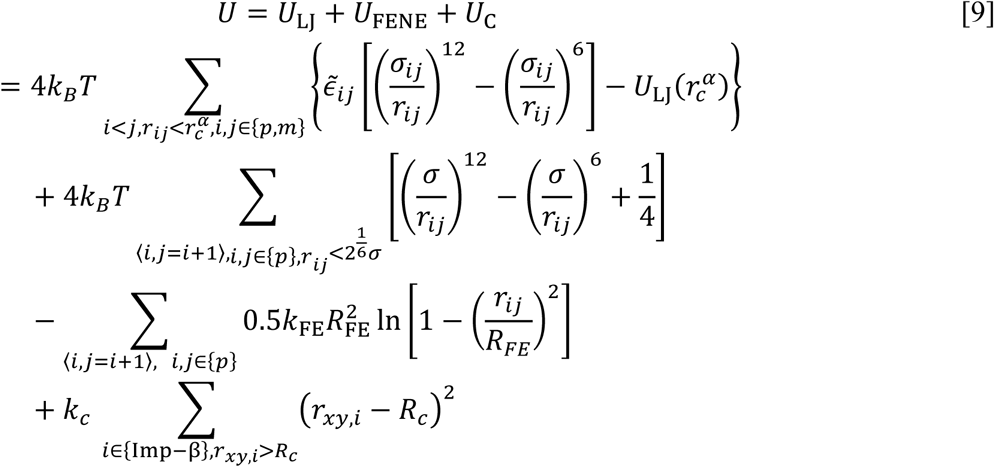

Non-bonded interactions between particles *i* and *j* are described by Lennard-Jones (LJ) potentials, whose strength ∈^∽^_i_, length *σ*_ij_, and cut-off values *r*^α^_c_ are provided in Supplementary Table 2. The LJ potentials were shifted by their values at the corresponding cut-off radii, i.e., *U*_LJ_(*r*^α^). The bond potential between adjacent beads of the FG-protein was modelled using the FENE potential^85^ with *k*_FE_ = 30*k*_B_*T* and *R*_FE_ = 1.5*σ*. The axial positions of importin-β were confined by a half-harmonic potential with stiffness *k_c_*=100*k*_B_*T*/*σ*^2^, limiting their radial distance to *r* ≲ *R*_c_ = 81*σ*≡48.6 nm.

Simulations were performed in LAMMPS (Release Sept. 2021)^86,87^. Time is reported in units of τ = *m*^2^/*k*_*B*_*T*, where *m* is the bead mass (identical for all bead types). Following earlier work^36^, we determined the reduced length scale of LJ potential obtained as *σ* ≡ 0.6 nm by matching the root-mean-square inter-residue distance (*R*_E_) of a single NUP98 FG chain (aa 1-499) in both homopolymer model and corresponding Martini representation in the weakly interacting regime.^67^ We found that for ∈^∽^_FG–FG_ = 0.5, *R*_E_ of a single NUP98 FG chain measured by single-molecule FRET, whereas ∈^∽^_FG–FG_ = 0.42 best matches in situ FLIM-FRET for NUP98 FG domain within the NPC, ^36^ reflecting in particular the effects of PTMs.

The membrane particles representing the nuclear envelope were constructed from a 100 × 100 nm^2^ coarse-grained POPC bilayer using the Python script insane.py^88,89^, with a reduced area per lipid (0.3 nm^2^) to prevent solvent permeation (command: *insane.py −l POPC -x 100 -y 100 -z 100-a 0.3 -o bilayer.gro*). A half-toroidal pore was then generated using BUMPy^88^ (command: *bumpy.py -s double_bilayer_cylinder -f bilayer.gro -z 10 -g l_cylinder:10 r_cylinder:430 r_junction:120 l_flat:1920*). To reduce computational cost, only phosphate beads were retained. NPC scaffold (*sc*) and membrane (*m*) particles were fixed throughout simulations. To prevent FG domains and importins from crossing the membrane, a LJ potential with a large repulsive range was applied (Supplementary Table 2).

We used the constricted NPC model II^36^ including FG-NUPs: NUP54, NUP58, NUP62, NUP98, POM121, NUP214, NUP153, and NUP358. Importin-β was modelled using PDBid: 3W5K (ligand Snail1 removed).^90^ Importin-β molecules were treated as rigid bodies using a rigid body integrator with a Langevin thermostat and a damping coefficient 10τ (LAMMPS command: *fix rigid langevin molecule*). FG beads were integrated using the Verlet algorithm.^91^ NUP98 FG chains were thermalised using Langevin thermostats with damping coefficient 10*τ*.^92^

#### Preparation and equilibration of importin-β inside the NPC

Importin-β molecules were added to systems of size 320*σ* × 320*σ* × 400*σ*. FG-NUPs were first equilibrated at ∈^∽^_FG–FG_ = 0.5 for 10^6^*τ*, during which they formed a surface condensate on the NPC scaffold. These compact conformations allowed us to efficiently introduce *N*_Imp-β_ =100 importin-β molecules^77^ with random initial positions and orientations around the NPC. With FG-NUPs frozen and weak interactions (∈^∽^_FG–Imp-β_ = 0.1), importin-β molecules were allowed to diffuse for 3.4 × 10^4^*τ* with thermostat damping coefficient 1000*τ*. Subsequently, the full system was equilibrated with ∈^∽^_FG–FG_ = 0.42 and ∈^∽^ _FG–Imp-β_ = 0.1 for 10^5^*τ* using a time step *δt* =10^-3^*τ* and thermostat damping coefficient 10*τ*. Production conditions were then applied by setting ∈^∽^ _FG–FG_ and ∈^∽^ _FG–Imp-β_ to their target values, followed by simulations of at least 10^6^*τ*. Three independent configurations from the equilibration phase were selected as starting points for independent production runs. Structural visualization was performed using VMD.^78^

#### Martini model of the NPC

To examine how O-glycosylation affects FG-NUP dynamics, we performed MD simulations of the human NPC using the Martini 2.2 coarse-graining scheme. We used the constricted NPC structure (PDBid: 7R5J^66^). NUPs were grouped into scaffold proteins and disordered FG-NUPs, and converted to Martini CG models using *martinize.py* script.^64,65^ FG-NUPs included NUP54, NUP58, NUP62, NUP98, POM121, NUP214, and NUP358 (Supplementary Table 3). All scaffold proteins and grafting-anchored ordered domains of FG-NUPs were restrained with a harmonic spring (1000 kJ mol^-1^ nm^-2^).

We generated O-glycosylated NPC models at three modification levels (*f*_gly_ = 0.18, 0.3 and 0.6), where *f*_gly_ denotes the fraction of O-glycosylated Ser/Thr sites (Supplementary Fig. 11). Potential O-glycosylation sites were predicted using YinOYang^93^, which assigns each residue a probability *p*∈[0,1]. Residues with *p*≥*p*_th_ were considered glycosylated. The resulting number of modified sites *N*_gly_ determined *f*_gly_=*N*_gly_/*N*_total_, where N_total_ = 58768 is the total number of Ser/Thr residues across all FG-NUPs.

Simulations were performed using GROMACS (versions 2020.6 and 2025.2)^94,95^. Interaction strengths for protein–protein, protein–sugar, and sugar–sugar interactions were rescaled by parameters *α*, *λ* and *γ*, respectively.^67,96^ The optimal *α* = 0.78 was determined by matching *R*_E_ of an untreated NUP98 FG chain to FLIM-FRET measurements (Supplementary Fig. 12) and the dense-phase density of FG condensates (Supplementary Fig. 14 and Supplementary Table 4,5). For O-glycosylated FG-NUPs, interactions involving sugars used *λ*=0.15 and *γ*=0.5, consistent with experimental osmotic second virial coefficients of aqueous sugars and protein-sugar mixtures.^68,97^

#### Equilibration of the Martini NPC systems

A cubic box (170 nm per side) was constructed to accommodate the NPC. Secondary-structure restraints were assigned using DSSP^98^. Elastic networks within ordered domains were generated using Elastic Network in Dynamics (*ElNeDyn*)^99^ (default parameters; a distance range of 0.9 nm; force constant 500 kJ mol^-1^ nm^-2^). All FG domains were modelled as coils except for locally ordered regions in NUP358 (Supplementary Table 3). The system was solvated with Martini water containing 10% antifreeze beads^100^ and neutralized with NaCl to 150 mM. The total number of solvent particles was approximately 4 × 10^7^.

Initial energy minimization used steepest descent with maximum step size 0.01 nm and force tolerance 1000 kJ mol^-1^ nm^-1^. During this step, position restraints (1000 kJ mol^-1^ nm^-2^) were temporarily removed to allow global relaxation. LJ and Coulomb potentials were first softened using the GROMACS free-energy framework (parameters: *couple-lambda0=vdw-q*; *sc-alpha*= 0.1; *sc-coul*= yes; *sc-power*= 2; *sc-sigma*= 0.3), followed by a second minimization using full potentials.

Equilibration proceeded as follows. First, NVT simulations at 310 K were performed for 100 ns (velocity-rescaling thermostat^101^, time constant 1 ps, time step *δt* = 0.015 ps) using reduced interaction scaling (*α* = *γ* = *λ* = 0.3) to relax chains into coil-like conformations. Next, NPT equilibration at 310 K and 1 bar was performed for 100 ns (velocity-rescaling thermostat^101^, semi-isotropic Berendsen barostat^102^, time constant 12 ps, compressibility 3 × 10^-4^ bar^-1^). Interaction parameters were then gradually increased. For untreated NPCs, *α* was raised from 0.3 to 0.5 and then 0.6. For O-glycosylated NPCs, *α* was increased from 0.3 to 0.6 and then 0.7, with *γ* = 0.15 and *λ* = 0.5. This stage lasted 1.5 μs. Production simulations were conducted for at least 2 μs at target *α* values, using the same thermostat and barostat parameters as in the final equilibration step and a timestep *δt* = 0.02 ps.

