## Extended Data Figures for "O-GlcNAcylation and an importin-β radial gradient keep the FG barrier liquid in live-cell nuclear pores"

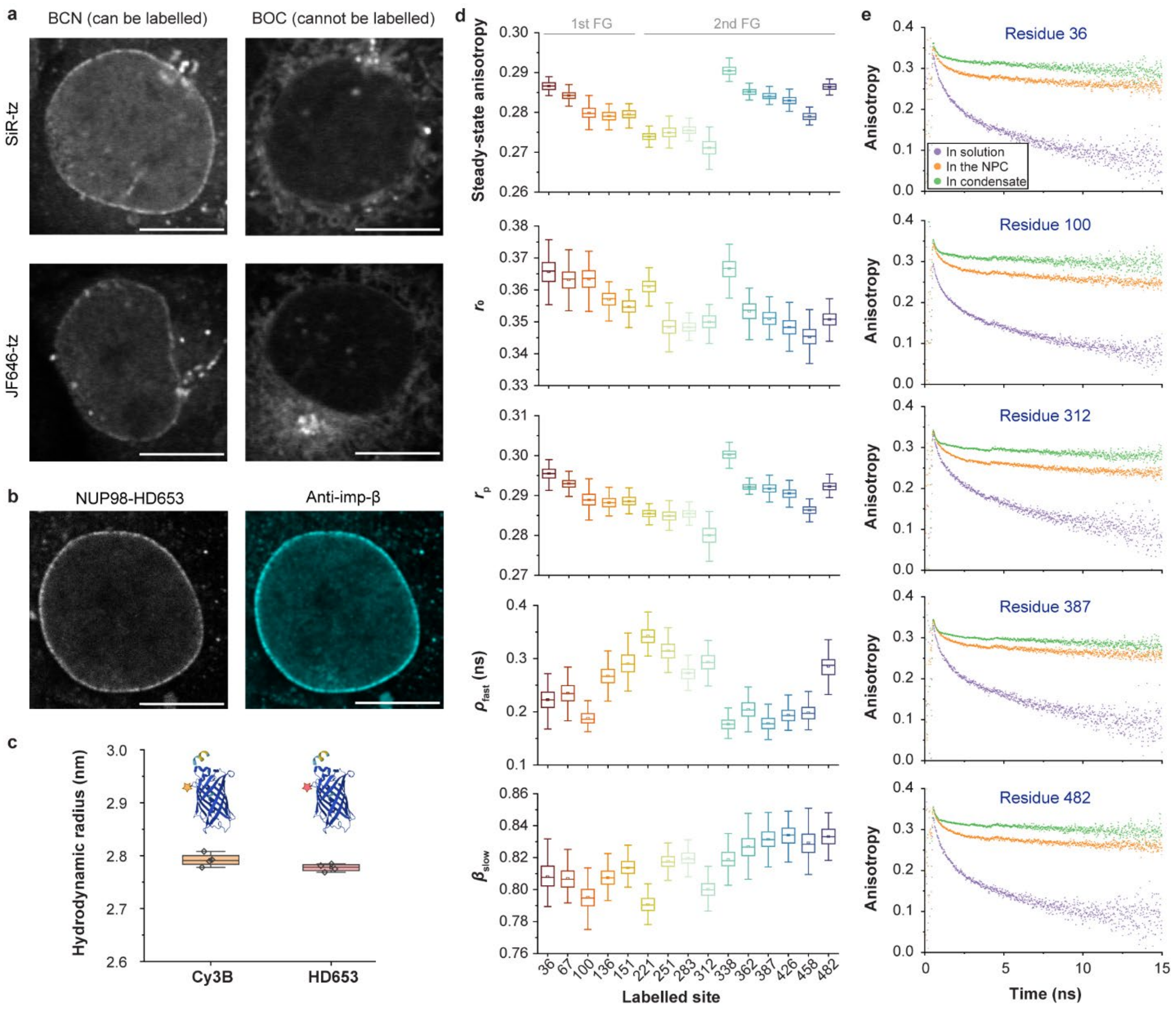


**Extended Data Fig. 1 Validation of site-specific labelling and TRFA for NUP98 dynamics in cells and in vitro. a**, COS-7 cells expressing NUP98$^{\mathrm{A221TAG}}$ were labelled with live-cell dyes (SiR-tetrazine and JF646-tetrazine) following genetic code expansion of either bicyclo[6.1.0]nonyne-lysine (BCN) or non-reactive control *t*-butyloxycarbonyl-L-lysine (BOC). Scale bars, 10 µm. **b**, Immunofluorescence images confirm that endogenous importin-β remains localized at the nuclear envelope after NUP98 labelling with HD653. Scale bars, 10 µm. **c**, Hydrodynamic radius of purified enhanced green fluorescent protein (EGFP) containing BCN at residue 39 and labelled either with HD653 or the established anisotropy dye Cy3B,[44,45] determined from TRFA measurements in solution. The measured radii match wild-type EGFP, validating HD653 as a reliable probe for fluorescence anisotropy. **d**, Summary of TRFA parameters extracted for all labelled positions across the FG domains in live cells: steady-state anisotropy, fundamental anisotropy ($r_0$), residual anisotropy ($r_p$), local rotational correlation time ($\rho_{fast}$), and relative amplitude of slow (segmental) component ($\beta_{slow}$) as defined in Eqs. 5 and 6. Box plots display the median (centre line), mean (central square), interquartile range (box), and whiskers extending to data points within 1.5×IQR. **e**, Representative anisotropy decay curves for residues 36, 100, 312, 387 and 482 measured in the NPC, in isolated FG chains in solution, and in freshly formed FG condensates.

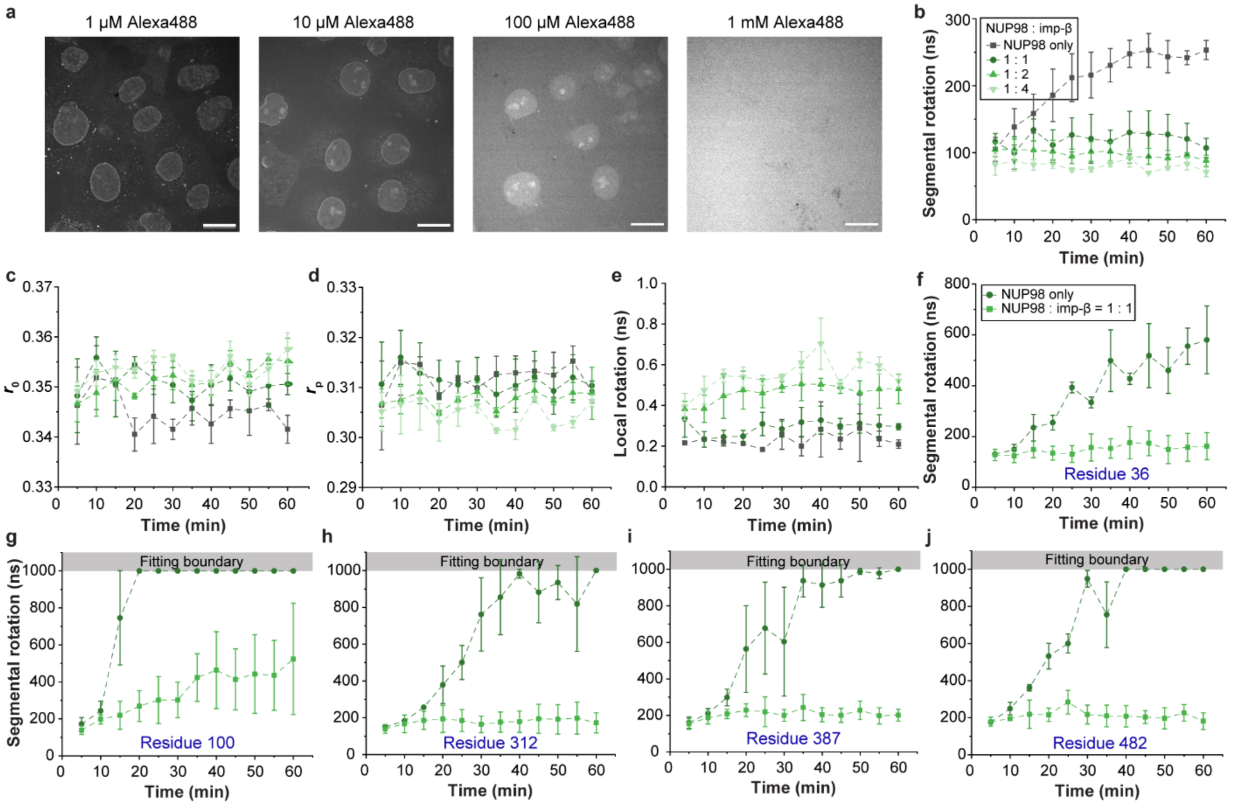


**Extended Data Fig. 2 Cellular abundance of importin-β and its modulation of NUP98 FG-domain dynamics in vitro. a,** Estimation of endogenous importin-β concentration at the nuclear pore by immunostaining with Alexa Fluor 488–labelled antibody, followed by titration with free Alexa Fluor 488 dye in solution (1 μM–1 mM). Importin-β remained detectable up to 100 μM dye but was obscured at 1 mM, indicating that the local importin-β concentration at the NPC is in the millimolar range. **b-e**, TRFA parameters of purified NUP98 FG domains co-phase separated with increasing molar ratios of importin-β: (b) segmental rotational correlation time $\rho_{\text{slow}}$, (c) fundamental anisotropy $r_0$, (d) residual anisotropy $r_p$, (e) local rotational correlation time $\rho_{\text{fast}}$. **f-j,** Segmental rotational correlation times measured across multiple labelled sites in NUP98 FG domains in the absence or presence of importin-β (1:1 molar ratio). Data represent mean ± s.d. from $n = 3$ independent replicates.

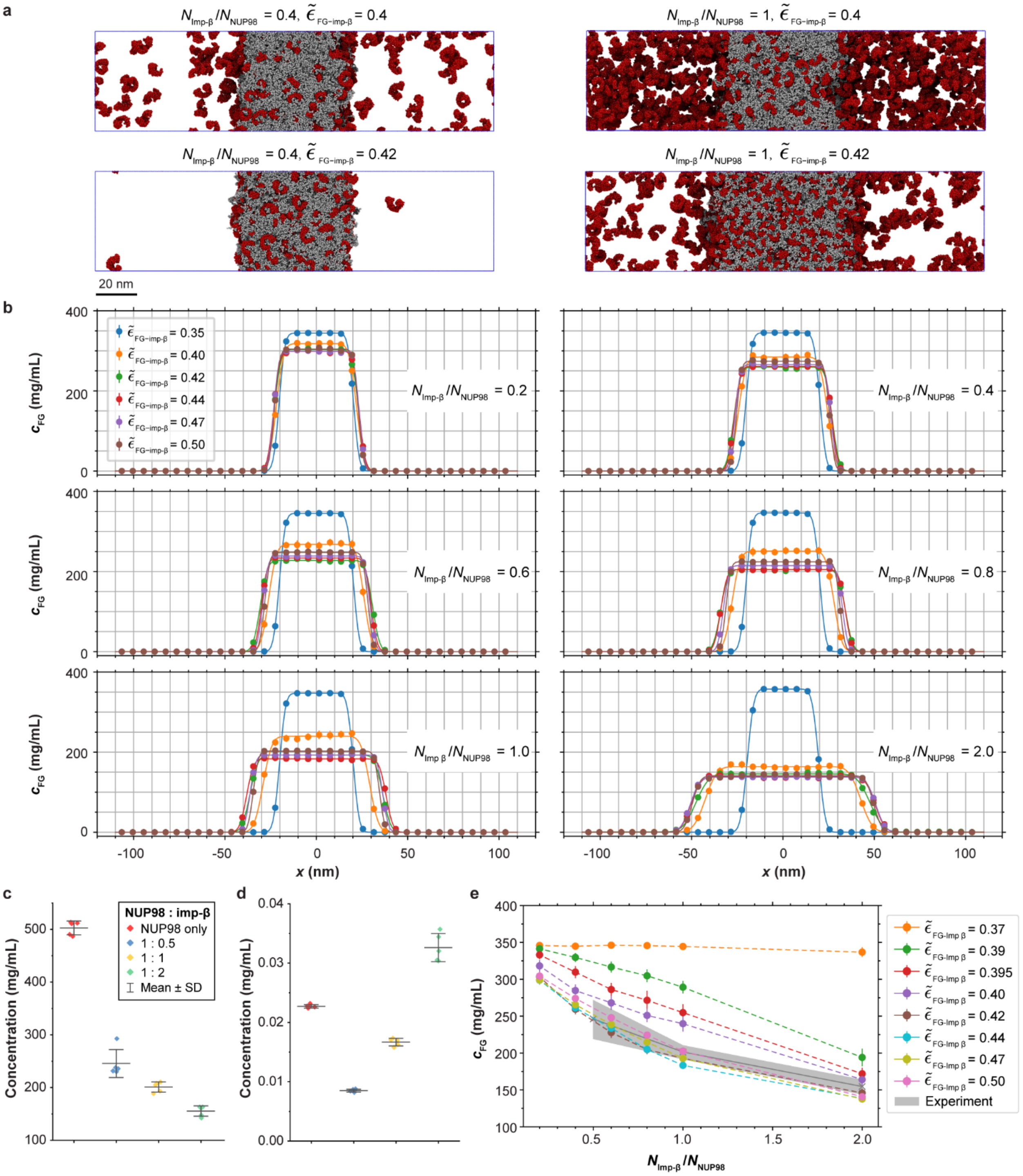


**Extended Data Fig. 3 Simulation of condensation of NUP98 FG domains with importin-β. a,** Snapshots from the end of coarse-grained MD simulations showing co-condensation of NUP98 FG domains with importin-β at varying molecular ratios ($N_{\text{Imp-β}}/N_{\text{FG}}$) and different FG–importin-β interaction strengths ($\tilde{\epsilon}_{\text{FG–Imp-β}}$). Importin-β is shown in red and NUP98 FG chains in grey. **b,** Concentration profiles of FG chains along the simulation box. The ratio of importin-β to FG chains

($N_{\text{Imp-β}}$ /$N_{\text{NUP98}}$) is indicated in each panel. The symbols and error bars represent the mean and standard error calculated over the last $45 \times 10^4\ \tau$ snapshots sampled at $10\tau$ intervals. The solid lines represent double error function fits to the concentration profiles.[36] **c, d,** Experimentally measured FG-domain concentrations in freshly formed NUP98 FG condensates co-phase separated with different molar ratios of importin-β: (**c**) FG concentration in the dense phase and (**d**) in the dilute phase from fluorescence experiments. Data represent mean ± s.d. from $n = 5$ independent replicates. Notably, in the absence of importin-β, FG chains form nanoscale assemblies that persist in solution, leading to a slightly elevated apparent concentration in the dilute phase. **e,** MD-simulated FG-chain concentrations in the dense phase of the co-condensates of NUP98 FG domains and importin-β as a function of molar ratio $N_{\text{Imp-β}}/N_{\text{NUP98}}$, comparing different FG–importin-β interaction strengths. Symbols and error bars are as in (b). The grey shading indicates the experimentally measured dense-phase concentrations from (c).

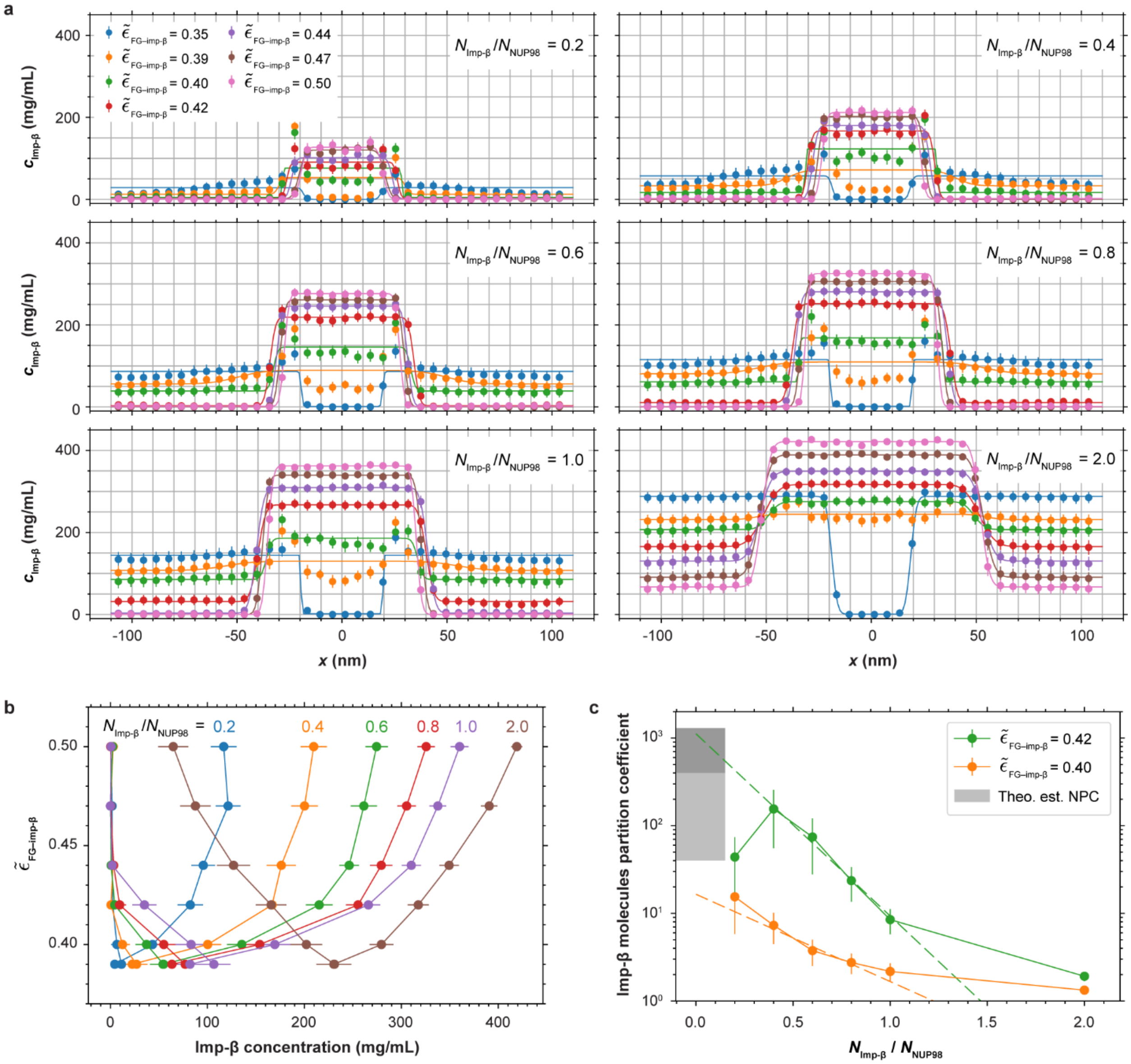


**Extended Data Fig. 4 Partitioning of importin-β into NUP98 FG condensates. a,** Concentration profiles of importin-β beads along the simulation box for different molar ratios of importin-β to NUP98 FG chains ($N_{\text{Imp-}\beta}/N_{\text{NUP98}}$). The symbols and error bars indicate the mean and standard error calculated over the last $45 \times 10^4\ \tau$ snapshots sampled every $10\tau$. The solid lines are double error function fits.[36] **b,** Phase diagram summarizing the concentrations of importin-β in the dilute and dense phases as a function of FG–importin-β interaction strength $\tilde{\epsilon}_{\text{FG–Imp-}\beta}$, obtained from the profiles in (a). **c,** The partition coefficient of importin-β ($c^{\text{dense}}_{\text{Imp}-\beta}/c^{\text{dilute}}_{\text{Imp}-\beta}$) plotted as a function of molar ratio $N_{\text{Imp-}\beta}/N_{\text{NUP98}}$ for $\tilde{\epsilon}_{\text{FG–Imp-}\beta} = 0.4$ and 0.42. The dashed lines are fits of $ae^{\frac{bN_{\text{Imp}-\beta}}{N_{\text{NUP98}}}}$ to the data points within the range $0.5 \leq N_{\text{Imp-}\beta}/N_{\text{NUP98}} \leq 1$. Extrapolation to the infinitely dilute limit (i.e., $N_{\text{Imp-}\beta}/N_{\text{NUP98}} \rightarrow 0$) yields partition coefficients of 16.5 ± 6 for $\tilde{\epsilon}_{\text{FG–Imp-}\beta} = 0.4$ and 1120 ± 364 for $\tilde{\epsilon}_{\text{FG–Imp-}\beta} = 0.42$. Proteomics data[77] estimate the partition coefficient of importins inside NPC to line between 400–1300 (dark grey rectangle; see Supplementary Information). To account for possible strong crowding effects in the NPC, compared to the cytosol/nucleus, the uncertainty band is extended downward by an additional factor of ten (light grey rectangle). For simulations with very low importin-β copy numbers, larger numerical uncertainties are expected, accounting for the observed outlier.

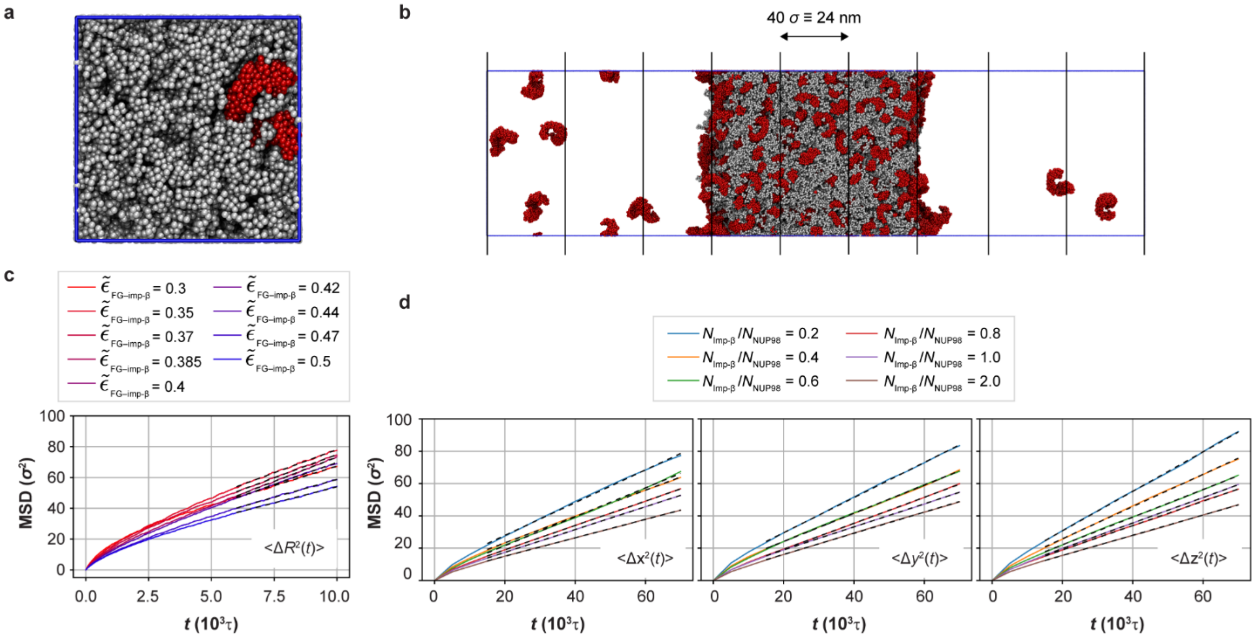


**Extended Data Fig. 5 Diffusion of importin-β inside NUP98 FG condensates. a,** Snapshot of a simulation containing a single importin-β molecule (red) embedded in an NUP98 FG condensate (grey) uniformly filling a cubic box of width 38$\sigma$. **b**, Simulation for multiple importin-β molecules partitioned into an FG condensate confined to a slab geometry (360$\sigma$ × 90$\sigma$ × 90). For mean-squared-displacement (MSD) analysis, the box was divided into nine bins of width 40$\sigma$ along the $x$-axis (vertical black lines). **c,** MSD of the centre of mass of a single importin-β, $\langle \Delta R^2(t) \rangle$, for different FG–importin-β interaction strengths $\tilde{\epsilon}_{\mathrm{FG-Imp\text{-}\beta}}$. **d,** Averaged MSD of multiple importin-β molecules located near the centre of the condensate slab (bin range $-20\sigma < x < 20\sigma$), measured at 48 × $10^4\tau$. The interaction strengths were $\tilde{\epsilon}_{\mathrm{FG-Imp\text{-}\beta}} = 0.42$ and $\tilde{\epsilon}_{\mathrm{FG-FG}} = 0.5$. MSD components along the $x$-axis (left), $y$-axis (middle) and $z$-axis (right) are shown. The black dashed lines in (c) and (d) indicate linear fits to the MSD curves: for (c), fits were applied for $t > 6 \times 10^3\tau$; for (d), fits were applied for $t > 15 \times 10^3\tau$.

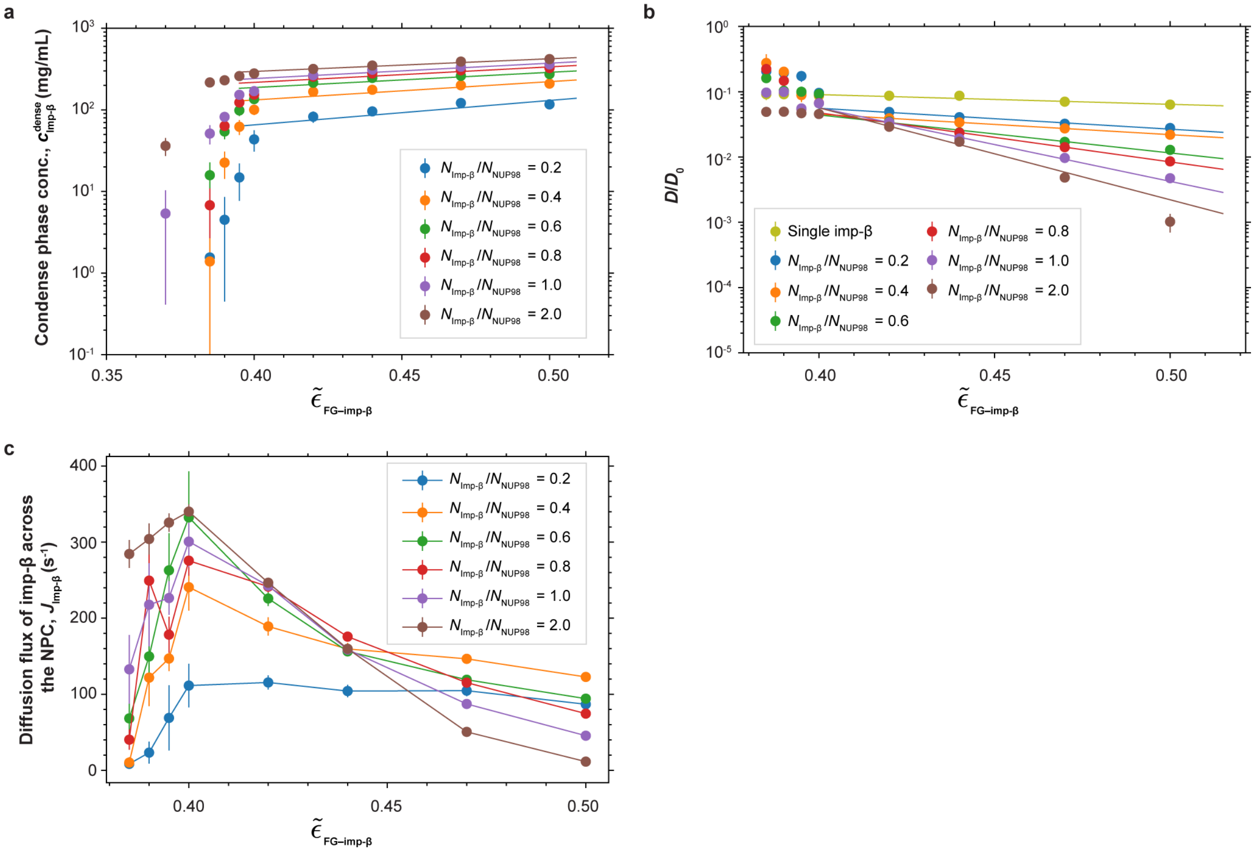


**Extended Data Fig. 6 Diffusive flux of importin-β inside NUP98 FG condensate. a,** Concentration of importin-β in the dense phase of NUP98 FG condensates. Symbols represent the importin-β concentration at the centre of the condensate slab, $c_{\text{Imp-β}}$ ($x = 0$); the error bars indicate the standard error of the mean (SEM). The solid lines show fits of the function $ae^{b\tilde{\epsilon}_{\text{FG-Impβ}}}$ to data point within the range $0.42 \leq \tilde{\epsilon}_{\text{FG–Imp-β}} \leq 0.5$. **b,** Translational diffusion coefficients ($D$) of importin-β inside FG condensates, rescaled by the diffusion coefficient of a single importin-β in solution, $D_0$ (see Supplementary Fig. 3). For molar ratios $N_{\text{Imp-β}}/N_{\text{NUP98}} = 0.2$ to 2, the symbols represent diffusion coefficients of importin-β molecules located in the centre bin of the condensate slab (see Extended Data Fig. 5b). Diffusion was averaged over the $y$- and $z$-directions, $D=(D_y+D_z)/2$, i.e., directions parallel to the condensate interfaces. The error bars indicate SEM across all importin-β molecules. The solid lines show fits of $a'e^{b'\tilde{\epsilon}_{\text{FG–Imp β}}}$ to the data points within $0.42 \leq \tilde{\epsilon}_{\text{FG–Imp-β}} \leq 0$. **c,** Estimated diffusion flux of importin-β across the NPC $J_{\text{Imp β}} = Ac^{\text{dense}}_{\text{Imp}} \times D_{\text{Imp β}}$. For reference, we used $D_0 = 51\ \mu m^2/s$ as the translational diffusion coefficient of free importin-β in solution.[103] The NPC was approximated as a cylinder of radius $R_{\text{NPC}} = 0.05\ \mu m$, height $H_{\text{NPC}} = 0.1$

μm and volume $V_{\mathrm{NPC}} = \pi R_{\mathrm{NPC}}^2 H_{\mathrm{NPC}}$. A geometric prefactor $V_{\mathrm{NPC}}/H_{\mathrm{NPC}}^2$ rescales the condensate-derived flux to the NPC geometry, neglecting effects from non-uniform spatial distributions.

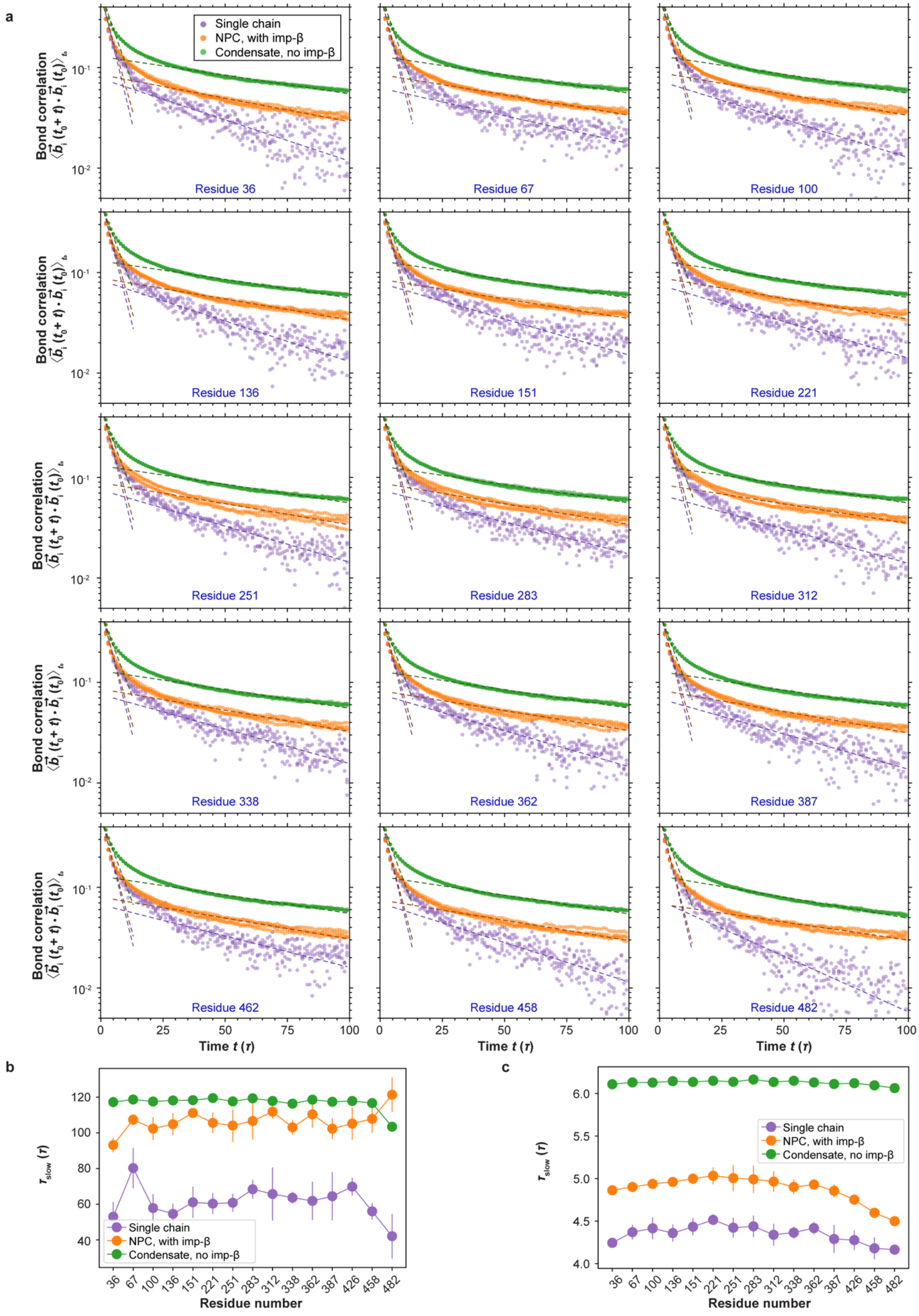
a
Single chain
NPC, with imp-β
Condensate, no imp-β
Bond correlation
Residue 36
Residue 67
Residue 100
Residue 136
Residue 151
Residue 221
Residue 251
Residue 283
Residue 312
Residue 338
Residue 362
Residue 387
Residue 462
Residue 458
Residue 482
Time t (τ)
b
c
τslow (τ)
Residue number

**Extended Data Fig. 7 Segmental rotations of NUP98 FG chains in solution, condensates and NPC. a,** Auto-correlation functions of bond vector $\langle \vec{b_i}(t_0 + t) \cdot \vec{b_i}(t_0) \rangle_{t_0}$ for NUP98 FG domain at residues 36, 67, 100, 136, 151, 221, 251, 283, 312, 338, 362, 387, 426, 458, 482. Here, $\vec{b_i}$ is the unit bond vector connecting residue $i$ to $i + 1$. The black dashed lines show exponential fits of the form $a\exp(-t/\tau_{\text{fast}})$ and $a'\exp(-t/\tau_{\text{slow}})$ to the early $[0,10\tau]$ and late $[25\tau,100\tau]$ decay regimes, respectively, providing a coarse approximation of the multi-exponential polymer dynamics. Points represent data from four independent simulation runs. Single-chain simulations were run for $10^5\tau$; condensate and NPC simulations for $10^4\tau$. Residue coordinates of the NUP98 FG domain were sampled every $\tau$. The interaction strengths were $\tilde{\epsilon}_{\text{FG–FG}} = \tilde{\epsilon}_{\text{FG–Imp-β}} = 0.42$ for NPC simulations, and $\tilde{\epsilon}_{\text{FG–FG}} = 0.5$ for single-chain and condensate simulations. **b,** Slow relaxation times, $\tau_{\text{slow}}$, obtained from fits to the late-time decay of the bond-vector autocorrelation functions. **c,** Fast relaxation times, $\tau_{\text{fast}}$. obtained from fits to the initial decay. The symbols denote mean values, and error bars are the SEM across the four independent runs.

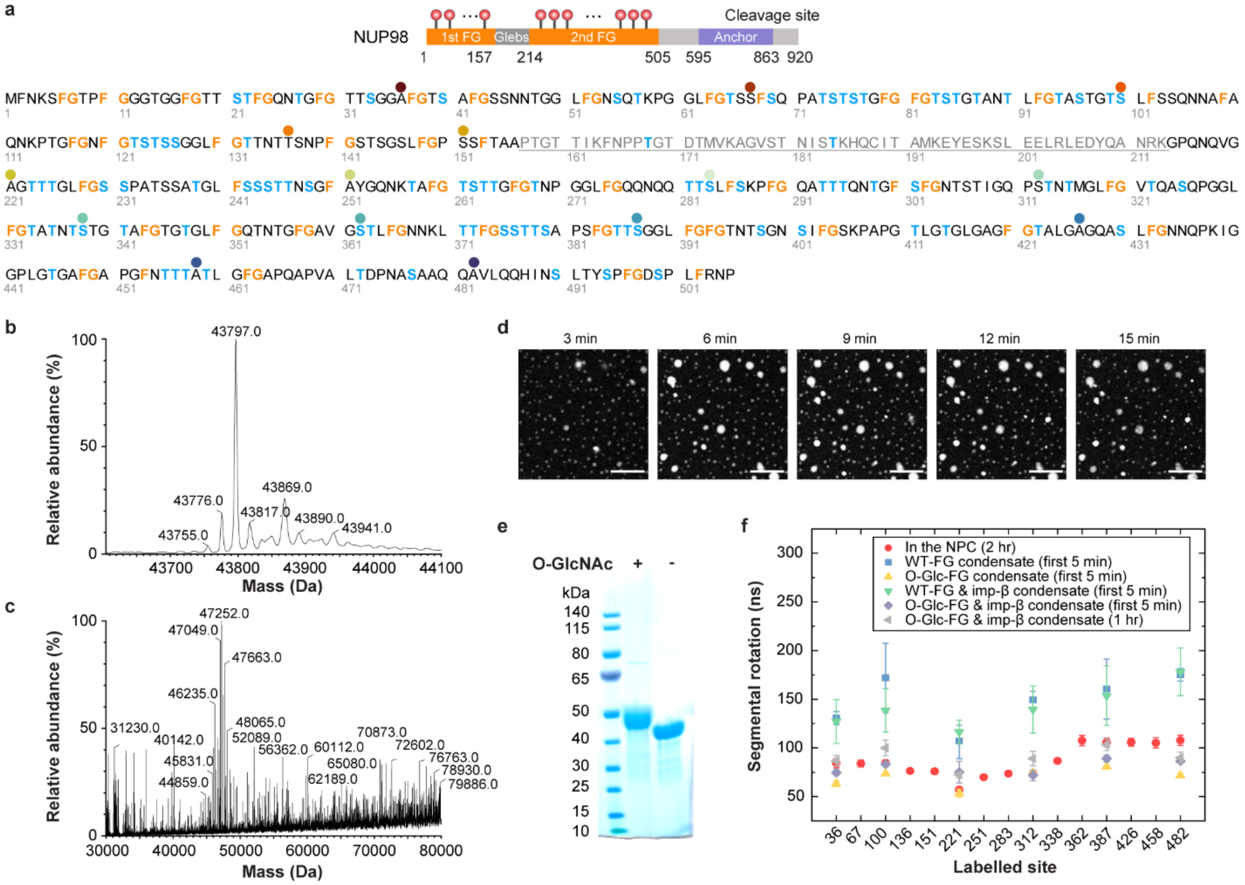


**Extended Data Fig. 8 Characterisation of O-GlcNAcylated NUP98 FG domains. a**, Predicted O-GlcNAc modification sites (blue) within the second FG domain of NUP98, annotated using the O-GlcNAc Database v2.0 (https://www.oglcnac.mcw.edu/). **b–c,** Mass spectrometry analysis of (b) wild-type and (c) enzymatically O-GlcNAcylated NUP98 FG domains. Each O-GlcNAc modification produces a mass increase of +203.08 Da. The dominant peak in (c) corresponds to approximately 17 GlcNAc moieties per polypeptide chain. **d**, Time-lapse fluorescence images showing phase separation of purified enzymatically O-GlcNAcylated NUP98 FG domains at 10 μM in vitro. Scale bar, 10 μm. **e**, Coomassie-stained SDS-PAGE gel showing that glycosylated NUP98 FG domains migrate more slowly than wild-type due to their increased molecular mass. **f**, Comparison of segmental rotational dynamics across multiple labelled sites in live-cell NPC versus in vitro condensates under the indicated conditions. Phase separation of O-GlcNAcylated FG domains with importin-β closely recapitulates the nanosecond-scale dynamics observed in the NPC.

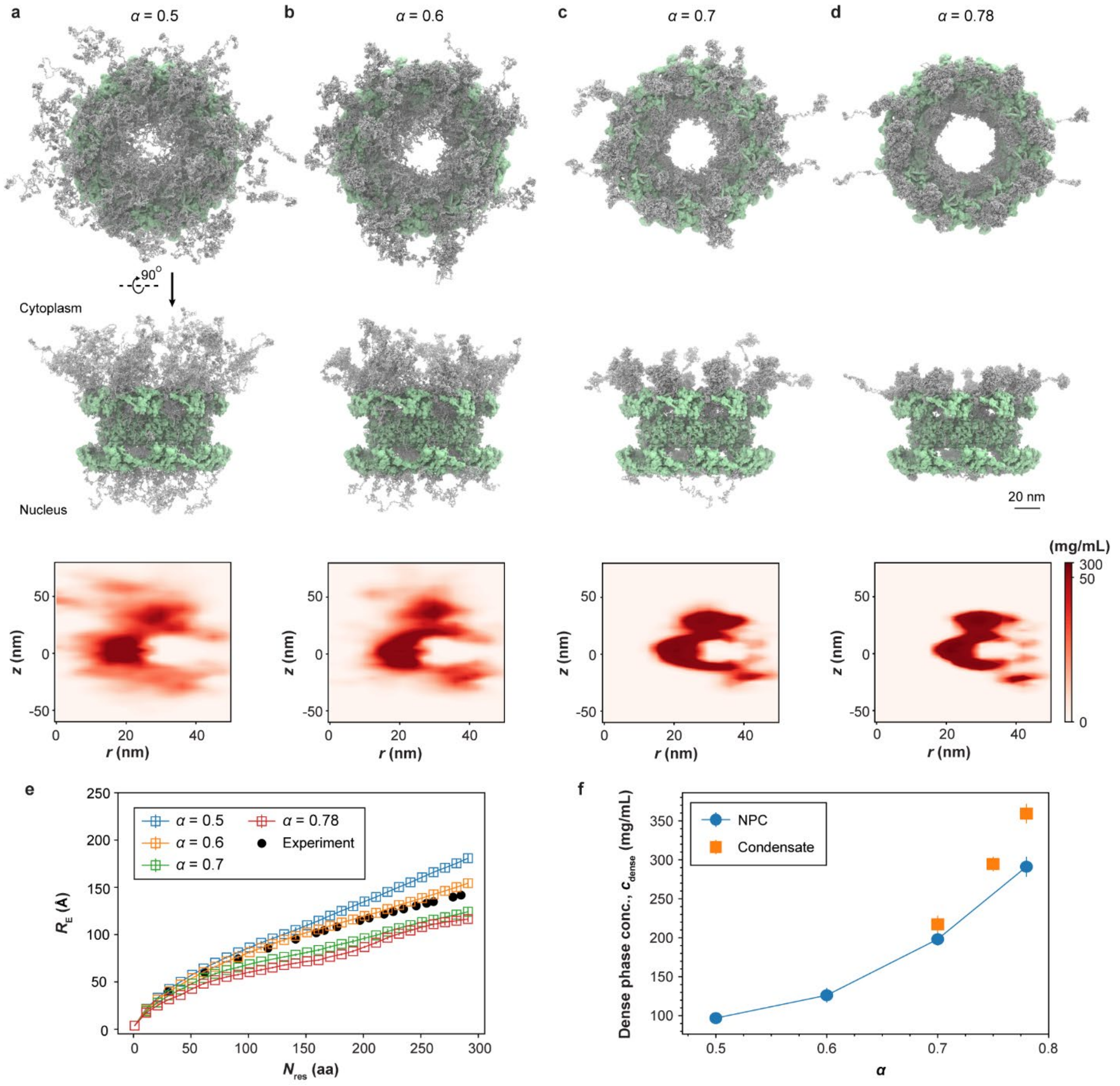


**Extended Data Fig. 9 O-GlcNAcylation places FG-NUPs at critical point to condensate formation in Martini MD simulations.** Shown are the results of Martini MD simulations of unglycosylated NPCs for different protein-protein interaction strengths $\alpha$. **a-d,** Martini MD simulation snapshots of unglycosylated NPCs at increasing levels of protein-protein interaction-scaling factor $\alpha$ applied to FG-NUPs (NUP54, NUP58, NUP62, NUP98, NUP214, NUP358, and POM121: (**a**) $\alpha = 0.5$, (**b**) $\alpha = 0.6$, (**c**) $\alpha = 0.7$, and (**d**) $\alpha = 0.78$. Top and middle rows: Top and side views of the final configurations (FG-NUPs: white, NPC scaffold: green). Bottom row: axially averaged FG-NUPs concentration (excluding FG-NUP358) over the last 600 ns of simulation (colour scale: linear 0–50 mg/mL; >50 mg/mL shown as uniform dark red; bin size $\Delta r = \Delta z = 3$

nm). Simulation in (a-d) were run for 2.5 μs. **e,** Root-mean-square inter-residue distance ($R_E$) of NUP98 FG chains in the NPC. The symbols represent mean values; error bars denote SEM computed from four non-overlapping 150-ns blocks. Black circles represent experimental in situ FLIM–FRET measurements[36], which are best matched at $\alpha \approx 0.6$, just above the critical point at $\alpha_c \approx 0.55$ for phase separation of FG-NUP98 (Supplementary Fig. 14). **f,** Dense phase concentration ($c_{\mathrm{dens}}$) of the FG-NUP network as function of $\alpha$ for NPC simulations (blue circles) and NUP98 condensates (orange squares). For NPC, $c_{\mathrm{dens}}$ corresponds to the maximum FG concentration in the protein-density maps (bottom panels of (a–d) measured near the inner ring at $r = 21$ nm, $z = 2.6$ nm (mean ± s.d.). For NUP98 condensates, $c_{\mathrm{dens}}$ was taken as the concentration at the centre of the condensate slab ($x = 0$) (mean ± s.d.).
