## Supplementary information for "O-GlcNAcylation and an importin-β radial gradient keep the FG barrier liquid in live-cell nuclear pores"

M. Yu *et al.*

mailto:

**This PDF file includes:**

Supplementary Text
Supplementary Figs. 1 to 15
Supplementary Tables 1 to 6
Captions for Supplementary Movies 1 to 9
Supplementary References (103-116)

**Supplementary text**

Concentration of NUP98 FG domain and partition coefficient of importin-β

To determine the optimum effective interaction strength between NUP98 FG and importin-β, $\tilde{\epsilon}_{\text{FG–Imp-}\beta}$, we built systems containing 500 NUP98 FG chains (aa 1-499; 499 monomers per chain) and 100, 200, 300, 400, 500, or 1000 importin-β molecules. This corresponds to molecular ratios $N_{\text{Imp-}\beta}/N_{\text{NUP98}}$ = 0.2, 0.4, 0.6, 0.8, 1, and 2. The simulation box size was $360\sigma\times90\sigma\times90\sigma$.

We first equilibrated a NUP98 FG condensate at $\tilde{\epsilon}_{\text{FG–FG}} = 0.5$ for $3\times10^4\tau$. We then froze the FG condensate (i.e., the equations of motion for FG beads were not integrated) and equilibrated the importin-β distribution for another $3\times10^4\tau$ at a weak cross-interaction strength $\tilde{\epsilon}_{\text{FG–Imp-}\beta} = 0.2$, using a time step $\delta t = 10^{-2}\tau$. This procedure allowed the importin-β proteins to rapidly diffuse through the simulation box and uniformly coat the FG condensate interfaces. Finally, we increased $\tilde{\epsilon}_{\text{FG–Imp-}\beta}$ to its target value. The total simulation time was at least $1.8\times10^6\tau$, with a time step was $0.01\tau$.

We computed the concentration profiles of importin-β particles inside the slab by time-averaging over the last $3\times10^4\tau$, sampling configuration every $10\tau$. To obtain concentration profiles, we uniformly distributed the total mass of each protein over its beads. The total masses of the FG chain (aa 1–499) and importin-β are 49,158 g/mol and 90,266 g/mol, respectively. Thus, the mass per bead of NUP98 FG and importin-β is 98.5 g/mol and 103 g/mol, respectively. The simulation box was discretized into bins of size 3 nm, and bead concentrations in each bin were computed and time-averaged.

For $\tilde{\epsilon}_{\text{FG–Imp-}\beta} < 0.3$, importin-β rarely penetrated the condensate, whereas for $\tilde{\epsilon}_{\text{FG–Imp-}\beta} > 0.3$, importin-β mostly partitioned into the condensate. We fitted the concentrations of FG monomers and importin-β particles using double error functions

$$c_\alpha(x) = A_\alpha[\text{erf}(x + B_\alpha)/w_\alpha) - \text{erf}(x - B_\alpha)/w_\alpha)] + c_\alpha^{\text{dilute}} \qquad [10]$$

where $\alpha\in\{\text{Imp-}\beta, \text{FG}\}$, and $A_\alpha = \left(c_\alpha^{\text{dense}} - c_\alpha^{\text{dilute}}\right)/2\,\text{erf}(B_\alpha/w_\alpha)$. Here, $B_\alpha$ and $w_\alpha$ are the position and width of the interface between dense and dilute phases, respectively. The solid lines in Extended Data Fig. 3 and 4 show the corresponding fits.

For the dense-phase concentrations of NUP98 FG domain at $\tilde{\epsilon}_{\text{FG–Imp-}\beta} = 0.42$, we could match the experimental dense-phase concentrations of NUP98 FG at different importin-β to FG ratios (Extended Data Fig. 3c). The importin-β concentrations in the dense and dilute phases at different $\tilde{\epsilon}_{\text{FG–Imp-}\beta}$ are shown in Extended Data Fig. 4b. For all $N_{\text{Imp-}\beta}/N_{\text{NUP98}}$, the two branches of dilute and dense phases meet in the window $0.39 \leq \tilde{\epsilon}_{\text{FG–Imp-}\beta} \leq 0.4$, in which the importin-β molecules can freely diffuse within the FG condensate, as evidenced by nearly flat concentration profiles in Extended Data Fig. 4a.

Partition coefficient of importin-β into NUP98 FG condensate and the NPC

The partition coefficient (PC) of importin-β into the FG condensate was computed as the ratio of the importin-β bead concentration in the dense and dilute phases, i.e., $c_{\mathrm{Imp}\,\beta}^{\mathrm{dense}}/c_{\mathrm{Imp}\,\beta}^{\mathrm{dilute}}$ (Extended Data Fig. 4c).[104]

We next estimated the partition coefficient for importin-β binding into the NPC pore. U87-MG cells contain approximately $N_{\mathrm{Imp}\,\beta}^{\mathrm{cell}} = 10^6$ importin-β molecules in total.[105] For a cell diameter $D_{\mathrm{cell}}$ = 13 μm and a volume $V_{\mathrm{cell}} = \pi D_{\mathrm{cell}}^3/6$, the number density of importin-β is approximately

$$\rho_{\mathrm{Imp}}^{0} = \frac{N_{\mathrm{Imp}\,\beta}^{\mathrm{cell}}}{V_{\mathrm{cell}}} = 8.7 \times 10^{20}\ \mathrm{m}^{-3} \qquad [11]$$

Representing the NPC FG-network as a cylinder of radius $R_{\mathrm{NPC}}$ = 30 nm and height $H_{\mathrm{NPC}}$ = 60–100 nm (spanning from the NPC scaffold to the tip of the FG-NUP358 extension on the cytoplasmic side), the NPC FG-network volume lies in the range $V_{\mathrm{NPC}} = \pi R_{\mathrm{NPC}}^2 H_{\mathrm{NPC}} = 8.48\times10^{-23}$ to $2.827\times10^{-22}$ $\mathrm{m}^3$. Thus, the expected number of importins in an NPC without binding is $n_{\mathrm{Imp}}^{0} = \rho_{\mathrm{Imp}}^{0} V_{\mathrm{NPC}}$ = 0.073–0.245.

Given that there are about 100 importin-β molecules per NPC,[106] the importin partition coefficient into NPCs is therefore, $\mathrm{PC} = 100/n_{\mathrm{Imp}}^{0} \approx$ 400 to 1300, as marked by the grey region in Extended Data Fig. 4c.

For the importin PC into the FG condensate, we fitted the function $ae^{\frac{bN_{\mathrm{Imp}\,\beta}}{N_{\mathrm{NUP98}}}}$ in the range $0.5 \leq N_{\mathrm{Imp}\text{-}\beta}/N_{\mathrm{NUP98}} \leq 1$, where importin-β is present in both dilute and dense phases. To estimate the importin-β PC in the infinitely dilute limit, we evaluated the fitted function at $N_{\mathrm{Imp}\text{-}\beta}/N_{\mathrm{NUP98}} = 0$. In this way, we obtained PC = 16.5 ± 6 for $\tilde{\epsilon}_{\mathrm{FG-Imp}\text{-}\beta} = 0.4$ and PC = 1120 ± 364 for $\tilde{\epsilon}_{\mathrm{FG-Imp}\text{-}\beta} = 0.42$. The uncertainty of the calculated PC corresponds to one standard deviation of the parameter $a$ , as obtained from the least-square fits. Thus, we consistently found that for $\tilde{\epsilon}_{\mathrm{FG-Imp}\text{-}\beta} = 0.42$ can reproduce the estimated importin-β partition coefficient in the NPC.

The above procedure is subject to several uncertainties, including NPC volume estimates and importin copy numbers. To account for potential strong crowding effects in the NPC relative to cytosol and nucleus, we further extended the uncertainty band downward by a factor of ten.

Dilution of aggregation-prone FG-NUPs by importin-β

We calculated the relative density of the aggregation-prone segment aa 85-124 with respect to the pure FG condensate, $n_{\mathrm{aa85\text{-}124}}$ [12] by placing a spherical probe of 15 nm radius at the centre of the condensates, avoiding interfacial regions of the slab-shaped condensate. Trajectories were sampled every $500\tau$ over at least $1.5\times10^5\tau$. For each sampled frame, we counted the number $N$ of chains having at least one monomer from segment aa 85–124 inside the probe. In a pure FG condensate, the average number of such chains was $N_0 = 23$ chains, which we used as a normalizer to compute the relative density $n_{\text{aa85-124}} = N/N_0$.

Translational diffusion coefficient of importin-β inside the NUP98 FG condensate

As a reference and consistency check, we first computed the translational diffusion coefficient of a single importin-β protein free in implicit solvent. The mean-squared displacement (MSD) of the importin-β center of mass (COM) was computed as $\langle \Delta\vec{R}(t)^2 \rangle = \langle \left(\vec{R}(t+t_0) - \vec{R}(t_0)\right)^2 \rangle_{t_0}$, where $\vec{R}(t) = (x(t), y(t), z(t))$ is the vector position of the COM at time $t$, and $t_0$ is the reference time for computing the ensemble averaging. The total simulation time was $10^6\tau$, and the COM was sampled every $\tau$.

In Supplementary Fig. 3 we show the MSD of a single importin-β protein for Langevin thermostat damping coefficients $\tau_M = 0.1\tau$, $\tau$ and $10\tau$. The implicit solvent friction coefficient is obtained as $\gamma = M/\tau_M$, where $M = 846m$ is the mass of importin-β. The dashed black lines are the MSD of a 3D Brownian particle with mass $M$ immersed in a solvent with the friction coefficient $\gamma$,[107]

$$\langle \Delta\vec{R}(t)^2 \rangle = 6Dt + \frac{6k_BT}{M}\tau_M^2\left[e^{-t/\tau_m} - 1\right] \qquad [12]$$

where $\tau_M = M/\gamma$ is the crossover time from ballistic to diffusive motion. There is an excellent agreement between the simulation and theory. For $\tau_M = 10\tau$, the translational diffusion coefficient is obtained from the fluctuation-dissipation theorem, $D_0 = (k_BT)/\gamma = 0.0011\sigma^2/\tau$. We used $D_0$ to rescale the translational diffusion of importins inside the FG condensate.

The translational diffusion coefficient of importin-β proteins inside NUP98 FG condensates was computed with two simulation setups. In the first setup, a single importin-β was immersed inside a cubic box of size $38\sigma \equiv 22.8$ nm with 44 NUP98 FG domain (aa 1-499) chains (Extended Data Fig. 5a). The resulting concentration of NUP98 FG domain is 303 mg/mL. To obtain the translational diffusion coefficients, we fitted a linear function, $\text{MSD}(t) = 6Dt + b$, to the MSD of importin COM for time $t > 6\times10^3\tau$ (Extended Data Fig. 5c). The total simulation time for each setup was at least $2\times10^5\tau$, and the importin-β COMs were recorded every $\tau$.

In the second setup, we computed the diffusion coefficient inside a slab-like condensate (Extended Data Fig. 5b). Here, the simulation box was partitioned along the $x$-axis into bins of size $\Delta x = 40\sigma = 24$ nm. For each importin-β molecule, we computed the MSD separately in the $x$, $y$, and $z$ directions and then averaged the MSDs of importin-β molecules in the same bin at time $48\times10^5\tau$ (Extended Data Fig. 5d). We fitted the MSD curves using $\text{MSD}_\alpha(t) = 2D_\alpha t + b$ to extract diffusion coefficients $D_\alpha$ for $\alpha = \{x, y, z\}$. At the centre of the slab, diffusion was nearly isotropic, $D_x \approx D_y \approx D_z$ (Extended Data Fig. 5d).

Rate of importin-mediated cargo transport across NPC matches experimental estimates

We estimated the rate of transport across individual NPCs by calculating the diffusive flux of importin-β. In this minimal model, each importin-β is assumed to be loaded with cargo at one face of the NPC and to release its cargo at the opposite face. For diffusion through a cylindrical pore of height $H_{\mathrm{NPC}}$, the mean first passage time is $t_{\mathrm{MFPT}} = H_{\mathrm{NPC}}^2/2D$, where $D$ is the diffusion coefficient of the importins. The translational diffusion coefficient of a single importin-β in solution is $D_0 = 51\ \mu\mathrm{m}^2/\mathrm{s}$,[103] and the NPC pore height is $H_{\mathrm{NPC}} \approx 0.1\ \mu\mathrm{m}$. The mean passage time is therefore $t_{\mathrm{MFPT}} = H_{\mathrm{NPC}}^2/2D = (D_0/10D)$ ms. In our simulation model, the ratio of diffusion coefficients in the condensate and in solution is $D_0/D \approx 10$–$100$, yielding passage times $t_{\mathrm{MFPT}} \approx 1$ –10 ms, in line with experimental estimates.[34,56,57]

The affinity of importin-β for NUP98 FG is tuned to maximize NPC cargo exchange

The total rate of cargo transport across an NPC is given by the ratio of the number of importins in its pore ($n_{\mathrm{Imp\text{-}\beta}}$) and the mean time ($t_{\mathrm{MFPT}}$), $J_{\mathrm{Imp\text{-}\beta}} = n_{\mathrm{Imp\text{-}\beta}} / t_{\mathrm{MFPT}}$. This is analogous to the product of charge carrier density and mobility that determines conductance in a semiconducting material. With $n_{\mathrm{Imp-\beta}} \approx V_{\mathrm{NPC}} c_{\mathrm{Imp-\beta}}^{\mathrm{dense}}$, where $V_{\mathrm{NPC}} = \pi R_{\mathrm{NPC}}^2 H_{\mathrm{NPC}}$ is the volume of an NPC pore (with radius $R_{\mathrm{NPC}} = 0.05\ \mu\mathrm{m}$) and $c_{\mathrm{Imp\ \beta}}^{\mathrm{dense}}$ is the dense-phase importin concentration, we obtain $J_{\mathrm{Imp-\beta}} = 2D\ c_{\mathrm{Imp-\beta}}^{\mathrm{dense}} \pi R_{\mathrm{NPC}}^2 / H_{\mathrm{NPC}}$. Extended Data Fig. 6a,b shows that the concentration of importins in an FG condensate increases with the cross-interaction strength $\tilde{\epsilon}_{\mathrm{FG–Imp\text{-}\beta}}$, while the apparent diffusion coefficient decreases. As a result, the product $D\ c_{\mathrm{Imp-\beta}}^{\mathrm{dense}}$ (and hence $J_{\mathrm{Imp\text{-}\beta}}$) reaches its maximum near $\tilde{\epsilon}_{\mathrm{FG–Imp\text{-}\beta}} = 0.4$, close to the interaction strength of 0.42 that we obtained by matching the partition coefficient. Thus, the interaction between FG-NUPs and importins appears to be tuned to maximize the cargo transport by balancing binding affinity (which increases the number of importins in the pore) against their mobility (which drops when the binding becomes too strong).

Moreover, the maximum in $J_{\mathrm{Imp\text{-}\beta}}$ occurs near the window $0.39 \leq \tilde{\epsilon}_{\mathrm{FG–Imp\text{-}\beta}} \leq 0.4$, where the two branches of dilute and dense phases of importin-β meet (Extended Data Figs. 4b, 6c). At this point, we expect the barrier to diffusion across the interface between dilute and dense phases of FG-NUPs to be minimal. Hence, an interaction strength of $\tilde{\epsilon}_{\mathrm{FG–Imp\text{-}\beta}} = 0.42$ appears to be near-optimal also for cargo proteins that are loaded onto the importins outside the dense FG-NUP network.

NUP98 extension inside NPC

***Residue-to-single-bead model***. We computed the root-mean-square inter-residue distance $R_{\mathrm{E}}$ of the NUP98 FG domain within the NPC as a function of sequence separation, following the procedure described in the reference[36]. The inclusion of importin-β molecules in the NPC did not significantly affect $R_{\mathrm{E}}$ of NUP98 FG domain (Supplementary Fig. 6).

***Martini model.*** We computed $R_{\mathrm{E}}$ of the NUP98 FG domain within the NPC in the Martini model as a function of sequence separation, again following the workd by Yu et al.[36]. In this case, instead of bead positions, we used the centers of geometry of the residues.

Segmental rotation of NUP98 FG domain

***Residue-to-single-bead model.*** We calculated the auto-correlation function (ACF) of unit bond vectors $\langle \vec{b_i}(t_0 + t) \cdot \vec{b_i}(t_0) \rangle_{t_0}$ to quantify segmental rotation at residues 36, 67, 100, 136, 151, 221, 251, 283, 312, 338, 362, 387, 426, 458, 482 along the NUP98 FG domain. For chains in the NPC, we extended this set by including the residues 500, 520, 540, 560 and 580. Here, $\vec{b_i}$ is the unit bond vector connecting residue $i$ to $i + 1$.

We used the four independent configurations of a single NUP98 chain, FG–importin-β condensate, and NPC systems, and continued the simulation for an additional $10^5\tau$ (single chain) and $10^4\tau$ (condensate and NPC). The coordinates of the selected residues were sampled every $\tau$. Consistent with TRFA experiments, the ACF of single FG chains at $\tilde{\epsilon}_{FG–FG} = 0.5$ has the fastest decay, and the FG condensates have the slowest decay. As a function of position, the slow component of segmental rotation slows down toward the anchoring point on the NPC scaffold, but for the simple coarse-grained model the effect is weaker than in the TRFA experiments (Supplementary Fig. 9).

***Martini model.*** We followed a similar procedure in the Martini model. Here, $\vec{b_i}$ is the unit bond vector connecting back-bone beads of residue $i$ to $i + 1$. We used three independent configurations each for the single FG chain, FG condensate and NPC systems, and continued the simulation for another 100 ns for all cases. The coordinates of the selected residues were sampled every ps. All simulations were performed at scaling $\alpha = 0.78$, $\gamma = 0.5$ and $\lambda = 0.15$. Again, consistent with dynamic anisotropy experiments, the ACF of single FG chains decays fastest, and FG condensates exhibit the slowest decay.

Mean contact number of NUP98 FG chains with importin-β

We computed the mean contact number between residues of the 48 NUP98 FG chains in the NPC and importin-β beads by averaging the number of importin beads located within $1.5\sigma$ of each residue over the last $5{\times}10^4\tau$. The box–whisker plot in Supplementary Fig. 10 summarizes the statistics of the mean contact number for the 48 NUP98 FG chains.

Maritni simulation details of single FG-NUP98 chain

An atomistic model of a single NUP98 FG chain (aa 1-499) was generated using the PyMOL *fab command* software[108] (using the sequence of UniProt P52948-2 sequence as an input). Model was converted to a Martini 2.2 representation using *martinize.py*[64,65], assigning coiled secondary structure ('*C*') to the entire protein.

The chain was first simulated in vacuum in a cubic NVT box of size $L_x = 30$ nm at $\alpha = 1$, to form a globular structure. The globular chain was then placed in a cubic box of the same size, solvated with Martini water, and neutralized by adding Cl- ions; 10% of the water beads were replaced with antifreeze beads. Initial energy minimization was performed using the steepest-descent algorithm with a maximum step size of 0.01 nm and a force tolerance of 1000 kJ $mol^{-1}$ $nm^{-1}$.

The system was then equilibrated at weak protein–protein interaction strength $\alpha = 0.3$ via sequential NVT and NPT simulations of 100 ns each. The temperature was set to 300 K using a velocity-rescaling thermostat[101] with time constant 1 ps. The pressure was set at 1 bar using Berendsen barostat[102] with time constant 12 ps and compressibility $3\times10^{-4}$ $bar^{-1}$.

For production runs, we used the velocity rescaling thermostat[101] and Parrinello-Rahman barostat[109] with parameters identical to those in the equilibration step. The time step was $\delta t = 0.02$ ps for $\alpha \leq 0.6$ and $\delta t = 0.03$ for $\alpha > 0.6$. The total production time was at least 8 μs.

Simulation details of Martini condensates

To build condensates, 100 globular FG-NUP chains (equilibrated at strong protein-protein interaction) were positioned randomly inside a box of size $L_x = 40$ nm and $L_y = L_z = 30$ nm. Martini water was added, and the system was neutralized with $Cl^-$ ions; 10% of the water beads were replaced with antifreeze beads. After setting $\alpha = 0.3$, the system was equilibrated in the NVT ensemble for 100 ns at 300 K, using the velocity-rescaling thermostat (time step $\delta t = 0.015$ ps, time constant 1 ps). Next, 1.5 μs NPT simulation at 300 K and 1 bar was performed using the velocity-rescaling thermostat[101] and a Berendsen barostat[102] (time constant 12 ps, compressibility $3\times10^{-4}$ $bar^{-1}$).

For production runs, the velocity-rescaling thermostat[101] and Parrinello-Rahman barostat[109] were used with similar parameters as in the equilibration process. By the end of the equilibration process, the chains had relaxed into coil-like configurations and completely filled the simulation box.

To generate a slab configuration, the periodic boundary was unwrapped using the final configuration, and the equilibrated box was placed at the centre of a larger slab of size $L_x = 120$ nm and $L_y = L_z = 30$ nm. Solvent particles and 10% antifreeze water beads were then added to the remaining volume. After setting $\alpha = 0.7$, the system was equilibrated in NVT and NPT ensembles as described above, for a total equilibration time of 500 ns. Finally, the equilibrated systems were used as the initial conditions for a production run at the target $\alpha$ values, with total production times of at least 10 μs.

Simulation details of Martini NPC

A cubic simulation box of size 170 nm was configured to include the NPC structure. Elastic Network in Dynamics (*ElNeDyn*)[99] was used to generate the elastic network between the ordered domains of NUPs to maintain the tertiary structure, with default parameters (distance range 0.9 nm, force constant of 500 kJ $mol^{-1}$ $nm^{-2}$). The DSSP algorithm was used to assign secondary structure restraints.[98] All FG domains were treated as coils, except for locally ordered structures in FG-NUP358 (Supplementary Table 3). Position restraints with stiffness 1000 kJ $mol^{-1}$ $nm^{-2}$ were applied.

The system was solvated, and 10% of water beads were replaced with antifreeze beads. NaCl ions were added to ensure charge neutrality and a salt concentration of 150 mM. The total number of solvent particles was approximately $4\times10^7$.

Initial energy minimization was performed using the steepest-descent algorithm with a maximum step size of 0.01 nm and a force tolerance of 1000 kJ $mol^{-1}$ $nm^{-1}$, after which position restraints were temporarily switched off to allow relaxation. We used the free energy package of GROMACS to soften Lennard-Jones and Columbic interaction potentials (parameters: *couple-lambda0=vdw-q*; *sc-alpha*= 0.1; *sc-coul*= yes; *sc-power*= 2; *sc-sigma*= 0.3). We then repeated energy minimization with the full (unsoftened) LJ and Columbic potentials.

Subsequently, we equilibrated the system with position restraints applied. First, an NVT simulation at 310 K was performed for 100 ns using a velocity-rescaling thermostat[101] (time constant 1 ps, time step $\delta t$ = 0.015 ps). We used low values of scaling parameters ($\alpha = \gamma = \lambda = 0.3$) to allow FG chains to equilibrate in coil-like conformations. Next, an NPT simulation at 310 K and 1 bar was performed for 100 ns using a velocity-rescaling thermostat[101] and semi-isotropic Berendsen barostat[102] (time constant 12 ps, compressibility $3\times10^{-4}$ $bar^{-1}$).

We then gradually increased $\alpha$. For untreated NPCs, $\alpha$ was raised from 0.3 to 0.5 and then to 0.6. For O-glycosylated NPCs, $\alpha$ was increased from 0.3 to 0.6 and then to 0.7, while $\gamma = 0.15$ and $\lambda = 0.5$ were used. The total simulation time of this step was 1.5 μs.
For production runs, $\alpha$ was set to its target value, and simulations were performed using the velocity-rescaling thermostat[101] and semi-isotropic Berendsen barostat[102] with the same parameters as in the final equilibration step. The time step was $\delta t$ = 0.02 ps. The production runs lasted approximately 3.5 μ*s* for untreated NPC and at least 2 μ*s* for O-glycosylated NPCs.

Selection of glycosylation sites

Potential O-glycosylation acceptor sites (Ser/Thr) were predicted using the *YinOYang* server,[93] which assigns each residue a probability $p \in [0,1]$ of being modified based on protein sequence. We classified a site as glycosylated when $p \geq p_{\mathrm{th}}$, where the probability threshold $p_{\mathrm{th}}$ controls the balance between sensitivity and specificity (higher $p_{\mathrm{th}}$) yields fewer, higher-confidence sites). For a given $p_{\mathrm{th}}$, we counted the selected sites $N_{\mathrm{gly}}$ and defined the modification level as $f_{\mathrm{gly}} = N_{\mathrm{gly}}/N_{\mathrm{total}}$, where $N_{\mathrm{total}}$ is the number of Ser/Thr residues in all FG-NUPs. There are $\mathrm{N_{total}} = 58768$ residues for FG-NUPs inside the NPC. We used the following number of O-glycosylated sites for the NPC: 10480 ($f_{\mathrm{gly}} = 0.18$), 18176 ($f_{\mathrm{gly}} = 0.3$) and 34880 ($f_{\mathrm{gly}} = 0.6$). Supplementary Fig. 11 shows the distribution maps of hydrophobic and O-glycosylated residues of FG-NUPs inside the NPC.

Glycan parametrization in the Martini force field

We model O-GlcNAc in the Martini 2 model with a standard sugar ring represented by a closed triangle of three CG beads, whose bead types were assigned adapting previously published N-glycan models[97,110,111]. The 'small' bead type was chosen for the ring bead lacking a hydroxyl group at the hemiacetal position to represent a covalent linkage to the amino acid side chain. Reference atomistic simulations of O-glycosylated tripeptides (Ala–Ser–Ala and Ala–Thr–Ala) were performed following the previously described protocol.[112] After maping atom groups to corresponding Martini beads, equilibrium distributions of bond lengths, angles, and dihedral angles across the glycosidic linkage and within the glycan moiety were extracted. These distributions served as input for the Boltzmann inversion to extract the initial CG bonded parameters. Subsequently, CG bonded parameters were iteratively refined by performing short CG simulations and comparing the resulting distributions to the atomistic reference. Minor adjustments to the bonded force constants and equilibrium values were introduced until convergence was achieved.

The final parameter set accurately reproduced the conformational flexibility of the glycosidic linkage and the overall glycan geometry while maintaining full compatibility with the Martini 2.2 force field. The optimized bonded parameters and bead mappings were implemented in the *martinize.py* script.

Flory-Huggins theory

We used the mean-field Flory-Huggins (FH) model[113–115] to estimate the phase-coexistence curves and phase diagram from Martini condensate simulations. The free energy of mixing per residue is given by

$$\frac{\Delta \bar{F}_{mix}}{k_B T} = \frac{\phi}{N} \ln \phi + (1-\phi)\ln(1-\phi) + \chi\phi(1-\phi) \qquad [13]$$

where $\phi$ is the residue volume fraction, $N$ is the number of residues per chain. The Flory parameter $\chi$, which is the interaction energy per residue, is related to the cross interaction energy between protein-solvent $\chi_{ps}$, protein-protein $\chi_{pp}$, and solvent-solvent $\chi_{ss}$ through $\chi = \chi_{ps}\text{-}1/2(\chi_{pp} + \chi_{ss})$. Since in our Martini simulations, the protein-protein interactions are rescaled by parameter $\alpha$, we should expect that the Flory parameter is linearly dependent on $\alpha$, i.e., $\chi = A\alpha + B$. In Eq. 13, the first two terms account for the entropy of mixing chains and solvent, respectively, and the third term accounts for the enthalpy of mixing. Thus, the balance between these terms determines the phase separation criteria. For Flory parameter larger than critical value $\chi_c = 1/2+N^{-1/2}$, the system phase separates into two coexisting dilute and dense phases with volume fractions of $\Phi_\alpha^{\text{dilute}}$ and $\Phi_\alpha^{\text{dense}}$, respectively.

The chemical potential of the system is determined by

$$\frac{\mu}{k_B T} = \frac{1}{k_B T}\frac{\partial \Delta \bar{F}_{mix}}{\partial \phi} = \frac{1}{N}\ln\phi - \ln(1-\phi) + \frac{1}{N} - 1 + \chi(1-2\phi) \qquad [14]$$

In the phase-separated regimes, the chemical potentials of the dilute and dense phases should be equal. Thus, in Eq. 14, by substituting the volume fractions of each phase and one can solve for $\chi$,

$$\chi(\alpha) = \frac{\frac{1}{\mathrm{N}}\log\left(\frac{\Phi_\alpha^{\text{dense}}}{\Phi_\alpha^{\text{dilute}}}\right) + \log\left(\frac{1-\Phi_\alpha^{\text{dilute}}}{1-\Phi_\alpha^{\text{dense}}}\right)}{2\left(\Phi_\alpha^{\text{dense}} - \Phi_\alpha^{\text{dilute}}\right)} \approx A\alpha + B \quad [15]$$

In the Martini condensate simulations at each $\alpha$, the volume fractions of each phase was computed by $\Phi_\alpha^{\text{dilute/dense}} = c_\alpha^{\text{dilute/dense}}/\bar{\rho}$. Here, $c_\alpha^{\text{dilute/dense}}$ is the concentartions of dilute or dense phase $\bar{\rho}$ is the ratio of molar mass and molar volume of an isolated NUP FG chain obtained at $\alpha$ = 0.7 (Supplementary Table 5). Thus, for each set of ($\alpha$, $c_\alpha^{\text{dilute}}$, $c_\alpha^{\text{dense}}$), the value of $\chi$ is determined. The concentrations of the dilute phases was estimated by the double-error function fits to the concentration profiles (Eq. 10 and Supplementary Fig. 13). All $\alpha$-$\chi$ relations are summarized in Supplementary Table 5. The linear relation between $\alpha$ and $\chi$ in Supplementary Fig. 14c validates the linear protein-protein scaling in the α-scaling procedure. The critical protein-protein scaling factor above which the proteins phase separate is given by $\alpha_c = (\chi_c - B)/A$ and they are shown with stars in Supplementary Fig. 14c. The phase-coexistence lines for different NUP FG are shown in Supplementary Fig. 14b. At $\alpha$ = 0.78, NUP54, NUP58C, NUP58N, NUP62 and NUP98 form stable condensates (See final snapshots at $\alpha$ = 0.75 in Supplementary Fig. 13, 14a). The experimental values measured for GLFG homologues of NUP98 FG with dilute and dense phase concentrations are respectively, $6\times10^{-3}$–$6\times10^{-1}$ mg/mL and 314–603 mg/mL[104,116] which is close to the NUP98 FG concentrations at $\alpha$ = 0.78. Additionally, there is a linear relation between $\alpha_c$ and $N^{-1/2}$ with $N$ being the number of residues in each NUP FG domain (Supplementary Fig. 14d). This indicates that the chain-length dependence observed in the Martini condensate simulations is consistent with FH theory.

### Supplementary Figures

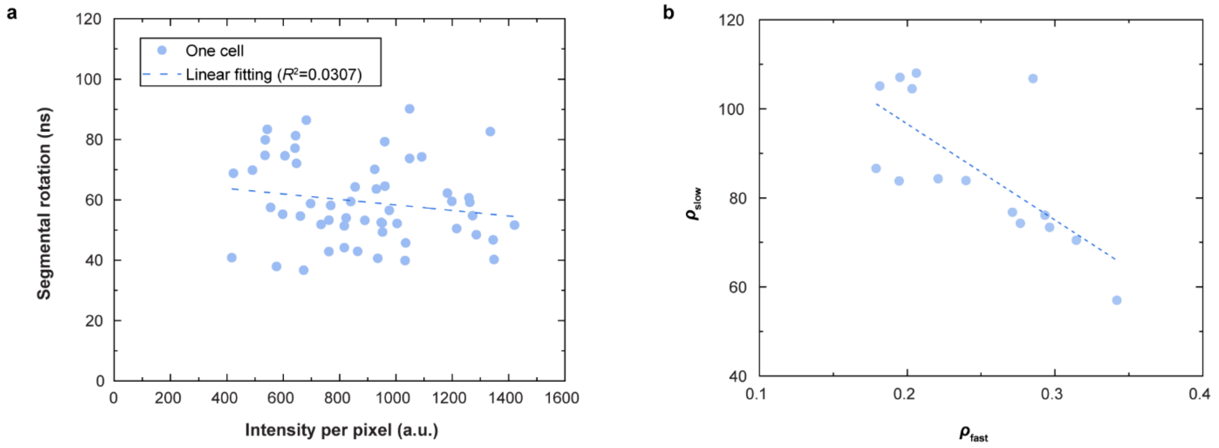


**Supplementary Fig. 1 Expression level does not affect on NUP98 FG segmental dynamics, and local and segmental motions are moderately coupled. a,** Average fluorescence intensity at the nuclear rim (a proxy for NUP98 FG expression level) plotted against segmental rotational correlation time for individual cells. The very low $R^2 = 0.0307$ indicates no correlation, showing that segmental dynamics are insensitive to expression level and are not influenced by labeling artifacts within the tested range. **b**, Relationship between the local rotational correlation time ($\rho_{\text{fast}}$) and segmental rotation correlation time ($\rho_{\text{slow}}$). An expected moderate negative correlation ($R \approx$ -0.5) is observed, consistent with dynamic coupling in intrinsically disordered polymers: slower segmental motion reduces constraints on local dye rotation, whereas faster segmental reorientation partially entrains the dye and slows its local motion.

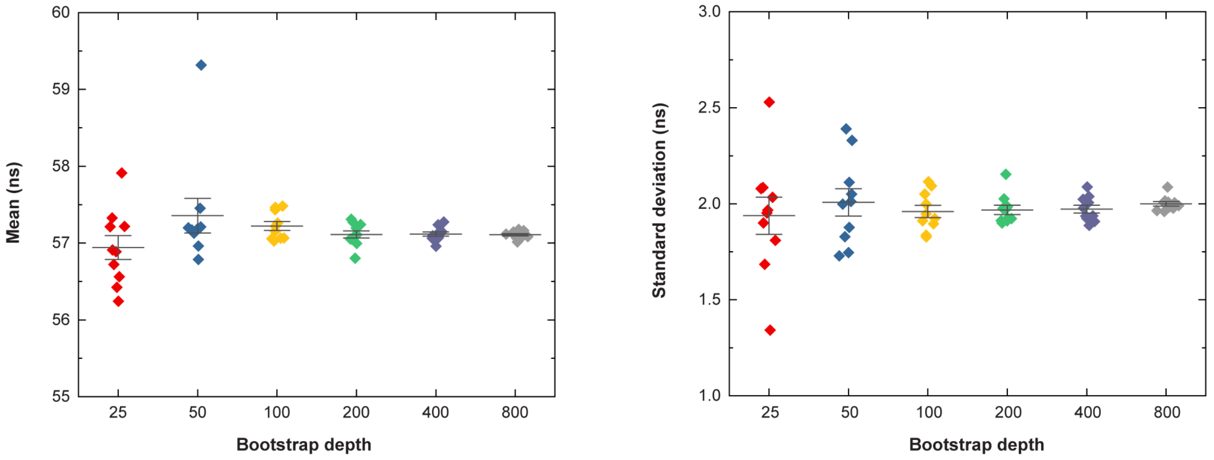


**Supplementary Fig. 2 Convergence analysis for selecting the bootstrap depth used in segmental rotational correlation time fitting.** (Left)  and (right) standard deviation of the fitted segmental rotational correlation time for residue 221 in NUP98 FG domains measured in the NPC, computed across 10 independent trials for each tested bootstrap depth (25, 50, 100, 200, 400, and 800 replicates). At bootstrap depths below 200, both the mean estimates and their variability across trials fluctuate substantially, indicating insufficient resampling. For bootstrap depths ≥200, both metrics stabilize and become nearly indistinguishable from those obtained with larger depths, demonstrating convergence. Based on this analysis, a bootstrap depth of 200 was selected for all experiments.

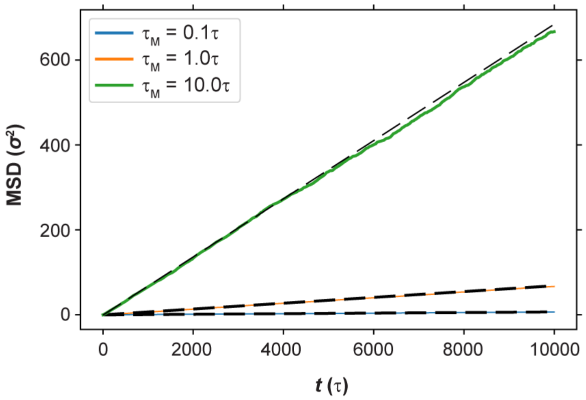


**Supplementary Fig. 3 Mean-squared displacement of a single importin-β in implicit solvent.** Mean-squared displacement (MSD) of a single importin-β molecule in implicit solvent using Langevin thermostats with damping coefficient $\tau_M = 0.1\tau$, $1\tau$, and $10\tau$. The dashed black lines show the theoretical MSD of a Brownian particle with mass $M = 879m$ immersed in a solvent characterized by a friction coefficient $\gamma = M / \tau_M$. Simulation units for mass, time, and distance are $m$, $\tau$, and $\sigma$, respectively.

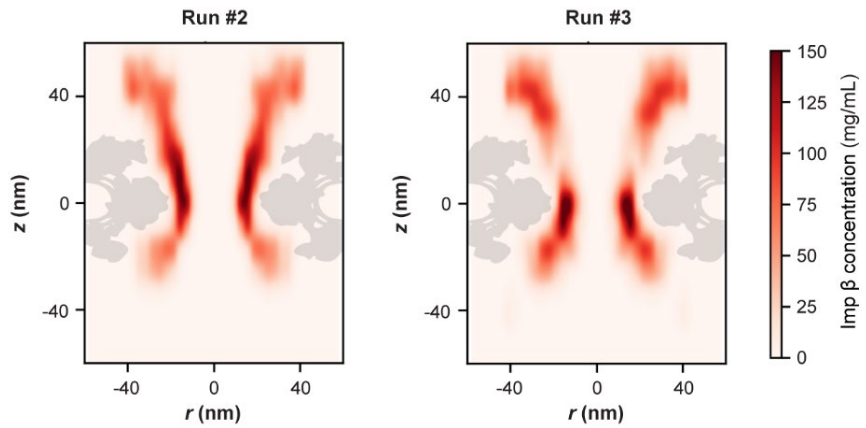


**Supplementary Fig. 4 Concentration maps of importin-β inside the NPC.** Two additional simulation replicates performed under the same conditions as in Fig. 3b (left), using $\tilde{\epsilon}_{\text{FG–Imp-}\beta}$ = 0.42.

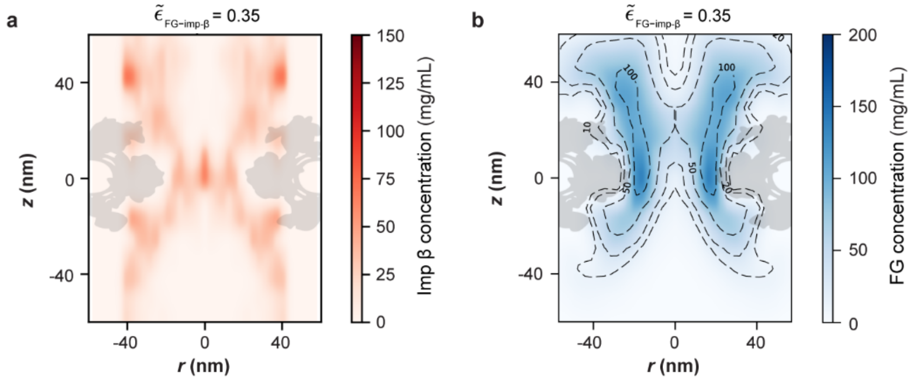


**Supplementary Fig. 5 Importin-β shapes the FG-NUP permeability barrier of the NPC. a,b,** Concentration map of (a) importin-β and (b) FG-NUPs shown in cylindrical coordinates ($r$: radial distance from the symmetry axis; $z$: axial position relative to the nuclear envelope centre with the cytosol at $z > 0$). These simulations use an FG–importin-β interaction strength of $\tilde{\epsilon}_{\text{FG–Imp-}\beta} = 0.35$, and an FG–FG interaction strength of $\tilde{\epsilon}_{\text{FG–FG}} = 0.42$. Maps were averaged over the final $5 \times 10^4\ \tau$, sampled at intervals of $10^2\ \tau$.

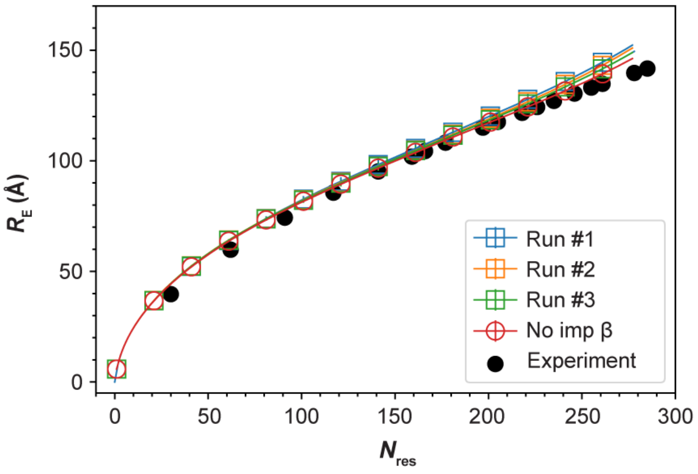


**Supplementary Fig. 6 Root-mean-square inter-residue distance $R_E$ of the NUP98 FG domain.** Root-mean-square inter-residue distance $R_E$ for beads along the NUP98 FG domain in the NPC, plotted as a function of residue separation $N_{res}$. The reference residue ($N_{res} = 0$) is Ala221, and $N_{res}$ is the residue separation toward the C-terminus (the grafting site). The interaction strengths are $\tilde{\epsilon}_{FG\text{–}FG} = 0.42$ for FG–FG interactions and $\tilde{\epsilon}_{FG\text{–}Imp\text{-}\beta} = 0.42$ for FG–importin-β interactions. The symbols and error bars denote the mean and SEM obtained from four non-overlapping blocks of $5 \times 10^4\ \tau$ during the final $2 \times 10^5\ \tau$. The black circles show the corresponding in situ FLIM-FRET measurements of NUP98 chains inside the NPC.[36]

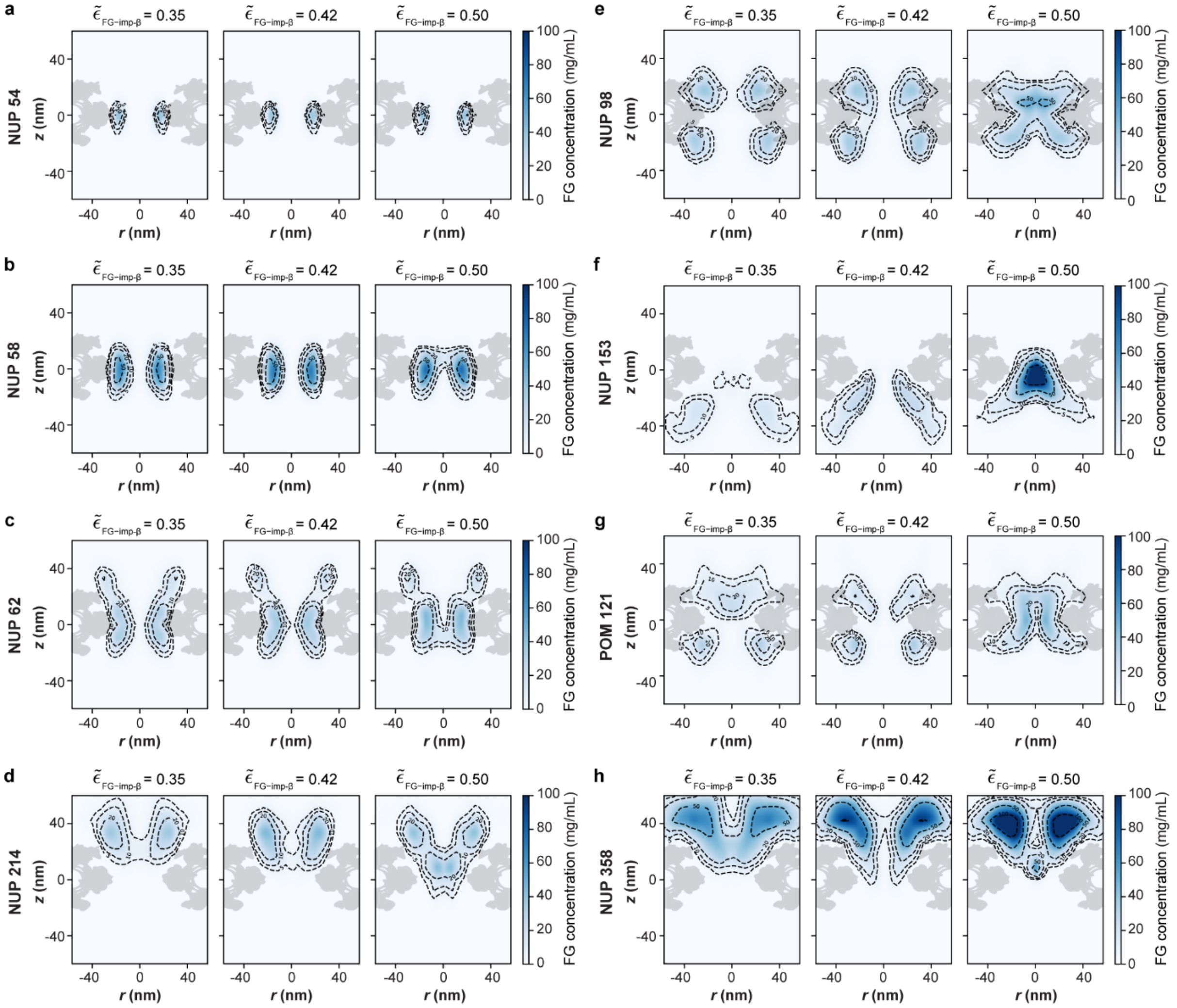


**Supplementary Fig. 7 Concentration maps of individual FG-NUP species inside the NPC in the presence of importin-β. a-h,** Concentration maps of (a) NUP54, (b) NUP58, (c) NUP62, (d) NUP214, (e) NUP98, (f) NUP153, (g) POM121 and (h) NUP358 inside the NPC. For each FG-NUP, the maps are shown for three FG–importin-β interaction strengths: (left) $\tilde{\epsilon}_{\text{FG–Imp-β}} = 0.35$, (middle) 0.42, and (right) 0.5, while the FG–FG interaction strength is fixed at $\tilde{\epsilon}_{\text{FG–FG}} = 0.42$. The concentration maps for all FG-NUPs combined, together with importin-β, are shown in Fig. 3 and Supplementary Fig. 5. Simulation times and the sampling frequency match those used in Fig. 3.

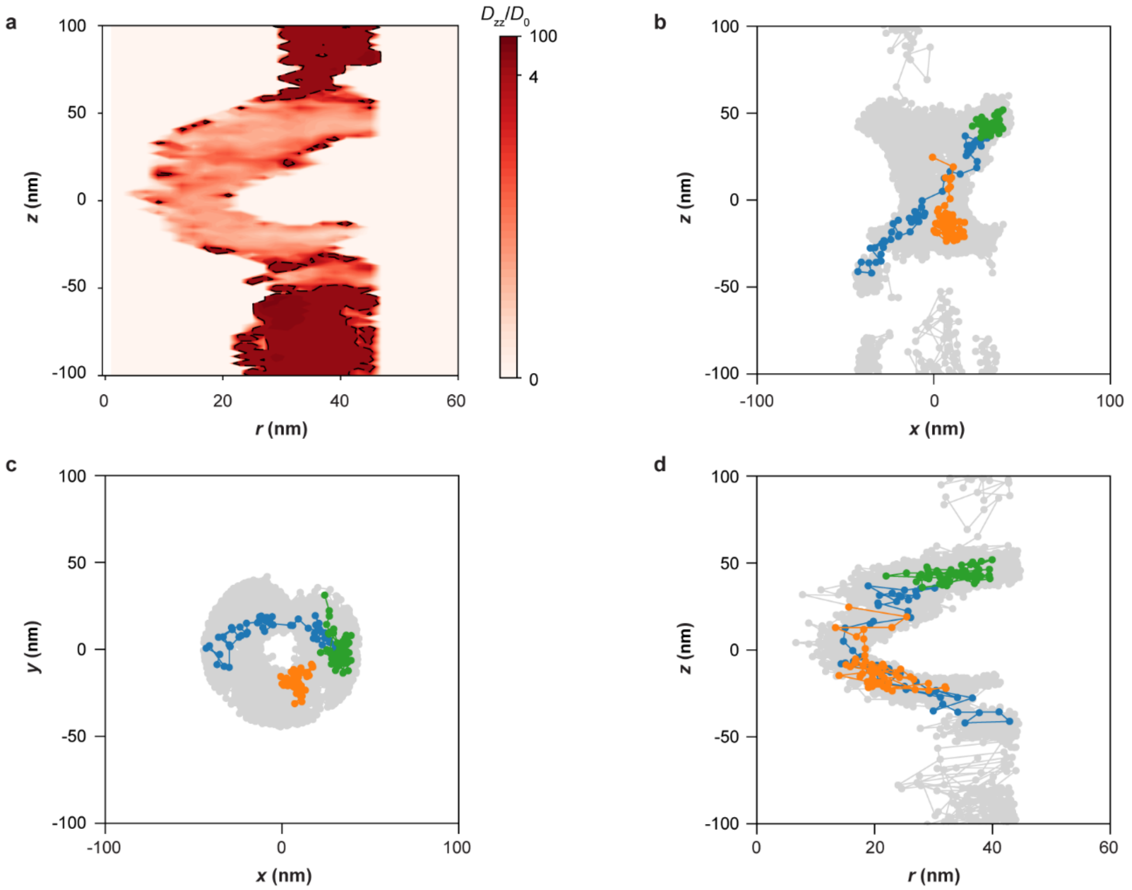


**Supplementary Fig. 8 Dynamic heterogeneity of importin-β molecules inside the NPC. a,** Translational diffusion coefficient map of importin-β molecules along the $z$-axis of the NPC, $D_{zz}$, normalized by $D_0$, the translational diffusion coefficient of importin-β in solution (see Supplementary text). The map was constructed on 2D grid in cylindrical coordinates ($r$, $z$) with a bin size of 2 nm. Importin-β molecules were sorted in bins based on their initial positions; for all molecules in each bin, the squared displacement along $z$-axis, $\langle \Delta z^2 \rangle$, was computed and averaged. Then the translational diffusion coefficient in each bin was obtained as $D_{zz}=\langle \Delta z^2 \rangle/2T$, with $T = 10^4\tau$. Iso-diffusion contours at $D_{zz}/D_0$ =4 are shown with black dashed lines. **b-d,** Representative trajectories of three importin-β molecules projected onto the (b) $x$-$z$, (c) $x$-$y$, and (d) $r$-$z$ planes. The trajectories of all importin-β molecules are shown in grey. The translational diffusion coefficient along the $z$-axis for the three highlighted importin-β molecules was estimated as $\frac{D_{zz}}{D_0} = \frac{\Delta Z^2}{2\Delta t D_0}$ =12.7 (blue), 4.8 (orange) and 0.06 (green). The total simulation time for the trajectories is $\Delta t = 4.96 \times 10^5\tau$. All simulations used interaction strengths $\tilde{\epsilon}_{\text{FG–FG}} = \tilde{\epsilon}_{\text{FG–Imp-β}} = 0.42$.

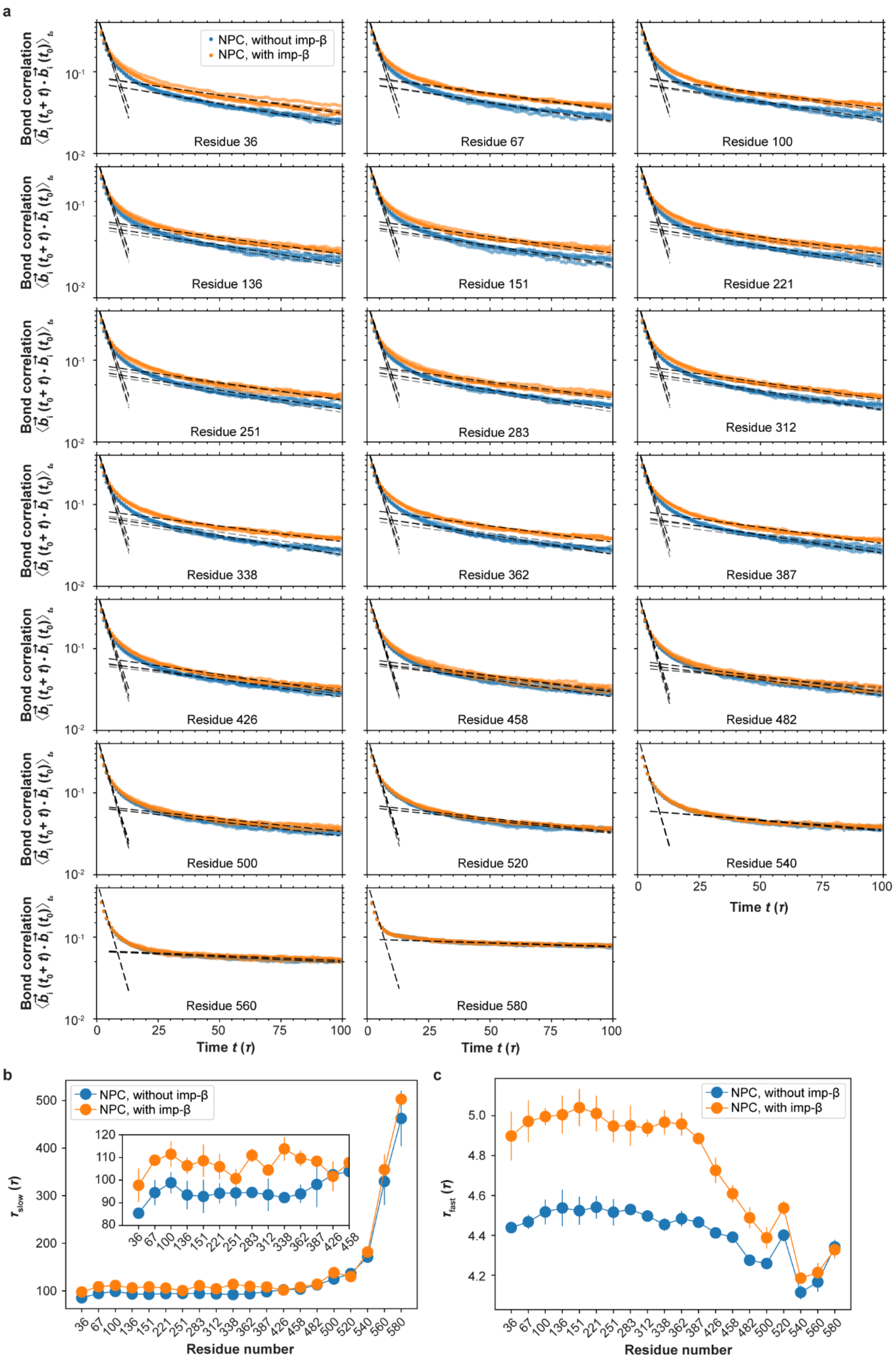

a
NPC, without imp-β
NPC, with imp-β
Bond correlation
$\langle \vec{b}_i(t_0+t) \cdot \vec{b}_i(t_0) \rangle_{t_0}$
$10^{-1}$
$10^{-2}$
Residue 36
Residue 67
Residue 100
Residue 136
Residue 151
Residue 221
Residue 251
Residue 283
Residue 312
Residue 338
Residue 362
Residue 387
Residue 426
Residue 458
Residue 482
Residue 500
Residue 520
Residue 540
Residue 560
Residue 580
0
25
50
75
100
Time $t$ ($\tau$)
b
NPC, without imp-β
NPC, with imp-β
$\tau_{slow}$ ($\tau$)
100
200
300
400
500
80
90
100
110
120
36 67 100 136 151 221 251 283 312 338 362 387 426 458
36 67 100 136 151 221 251 283 312 338 362 387 426 458 482 500 520 540 560 580
Residue number
c
NPC, without imp-β
NPC, with imp-β
$\tau_{fast}$ ($\tau$)
4.2
4.4
4.6
4.8
5.0
36 67 100 136 151 221 251 283 312 338 362 387 426 458 482 500 520 540 560 580
Residue number

**Supplementary Fig. 9 Segmental rotations of the NUP98 FG domain in the NPC with and without importin-β. a,** Auto-correlation function of the bond vector $\langle \overrightarrow{b_i}(t_0 + t) \cdot \overrightarrow{b_i}(t_0) \rangle_{t_0}$ for residues 36, 67, 100, 136, 151, 221, 251, 283, 312, 338, 362, 387, 426, 458, 482, 500, 520, 540, 560 and 580 of the NUP98 FG domain. Here, $\overrightarrow{b_i}$ is the unit bond vector connecting residue *i* to *i*+1. The black dashed lines represent single-exponential fits to the averaged autocorrelation function $a\exp(-t/\tau_{\text{fast}})$ and $a'\exp(-t/\tau_{\text{slow}})$ over the short-time window [0, $10\tau$] and the long-time window [$25\tau$, $100\tau$], respectively, as a coarse two-timescale approximation to inherently multiexponential dynamics. The data points show the results from four independent simulation runs. The total simulation times were $10^5\tau$ for the single chain and $10^4\tau$ for the condensate and the NPC simulations, with the coordinates of each selected residue sampled every $\tau$. All simulations used interaction strengths $\tilde{\epsilon}_{\text{FG–FG}} = \tilde{\epsilon}_{\text{FG–Imp-β}} = 0.42$. **b,c** Slow ($\tau_{\text{slow}}$, b) and fast ($\tau_{\text{fast}}$, c) relaxation times extracted from the fits in (a). Symbols and error bars denote the mean and SEM across the four independent simulation runs.

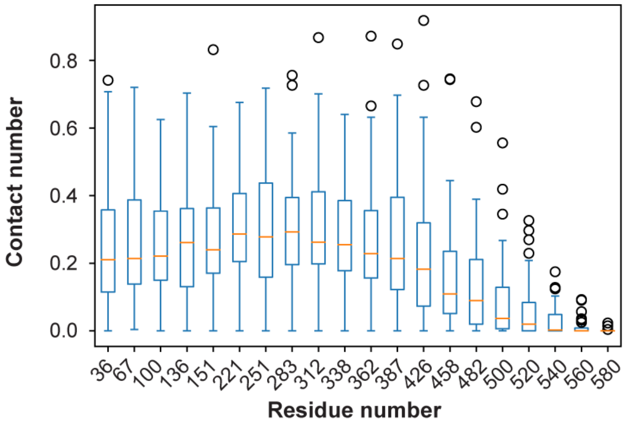


**Supplementary Fig. 10 Average number of contacts between NUP98 FG domain and importin-β inside the NPC.** The box-whisker plot shows the distribution of the mean number of contacts between residues of the NUP98 FG domain and importin-β molecules. The orange line indicates the median, the blue box the interquartile range, the whiskers the full data range, and circles denote outliers. Each box summarizes the mean contact number computed for 48 NUP98 FG chains inside the NPC, averaged over the final $5 \times 10^4\tau$ of the simulation.

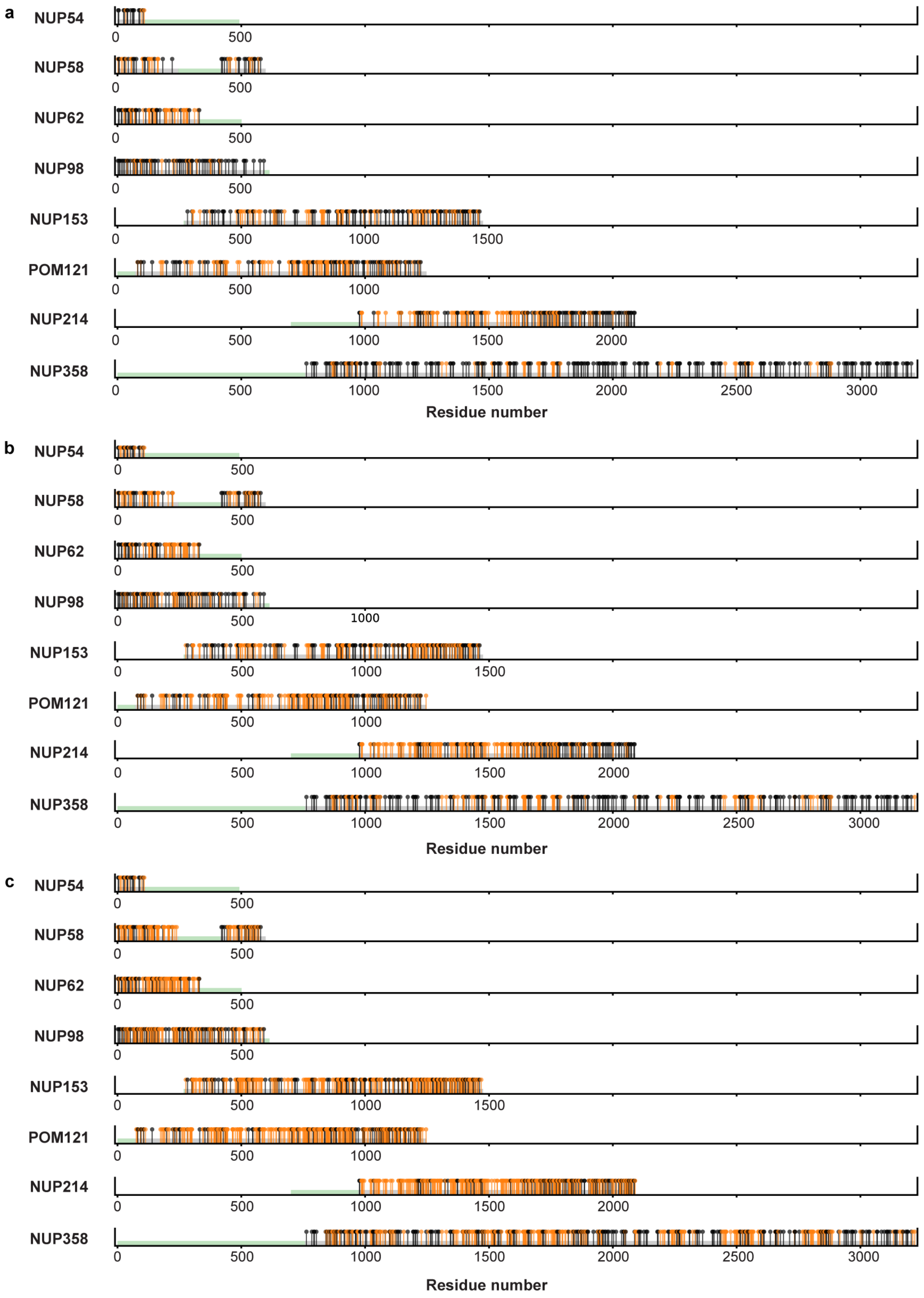


**Supplementary Fig. 11 O-Glycosylation sites along FG-NUPs inside the NPC. a-c,** The position of O-glycosylated Ser/Thr (orange pins) and hydrophobic residues with aromatic rings (black pins; Phe/Trp/Tyr) along FG-NUP sequences for three levels of O-glycosylation: (a) $f_{gly}$ =

0.18, (b) $f_{gly}$ = 0.3 and (c) $f_{gly}$ = 0.6. The green and grey strips represent the resolved ordered and disordered domains, respectively, along each FG-NUP chain.

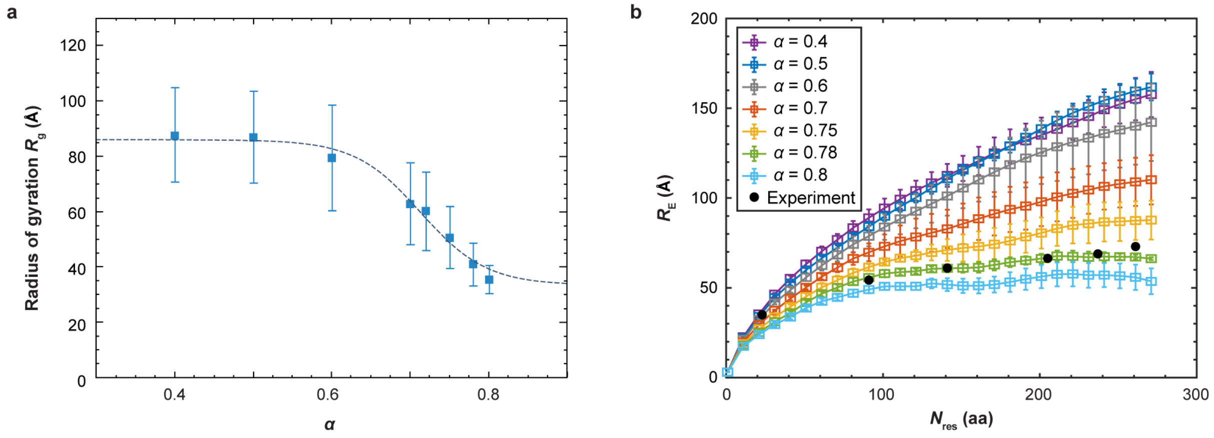


**Supplementary Fig. 12 Radius of gyration and root-mean-squared inter-residue distance of NUP98 FG domain in the Maritni force field. a,** Radius of gyration $R_g$ of a single NUP98 FG chain as a function of the interaction rescaling parameter $\alpha$. The solid line shows a logistic function fit to the form $R_g(\alpha) = a+b/(1+\exp[-c(\alpha-\alpha_{cg})])$ with the values $a$ = 33.41, $b$ = 52.59, $c$ = -24.19 and $\alpha_{cg}$ = 0.71. Symbols and error bars represent the mean and standard deviation of $R_g$ over the final 8 μs of the trajectory. **b**, Root-mean-squared inter-residue distance $R_E$ of a single NUP98 FG for different values of $\alpha$. The value at $\alpha$ = 0.78 reproduces single-molecule FRET measurements.[36] The symbols and error bars denote the mean and standard error of the mean computed over four non-overlapping 2 μs blocks.

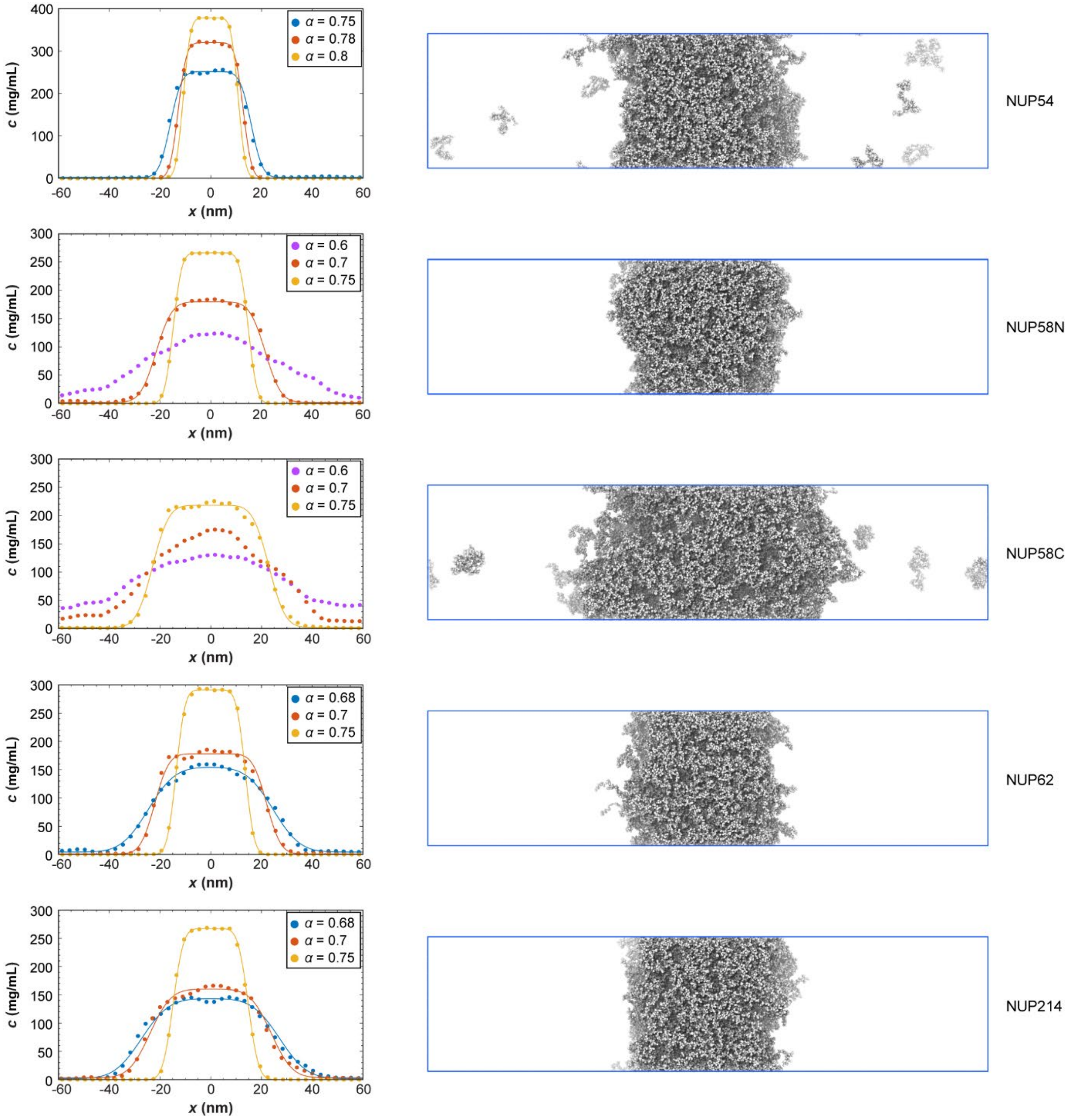


**Supplementary Fig. 13 Concentrations of FG-NUPs condensates at different $\alpha$ scaling values.** Concentration profiles (left column) of condensates formed by FG-NUP54, FG-NUP58N, FG-NUP58C, FG-NUP62, FG-NUP214 at different interaction rescaling parameters $\alpha$. The solid lines indicate fits to a double error function model. Representative simulation snapshots (right column) correspond to $\alpha$ = 0.75. All systems have dimensions $L_x \approx 120$ nm and $L_y = L_z = 30$ nm, and simulation details are provided in Supplementary Table 4.

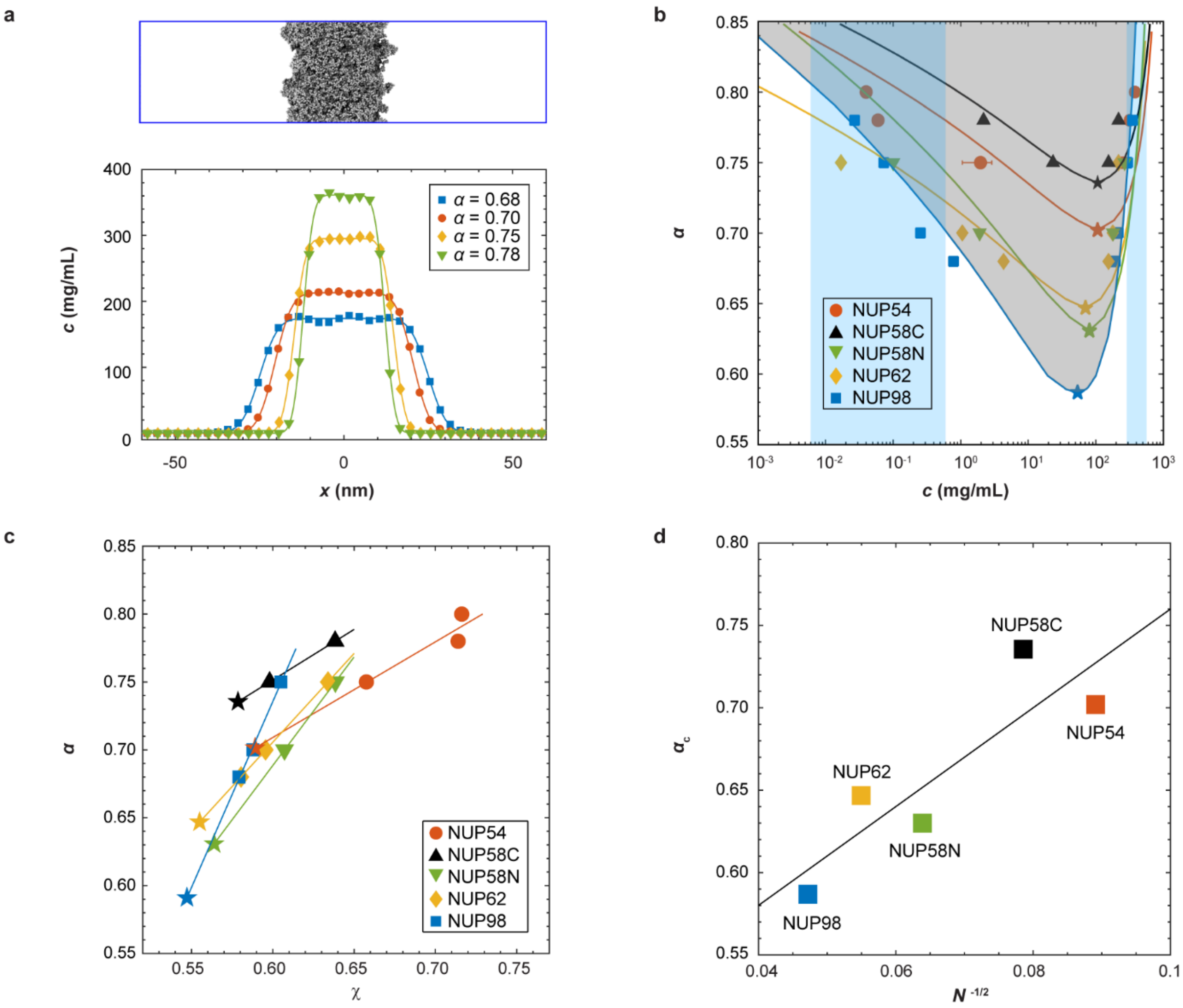


**Supplementary Fig. 14 Concentration profiles and phase behaviour of FG-NUPs. a**, (Top) Snapshot of a system composed of 100 FG-NUP98 chains at $\alpha$ = 0.78. The chains are shown in white; solvent particles and ions are omitted for clarity. (Bottom) The concentration profiles of FG-NUP98 at different protein–protein interaction scaling parameters $\alpha$. The solid lines represent the double error function fits (See Supplementary text). **b**, Phase diagram of the interaction scaling parameter and concentration ($\alpha$-$c$) for different FG-NUPs. The solid lines represent the coexistence curves predicted by Flory-Huggins theory (See Supplementary text). The critical interaction scaling parameter and concentration ($\alpha_c$, $c_c$) for each FG-NUP are indicated by color-matched symbols. Transparent vertical blue bands denote the experimentally measured dilute-phase and dense-phase concentrations for GLFG-homology regions of NUP98 FG domains ($6\times10^{-3}$–$6\times10^{-1}$ mg/mL and 314–603 mg/mL)[104,116]. **c**, Linear relationship between $\alpha$ and the Flory interaction parameter $\chi$ for different FG-NUPs (See Supplementary Text). The data points were obtained by enforcing equality of the chemical potential between dilute and dense phases. **d**, Scaling of the critical protein-protein scaling factor $\alpha_c$ versus $N^{-1/2}$, with $N$ being the number of residues in each FG-NUP. The solid line shows a linear fit, $\alpha_c = AN^{-1/2} + B$, with fitted parameters $A$=3 and $B$=0.46.

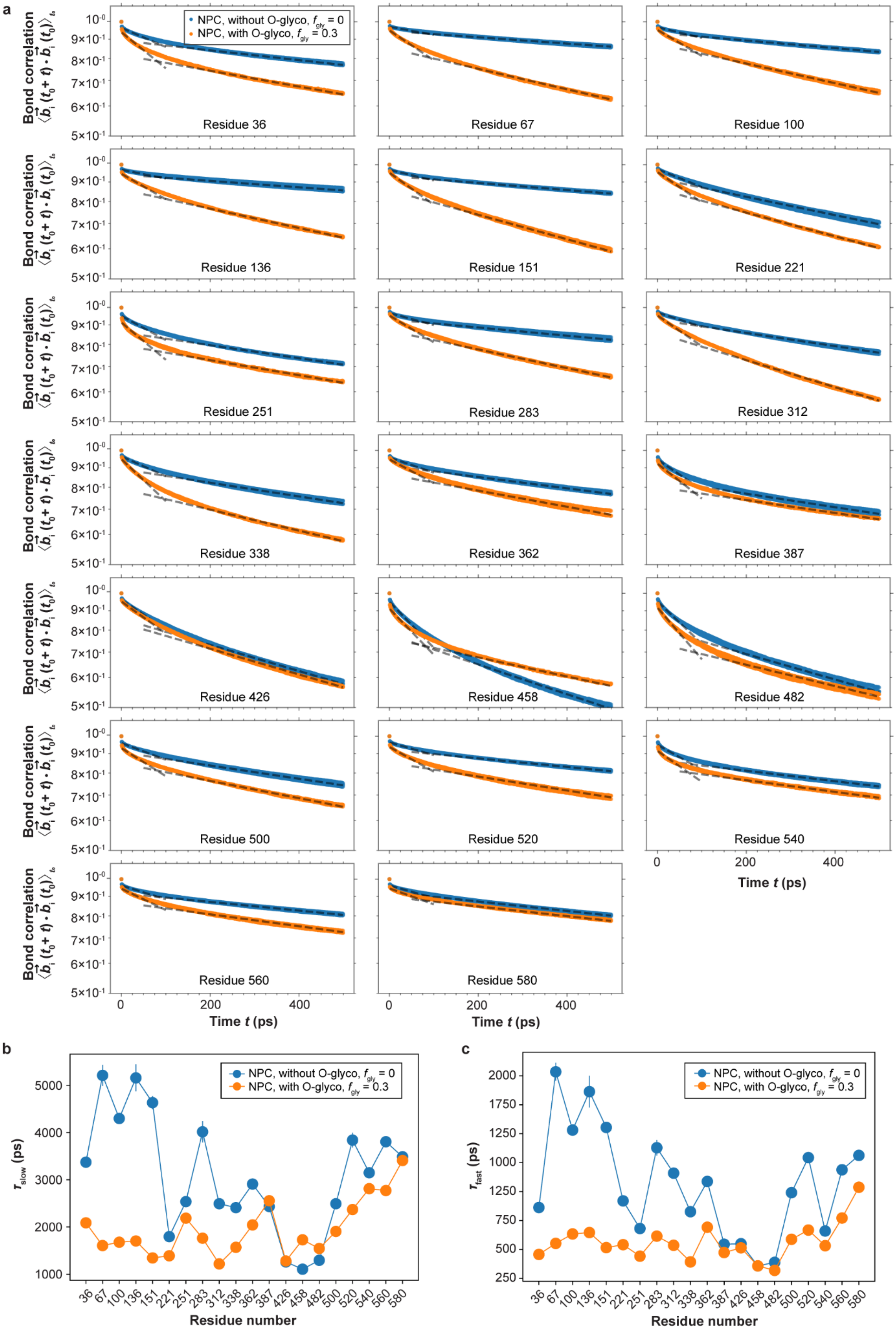
a
NPC, without O-glyco, $f_{gly}$ = 0
NPC, with O-glyco, $f_{gly}$ = 0.3
Bond correlation
Residue 36
Residue 67
Residue 100
Residue 136
Residue 151
Residue 221
Residue 251
Residue 283
Residue 312
Residue 338
Residue 362
Residue 387
Residue 426
Residue 458
Residue 482
Residue 500
Residue 520
Residue 540
Residue 560
Residue 580
Time t (ps)
b
$\tau_{slow}$ (ps)
Residue number
c
$\tau_{fast}$ (ps)
Residue number

**Supplementary Fig. 15 Segmental rotations of the NUP98 FG domain in the Martini model of the NPC with and without O-glycosylation. a,** Auto-correlation function of the bond vector $\langle \vec{b_i}(t_0+t)\cdot\vec{b_i}(t_0)\rangle_{t_0}$ for residues 36, 67, 100, 136, 151, 221, 251, 283, 312, 338, 362, 387, 426, 458, 482, 500, 520, 540, 560 and 580 of the NUP98 FG domain. $\vec{b_i}$ is the unit bond vector connecting backbone beads of residue *i* to *i*+1. The black dashed lines represent single-exponential fits to the averaged autocorrelation functions $a\exp(-t/\tau_{\mathrm{fast}})$ and $a'\exp(-t/\tau_{\mathrm{slow}})$ over the short-time window [0, 10$\tau$] and the long-time window [25$\tau$, 100$\tau$], respectively, as a coarse two-timescale approximation to inherently multiexponential dynamics. The data points show the results from three independent simulation runs, each with a total duration of 100 ns. **b,c** Slow ($\tau_{\mathrm{slow}}$, b) and fast ($\tau_{\mathrm{fast}}$, c) relaxation times extracted from the fits in (a). Symbols and error bars denote the mean and SEM across the three independent simulation runs.

### Supplementary Tables

**Supplementary Table 1. Plasmids used in this work**

| **Construct** | **In Figure** |
|---|---|
| pcDNA3.1-FLAG-hsNUP98$^{A221TAG}$-boxB | Fig.1,4, Extended data Fig.1,8 |
| pcDNA3.1-FLAG-hsNUP98$^{A36TAG}$-boxB | Fig.1, Extended data Fig.1,8 |
| pcDNA3.1-FLAG-hsNUP98$^{S67TAG}$-boxB | Fig.1, Extended data Fig.1,8 |
| pcDNA3.1-FLAG-hsNUP98$^{S100TAG}$-boxB | Fig.1,4, Extended data Fig.1,8 |
| pcDNA3.1-FLAG-hsNUP98$^{T136TAG}$-boxB | Fig.1, Extended data Fig.1,8 |
| pcDNA3.1-FLAG-hsNUP98$^{S151TAG}$-boxB | Fig.1, Extended data Fig.1,8 |
| pcDNA3.1-FLAG-hsNUP98$^{A251TAG}$-boxB | Fig.1,4, Extended data Fig.1,8 |
| pcDNA3.1-FLAG-hsNUP98$^{S283TAG}$-boxB | Fig.1,4, Extended data Fig.1,8 |
| pcDNA3.1-FLAG-hsNUP98$^{S312TAG}$-boxB | Fig.1,4, Extended data Fig.1,8 |
| pcDNA3.1-FLAG-hsNUP98$^{S338TAG}$-boxB | Fig.1,4, Extended data Fig.1,8 |
| pcDNA3.1-FLAG-hsNUP98$^{S362TAG}$-boxB | Fig.1,4, Extended data Fig.1,8 |
| pcDNA3.1-FLAG-hsNUP98$^{S387TAG}$-boxB | Fig.1,4, Extended data Fig.1,8 |
| pcDNA3.1-FLAG-hsNUP98$^{A426TAG}$-boxB | Fig.1,4, Extended data Fig.1,8 |
| pcDNA3.1-FLAG-hsNUP98$^{458TAG}$-boxB | Fig.1,4, Extended data Fig.1,8 |
| pcDNA3.1-FLAG-hsNUP98$^{A482TAG}$-boxB | Fig.1,4, Extended data Fig.1,8 |
| pcDNA3.1-TOM20$_{1\text{-}70}$-FUS$_{1\text{-}478}$-4x$\lambda_{N22}$-PylRS$^{Y306A,Y384F}$-U6-tRNA$^{Pyl}$ | Fig.1,4, Extended data Fig.1,8 |
| pQE-14His-TEV-hsNUP98$_{FG\ 1\text{-}505,\ \Delta GLEBS}$ | Fig.1, 2 and 4, Extended data Fig.2,8 |
| pQE-14His-TEV-hsNUP98$_{FG\ 1\text{-}505,\ \Delta GLEBS}{}^{A221C}$ | Fig. 1, 2 and 4, Extended data Fig.2,8 |
| pQE-14His-TEV-hsNUP98$_{FG\ 1\text{-}505,\ \Delta GLEBS}{}^{A36C}$ | Fig. 2,4, Extended data Fig.2,8 |
| pQE-14His-TEV-hsNUP98$_{FG\ 1\text{-}505,\ \Delta GLEBS}{}^{S100C}$ | Fig. 2,4, Extended data Fig.2,8 |
| pQE-14His-TEV-hsNUP98$_{FG\ 1\text{-}505,\ \Delta GLEBS}{}^{S312C}$ | Fig. 2,4, Extended data Fig.2,8 |
| pQE-14His-TEV-hsNUP98$_{FG\ 1\text{-}505,\ \Delta GLEBS}{}^{S387C}$ | Fig. 2,4, Extended data Fig.2,8 |
| pQE-14His-TEV-hsNUP98$_{FG\ 1\text{-}505,\ \Delta GLEBS}{}^{A482C}$ | Fig. 2,4, Extended data Fig.2,8 |

**Supplementary Table 2. Interaction parameters used in coarse-grained MD simulations of importin-β and the NPC.** Lennard-Jones (LJ) interaction parameters for all bead-type pairs used in the coarse-grained simulations. For beads that are fixed in space or that belong to the same rigid body (including scaffold residues, membrane particles, and importin-β beads), cross-interaction terms were omitted from the energy function and therefore are not listed.

| Type | $j \in$ sc | $j \in$ FG | $j \in m$ | $j \in$ Imp-β |
|---|---|---|---|---|
| $i \in$ sc | - | $\sigma_{ij} = \sigma$, $r_c = 2\sigma$, $\tilde{\epsilon}_{ij} = 0.1$ | - | $\sigma_{ij} = \sigma$, $r_c = 2\sigma$, $\tilde{\epsilon}_{ij} = 0.1$ |
| $i \in$ FG | Symmetric | $\sigma_{ij} = \sigma$, $r_c = 2\sigma$, $\tilde{\epsilon}_{ij} = \tilde{\epsilon}_{\text{FG-FG}}$ | $\sigma_{ij} = 1.78\sigma$, $r_c = 1.99\sigma$, $\tilde{\epsilon}_{ij} = 0.1$ | $\sigma_{ij} = \sigma$, $r_c = 2\sigma$, $\tilde{\epsilon}_{ij} = \tilde{\epsilon}_{\text{FG-Imp}\,\beta}$ |
| $i \in m$ | - | Symmetric | - | $\sigma_{ij} = 1.78\sigma$, $r_c = 1.99\sigma$, $\tilde{\epsilon}_{ij} = 0.1$ |
| $i \in$ Imp-β | Symmetric | Symmetric | Symmetric | $\sigma_{ij} = \sigma$, $r_c = 2\sigma$, $\tilde{\epsilon}_{ij} = 0.1$ |

**Supplementary Table 3. Amino acid sequence of FG-NUPs used in the MD simulations of the NPC.**

| Chain | Number of copies | Beginning/ ending residues | Chain name in PDB file | Grafting residues of dynamic region(s) | Restrained residue range |
|---|---|---|---|---|---|
| NUP54 | 32 | (2-493) | chains H | THR112 | 112-493 |
| NUP58 | 32 | (2-599) | chains I | LEU247 & TYR416 | 247-416 |
| NUP62 | 48 | (2-502) | chains J | GLN332 | 332-502 |
| NUP98 | 48 | (2-615) | chains U | SER596 | 596-615 |
| NUP98 | 8 | (731-880) | chains UCo | - | 731-880 |
| NUP214 | 8 | (954-2090) | chains VCi | ARG954 | 954 |
| NUP214 | 8 | (700-2090) | chains VCo | LEU973 | 700-973 |
| NUP153 | 16 | (272-1475) | chains Y | ALA272 | 272 |
| POM121 | 16 | (2-1249) | chains 2 | GLU72 | 2-72 |
| NUP358* | 40 | (4-3224) | chains 0 | SER833 | 4-833 |

*The sequence range of local ordered domains of NUP358 is as follows: 1190-1306, 1354-1379, 1417-1442, 1481-1506, 1546-1572, 1608-1633, 1669-1692, 1727-1751, 1783-1808, 2031-2148, 2328-2446, 2929-3050, 3054-3224.

**Supplementary Table 4. FG condensate systems and corresponding simulation parameters.**

| Condensate | Number of chains | Residue range | Simulation box $L_x \times L_y \times L_z$ (nm$^3$) | Molecular weight of single chain (g/mol) | Simulation time (μs) |
|---|---|---|---|---|---|
| NUP54 | 400 | (2-127) | 125×30×30 | 11124 | ≈20 |
| NUP58N | 350 | (438-599) | 125×30×30 | 21412 | ≈20 |
| NUP58C | 200 | (2-246) | 125×30×30 | 15764 | ≈20 |
| NUP62 | 150 | (2-332) | 125×30×30 | 28707 | ≈20 |
| NUP98 | 100 | (1-499) | 125×30×30 | 49158 | ≈20 |
| NUP214 | 150 | (1786-2090) | 125×30×30 | 28631 | ≈20 |

**Supplementary Table 5. $\alpha$-$\chi$ relation for FG-NUPs.** The fitting parameters used to map the Flory–Huggins $\chi$-$\phi$ phase diagram onto the interaction–concentration ($\alpha$-$c$) representation. These $\alpha$-$\chi$ relations were obtained by fitting the coexistence data for each FG-NUP species.

| Chain | $N$ | $\alpha - \chi$ | $\bar{\rho}$ (mg/mL) |
|---|---|---|---|
| NUP54 | 126 | $\alpha = 0.7046\chi + 0.2863$ | 1338 |
| NUP58C | 162 | $\alpha = 0.7452\chi + 0.3044$ | 1469 |
| NUP58N | 245 | $\alpha = 1.5499\chi - 0.2707$ | 1338 |
| NUP62 | 331 | $\alpha = 1.307\chi - 0.0785$ | 1356 |
| NUP98 | 499 | $\alpha = 2.738\chi - 0.9074$ | 1263 |

**Supplementary Table 6. Bond-orientation relaxation times of the Ala221 residue in the NUP98 FG domain.** Relaxation times for the Ala221 bond vector in a single FG chain, an FG condensate, and the NPC. Uncertainties represent the standard deviation of the relaxation times across three independent replicas.

| System | $\tau_{\mathbf{fast}}$**(ps)** | $\tau_{\mathbf{slow}}$**(ps)** |
|---|---|---|
| Single chain | 412 ± 8 | 1267 ± 5 |
| Condensate | 679 ± 15 | 2039 ± 6 |
| NPC ($f_{gly}$ = 0) | 919 ± 13 | 1800 ± 4 |
| NPC ($f_{gly}$ = 0.3) | 541 ± 10 | 1390 ± 3 |

### Movie captions

**Supplementary Movie 1:** Side-by-side comparison of (co-)phase separation under four conditions: (1) wild-type NUP98 FG domains alone, (2) wild-type NUP98 FG domains with importin-β, (3) O-GlcNAcylated NUP98 FG domains alone, and (4) O-GlcNAcylated NUP98 FG domains with importin-β.

**Supplementary Movie 2:** Simulation trajectories of NUP98 FG and importin-β mixture during the final $4.2 \times 10^3\tau$ at different molecular ratios. From top to bottom, the left column corresponds to $N_{\text{Imp-}\beta}/N_{\text{NUP98}} = 0.2$, 0.4 and 0.6, and the right column to $N_{\text{Imp-}\beta}/N_{\text{NUP98}} = 0.8$, 1 and 2. The interaction strengths are $\tilde{\epsilon}_{\text{FG}-\text{FG}} = 0.5$ between FG-FG interactions and $\tilde{\epsilon}_{\text{FG–Imp-}\beta} = 0.42$ for FG–importin-β interactions.

**Supplementary Movie 3:** Simulation trajectories of the NPC containing 100 importin-β molecules during the final $5 \times 10^4\tau$. The top panel shows the cytoplasmic-side view; the bottom panel shows a cross-sectional side view. Interaction strengths are $\tilde{\epsilon}_{\text{FG–FG}} = 0.42$ and $\tilde{\epsilon}_{\text{FG–Imp-}\beta} = 0.42$.

**Supplementary Movie 4**: Simulation trajectories of the NPC containing 100 importin-β molecules during the final $5 \times 10^4\tau$. The top panel shows the cytoplasmic-side view; the bottom panel shows a cross-sectional side view. Interaction strengths are $\tilde{\epsilon}_{\text{FG–FG}} = 0.42$ and $\tilde{\epsilon}_{\text{FG–Imp-}\beta} = 0.50$.

**Supplementary Movie 5**: Simulation trajectories of the NPC containing 100 importin-β molecules during the final $5 \times 10^4\tau$. The top panel shows the cytoplasmic-side view; the bottom panel shows a cross-sectional side view. Interaction strengths are $\tilde{\epsilon}_{\text{FG–FG}} = 0.42$ and $\tilde{\epsilon}_{\text{FG–Imp-}\beta} = 0.35$.

**Supplementary Movie 6**: Simulation trajectories of untreated NPC ($f_{\text{gly}} = 0$) during the final 500 ns. The interaction scaling parameters are $\alpha = 0.78$, $\gamma = 0.5$ and $\lambda = 0.15$. The FG-NUPs are shown in white and the scaffold proteins in green.

**Supplementary Movie 7**: Simulation trajectories of partially O-glycosylated NPC ($f_{\text{gly}} = 0.18$) during the final 500 ns. The interaction scaling parameters are $\alpha = 0.78$, $\gamma = 0.5$ and $\lambda = 0.15$. Unglycosylated FG-NUPs are shown in white; O-glycosylated FG-NUPs in yellow; scaffold proteins in green.

**Supplementary Movie 8**: Simulation trajectories of partially O-glycosylated NPC ($f_{\text{gly}} = 0.3$) during the final 500 ns. The interaction scaling parameters are $\alpha = 0.78$, $\gamma = 0.5$ and $\lambda = 0.15$. Unglycosylated FG-NUPs are shown in white; O-glycosylated FG-NUPs in yellow; scaffold proteins in green.

**Supplementary Movie 9**: Simulation trajectories of partially O-glycosylated NPC ($f_{\text{gly}} = 0.6$) during the final 500 ns. The interaction scaling parameters are $\alpha = 0.78$, $\gamma = 0.5$ and $\lambda = 0.15$.

Unglycosylated FG-NUPs are shown in white; O-glycosylated FG-NUPs in yellow; scaffold proteins in green.
